# Unlocking Multi-Sample Differential Expression for Spatial Transcriptomics Data with TESSERA

**DOI:** 10.64898/2026.04.27.720955

**Authors:** Florica Constantine, Zoltan Laszik, Sandrine Dudoit, Elizabeth Purdom

**Author notes:** These authors contributed equally to this work.

## Abstract

Spatial transcriptomics allows the unprecedented examination of gene expression levels at the resolution of spatially-situated single cells in a high-throughput manner. As the technology is adopted more broadly, studies frequently collect data from multiple tissue samples, which leads to unique challenges that traditional spatial statistical methods are not equipped to handle. In particular, factors that differ across samples, such as different coordinate systems, different numbers and types of cells, different underlying tissue architectures, among others, preclude the application of traditional methods to our new setting. In this work, we propose a novel method, `TESSERA`, based on a spatial generalized linear model, for analyzing multi-sample spatial transcriptomics count data. Importantly, we provide a mathematical and computational framework for efficient and scalable model fitting and statistical inference to accompany the specification of our model. Our method for fitting the model enables the estimation of a common set of fixed effects across samples. This allows us to address a variety of differential expression questions, such as identification of which genes are differentially expressed between conditions (e.g., diseases, treatments), while accounting for spatial correlation between cells within a sample. We benchmark our proposed method on simulated data and apply it to a spatial transcriptomics dataset of human kidney samples. We find that our method provides a hitherto nonexistent extension to the multi-sample setting while remaining competitive with or outperforming existing algorithms in the single-sample setting.

## 1. Introduction

### 1.1. Biological Background

Spatial transcriptomics allows for the extraction of mRNA expression measurements at cellular or near-cellular resolution while simultaneously recording the physical coordinates for cell locations within a tissue (Williams et al., 2022; Ståhl et al., 2016; Rodriques et al., 2019; Vickovic et al., 2019). Spatial transcriptomics datasets typically contain hundreds to tens of thousands of cells or “spots” (i.e., groups of approximately 1 to 10 cells (10x Genomics, 2025a)). The data are highly sparse, with the majority of gene–cell or gene–spot entries equal to zero. This sparsity arises from a combination of true biological absence of expression and technical dropout, where only a fraction of the total transcripts present are successfully captured during data collection (Moses and Pachter, 2022; Williams et al., 2022; Zhao et al., 2022). Prior to the introduction of spatial transcriptomics technology, practitioners had been using single-cell RNA sequencing (scRNA-seq) to obtain gene expression measurements at the resolution of individual cells. However, scRNA-seq cannot be applied *in situ* as the tissue is dissociated to allow for the extraction of mRNA from each cell, which results in the loss of spatial information regarding cell locations within the tissue. By contrast, spatial transcriptomics technology operates at a cellular or near-cellular resolution by retaining a cell’s or spot’s spatial location in a tissue. There are two main categories of technologies, next-generation sequencing and imaging. Next-generation sequencing technologies capture mRNA molecules using barcodes along a grid of known spatial locations (Vickovic et al., 2019; Ståhl et al., 2016; Rodriques et al., 2019), while imaging-based methods detect mRNA molecules through the generation of fluorescence images from the barcoding of fluorescent probes via *in situ* sequencing or hybridization (Rao et al., 2021; Lee et al., 2014; Chen et al., 2015). The retention of the spatial positioning of the cells in the tissue enables identification of neighborhoods/neighboring cells, which allows us to better study cell behavior, as a cell’s function and associated cell-cell interactions are often highly location-dependent (Dries et al., 2021; Pareja-Lorente and Aloy, 2025; Joost et al., 2016; Hunter et al., 2021).

### 1.2. Motivation: Multiple Samples and Differential Expression

Researchers will often use mRNA expression measurements to identify differentially expressed genes between populations (e.g., patients with different responses to treatment, wild-type and mutant mice) in order to better understand biological processes, disease mechanisms, or the effects of treatments. A gene that exhibits distributional differences in mRNA abundance measures between groups is referred to as differentially expressed (DE). In a typical setting for bulk RNA-seq or scRNA-seq, we seek to identify differential expression for individual genes between two different cell types or states (scRNA-seq) or phenotypes (bulk RNA-seq and scRNA-seq) and can draw on a range of well-established methods that generally involve testing for differences in means (Love, Huber and Anders, 2014; Ritchie et al., 2015; Robinson, McCarthy and Smyth, 2010; Soneson and Robinson, 2018; Finak et al., 2015). However, for spatial transcriptomics, framing the question of differential expression is more complex due to the range of dimensions along which genes can be differentially expressed, including spatial location, cell type, and condition (Figure 1); i.e., there are multiple ways in which one can define differential expression, some of which may be of greater biological interest than others. Statistical analyses for spatial transcriptomics have focused thus far on inference for a single tissue sample at a time, e.g., nnSVG (Weber et al., 2023), C-SIDE (Cable et al., 2022), SPARK-X (Zhu, Sun and Zhou, 2021), spatialDE (Svensson, Teichmann and Stegle, 2018), trendsceek (Edsgärd, Johnsson and Sandberg, 2018), among others. Rather than the conventional differential expression analysis just described, these methods address questions such as finding genes that are “spatially variable”, i.e., genes whose expression exhibits systematic variation across a given sample, forming discernible spatial patterns. Such spatial patterning might manifest as gradients of expression, localized hotspots, or regional differences that reflect underlying tissue structure, cell organization, or microenvironmental influences. Most methods also include a way to perform differential gene expression across groups, such as cell types, but only within a single tissue sample. As spatial transcriptomics technology is adopted more broadly, we can now collect data from multiple tissue samples. Existing methods do not adapt to this new setting.

**Figure 1:**
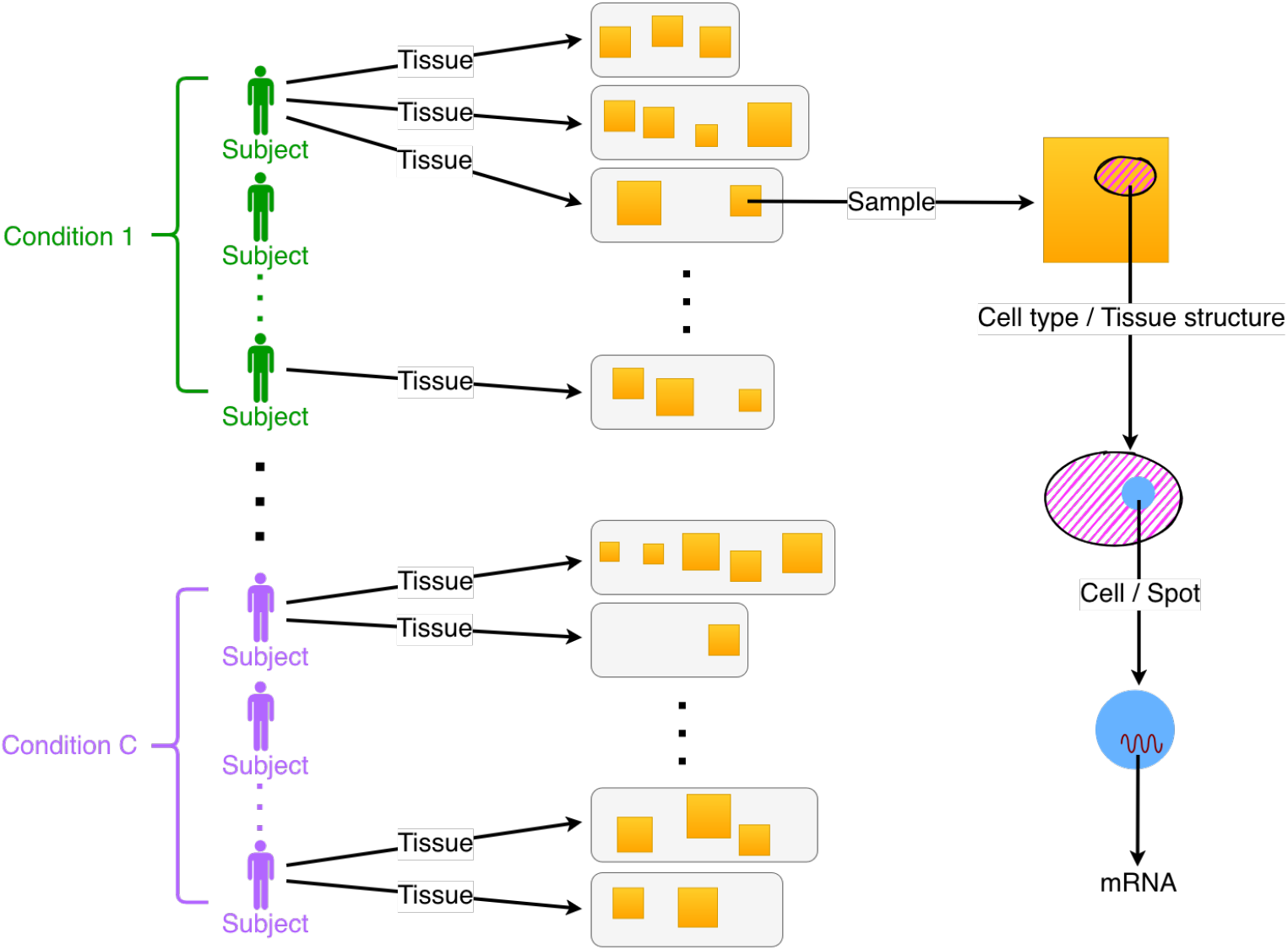
Experimental design and nomenclature. We present a general multi-sample spatial transcriptomics experimental design and associated nomenclature for discussing said design.

Multiple samples lead to unique challenges that traditional spatial statistical methods are not equipped to handle, such as different coordinate systems that do not necessarily align, different numbers and types of cells, different tissue architectures, etc. The geospatial analog would be analysis of data from disconnected islands, with highly complex and diverse within-island spatial structure. Existing approaches would analyze each island separately, whereas we seek to use all of the data at the same time (Freni-Sterrantino, Ventrucci and Rue, 2018). In the same analogy, we would seek to discover differences in some quantity across islands, while controlling for the spatial correlation within islands. In fact, the presence of data from multiple samples or islands is a benefit, as the greater sample size yields greater statistical power and generalizability relative to data from a single sample.

The multi-sample experimental setup, besides necessitating novel statistical methodology, enables us to ask and answer a richer set of biological questions, especially with regards to DE analyses. Consider that if the goal is to find differences in expression across diseases, a single sample comes from a patient with a single disease, and hence cannot alone be used to infer a disease effect. With multiple specimens, and hence multiple samples from different groups, we may now ask if there are differences in expression across diseases or phenotypes, while controlling for spatial correlation of measurements within the samples. Were we to apply a model designed for a single sample to our multi-sample setting, we would be restricted to estimating a set of fixed effects for each sample; in contrast, the question of interest calls for the estimation of a single set of fixed effects that are shared across all samples. In the example of spatial transcriptomics, an appropriately designed multi-sample experiment and corresponding model should allow the estimation of a common disease effect as well as a single, unambiguous set of cell type fixed effects.

### 1.3. Contributions and Outline

In this paper, we propose a novel method, `TESSERA` (Tool for Estimating Spatial and Sample-level Effects via Regression Analysis), for analyzing multi-sample spatial transcriptomics data. Mirroring its namesake in mosaic art, `TESSERA` integrates data from multiple individual samples—representing discrete “tessera” pieces— to estimate a single, unified set of fixed effects across samples, thereby enabling broader biological inferences that are not achievable when analyzing samples in isolation. To this end, we extend well-established single-sample spatial models to the multi-sample setting by allowing each sample to have its own spatial correlation structure while sharing a common set of fixed effects, and we provide a mathematical and computational framework that makes this extension efficient and scalable for model fitting and statistical inference. In particular, our framework explicitly leverages the sparsity in the four spatial correlation structures we consider. The `TESSERA` method is also novel within spatial statistics, as it provides a hitherto nonexistent extension to the multi-sample setting, and outperforms relevant prior baselines. It has applications beyond the biological focus of this manuscript, such as in urban planning or ecology. For example, we might identify drivers of economic activity within and across multiple metropolitan areas or study species interactions across multiple disconnected geographical regions.

We provide the mathematical specification of our `TESSERA` method in Section 2, with a description of our notation and spatial generalized linear mixed model (GLMM) in Section 2.1, an overview of covariance structures in Section 2.2, an outline of our model fitting method in Section 2.3, a framework for performing hypothesis testing in Section 2.4, and an overview of prior work in Section 2.5. We present a simulation study designed to validate the `TESSERA` method in Section 3, including power results in Section 3.4. We report on an application of the `TESSERA` method to a human kidney spatial gene expression dataset in Section 4. Details about data processing, design matrices, and computation are given in Section S1, and a longer discussion of the covariance structures is in Section S2. Detailed derivations and algorithmic specifications are given in Section S3. Theorems characterizing model identifiability are presented in Section S4, with proofs in Section S5.

The `TESSERA` method is implemented in an `R` package of the same name and available under the Artistic License 2.0. The package source code and documentation are currently available at https://github.com/floricaconstantine/TESSERA, and the code to reproduce the results in the manuscript is provided at https://github.com/floricaconstantine/TESSERA_manuscript. The package will be submitted to Bioconductor (https://www.bioconductor.org), the primary open-source repository for computational biology software, following publication.

## 2. The TESSERA Method

### 2.1. Model Specification

Consider the setting wherein we have *n* spatially-resolved samples indexed by *i* = 1, 2, …, *n*. For example, we might have disconnected samples from several tissue specimens from multiple subjects as seen in Figure 1. Within the samples, we have multiple measurements of a quantity of interest and their corresponding local (i.e., specific to the sample) spatial coordinates. In the example of samples from tissue specimens, we have numerous cells per sample, identified to spatial coordinates via the position of their centroids, and measurements of gene expression associated to each cell. To accompany each measurement, we might have values of covariates that can include cell-level or sample-level biological factors, such as sample condition, tissue structure, or cell type, as well as technical factors, such as the specific microarray slide or batch in which the cells were processed. We focus on modeling counts taken at the centroids, i.e., gene expression measures, as a function of these covariates. We want to model the counts with the primary goal of estimating fixed effects, shared across samples, for covariates of interest. However, unlike standard regression models used for non-spatial count data, such as scRNA-seq data, we have spatially-resolved count measurements and must account for the spatial correlation in our model.

In what follows, we will develop theory and methods for a single gene (or a single outcome or dependent variable in a general setting, of which spatial transcriptomics is a special case) at a time and will not use an index for genes to simplify notation. Let *Z*_*i,j*_ denote the count for location (cell) *j* in sample *i*, where there are *J*_*i*_ locations in sample *i*. We model the measured counts as

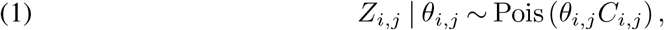

where the *Z*_*i,j*_ are assumed to be independent across samples (for different indices *i*) but not necessarily across locations within a sample (for different indices *j* and the same index *i*). Here, *θ*_*i,j*_ is an unobserved random variable that provides a location-specific mean which depends on the covariate fixed effects as well as a random effect that encapsulates spatial dependence. *C*_*i,j*_ is a known location-dependent scaling factor, which may be a random variable, e.g., the library size (i.e., total read count across all genes for cell *j* in sample *i*), in which case we condition on *C*_*i,j*_ for the distribution of *Z*_*i,j*_. Without loss of generality, one can set *C*_*i,j*_ = 1, if scaling is not needed for a given problem.

Applying a logarithmic link, we define *η*_*i,j*_ = log *θ*_*i,j*_ and let

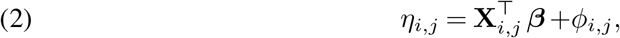

where **X**_*i,j*_ ∈ ℝ^*p*^ denotes known covariates to adjust for for location *j* in sample *i*, ***β*** ∈ ℝ^*p*^ is a vector of regression coefficients (the fixed effects) to be estimated, and *ϕ*_*i,j*_ is an unobserved spatially-varying random effect (Cressie, 2015, Section 7.5.2, p. 544–548). This is a very natural and well-known strategy for modeling count data with spatial correlation in the context of a single spatial sample (Stoehr and Robin, 2026; Murakami and Matsui, 2022; Chiquet, Mariadassou and Robin, 2021; Wang and Kockelman, 2013; Leroux, Lei and Breslow, 2000; Clayton and Kaldor, 1987). Importantly, the vector ***β***, our parameter of interest, has no index *i*, as it is shared across all samples. Additionally, we note that the random effects, *ϕ*_*i,j*_, while accounting for spatial dependence, also add over-dispersion between samples and cells beyond that of the Poisson model, similar to the negative binomial distribution for count data.

We model the vector 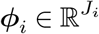, consisting of *ϕ*_*i,j*_, *j* = 1, …, *J*_*i*_, for sample *i*, as a multivariate normal random vector ***ϕ***_*i*_ ∼ 𝒩 (**0, ∑**_*i*_), with zero mean and a covariance matrix 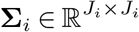 that captures spatial dependence between the locations. We assume that the *ϕ*_*i,j*_ are independent of the **X**_*i,j*_. Thus, we can write the model, which we call the `TESSERA` model, as

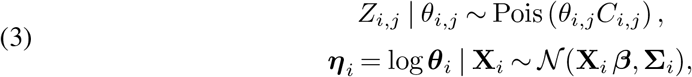

where 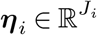 is the collection of *η*_*i,j*_, *j* = 1, …, *J*_*i*_, for sample *i*, and 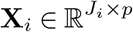 is the design/covariate matrix for sample *i*, with the *j*th row set to 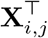 . The matrix **∑**_*i*_ defines how a given measurement’s nearest neighbors’ values affect its own value. For example, we might expect nearby cells in a tissue sample to influence each others’ gene expression.

We use ***ν*** to denote the entire set of model parameters for `TESSERA`, including the shared regression parameter ***β*** and the sample-specific parameters ***ψ***_*i*_ for the covariance matrices **∑**_*i*_ of the random effects ***ϕ***_*i*_, that is, ***ν*** = (***β, ψ***_1_,…, ***ψ***_*n*_). Section 2.2 below introduces two classes of models for the covariance structure.

The `TESSERA` method goes beyond the single-sample setting and enables modeling in the multi-sample setting. Importantly, we jointly model multiple samples with a single set of shared fixed effects ***β*** but a different covariance structures **∑**_*i*_ for each sample. The `TESSERA` method is also easily adaptable to many realistic settings and allows for statistical hypothesis testing concerning biological parameters of interest. The design matrices **X**_*i*_ can incorporate multiple covariates to account for complex study designs and, by testing contrasts of the elements of ***β***, we can perform DE analyses with respect to any quantity that can be encoded into the design matrices **X**_*i*_.

### 2.2. Models for Covariance Structure

The distribution of the spatially-varying random effects ***ϕ***_*i*_ is governed by their covariance matrices **∑**_*i*_. For the implementation of the `TESSERA` method, we assume the **∑**_*i*_ come from one of two classes of models for the covariance structure: Lattice-based models and sparse nearest-neighbor Gaussian process (spNNGP) models (Datta et al., 2016; Vecchia, 1988). Each of these have been proposed previously, only in the context of a single spatial sample. We show in Section S4 that, under relatively mild conditions, the multi-sample `TESSERA` model of Equation (3), coupled with any of the covariance models described below, results in parameters ***β*** that are identifiable.

In what follows, we briefly describe the two different classes of covariance models and how they are parameterized. We discuss the parameters involved for a single **∑**, with the understanding that for `TESSERA`, these parameters are defined separately for each sample, resulting in distinct **∑**_*i*_ for each sample (i.e., the subscript *i* is dropped to simplify notation). The models are discussed in greater detail in Section S2.

Both the lattice and spNNGP models specify a precision matrix **P** = **∑**^−1^ ∈ ℝ^*J×J*^, describing aspects of the spatial relationship between the observations in a given sample, but differ in how **P** is parameterized (Table 1). We denote by ***ψ*** the (typically unknown) parameters for **P** and, when relevant, use the longer notation **P**(***ψ***) to indicate the dependence of the precision matrix on these parameters.

**Table 1:** Parameterization of the precision matrix for covariance models considered for TESSERA. For the lattice models, the precision matrix **P** depends on two typically unknown parameters, τ ^2^ and γ, and a known cell-cell adjacency matrix **W**; **D** denotes the diagonal matrix with entries equal to the row-sums of **W**. For spNNGP, **K** is a kernel matrix that depends on some set of possibly unknown parameters **ψ** (e.g., number of neighbors, range, nugget, etc.) and the coordinates of the cells.

| Model | Precision matrix $\mathbf{P}$ | Contraints on parameters |
| --- | --- | --- |
| CAR | $(\mathbf{D} - \gamma \mathbf{W}) / \tau^2$ | $\tau^2 \in (0, \infty), \gamma \in (-1, 1), \mathbf{W} = \mathbf{W}^\top$ |
| SAR | $(\mathbf{I} - \gamma \mathbf{D}^{-1} \mathbf{W})^\top \mathbf{D} (\mathbf{I} - \gamma \mathbf{D}^{-1} \mathbf{W}) / \tau^2$ | $\tau^2 \in (0, \infty), \gamma \in (-1, 1)$ |
| Leroux | $[(1 - \gamma) \mathbf{I} + \gamma (\mathbf{D} - \mathbf{W})] / \tau^2$ | $\tau^2 \in (0, \infty), \gamma \in [0, 1), \mathbf{W} = \mathbf{W}^\top$ |
| spNNGP | $\mathbf{K}(\boldsymbol{\psi})$ | $\mathbf{K}(\boldsymbol{\psi}) = \mathbf{K}(\boldsymbol{\psi})^\top$ |

#### 2.2.1. Lattice-Based Models

For the lattice models, the precision matrix **P** is parameterized in terms of a scale parameter *τ* ^2^, a parameter *γ* controlling the strength of the correlation between neighboring observations, and a user-defined (i.e., assumed to be known) adjacency matrix **W** specifying the neighbors of each observation. Thus, ***ψ*** = (*τ* ^2^, *γ*) and the precision matrix may be written as **P**(***ψ***) = **P**^∗^(*γ*)*/τ* ^2^, where the matrix **P**^∗^ is also a function of the known **W**. We consider three lattice models: Simultaneous autoregressive (SAR) and conditional autoregressive (CAR) models (Besag, 1974), and a modification of the CAR model known as the Leroux model (Leroux, Lei and Breslow, 2000). For the CAR and SAR models, *γ*∈ (− 1, 1), and for the Leroux model, *γ* ∈ [0, 1), where we exclude *γ* = ±1 to ensure that **P** is positive-definite. The exact role of *γ* differs between the models. In the CAR and SAR models, *γ* quantifies the spatial dependence between neighboring observations. In the Leroux model, the random effect is modeled in terms of both a spatially-correlated component as well as a non-spatially-correlated component, so that *γ* defines a tradeoff between i.i.d. observations at *γ* = 0 and an improper CAR model (i.e., CAR with *γ* = 1) at *γ* = 1. In all models, *γ* controls the strength of the correlation between neighboring observations. Unlike the Leroux models, the CAR and SAR models cannot represent i.i.d. data for arbitrary **W**; setting *γ* = 0 leads to independent but not identically distributed measurements. Importantly, as each location or cell has only a handful of direct neighbors, the matrix **P** is extremely sparse. This sparsity enables scalable fitting and inference for our model. In application to spatial transcriptomics, **W** could be defined by determining neighboring cells or, if the cell boundaries are not available, by thresholding the distances between cell centroids. While the SAR model allows an asymmetric **W**, both the CAR and Leroux models require **W** to be symmetric; thus we will assume a symmetric **W** in our applications. We also require that **W** have no isolated points to ensure that **P** is invertible in all models, though not strictly required for Leroux.

#### 2.2.2. Sparse Nearest-Neighbor Gaussian Process Models

A Gaussian process model for **∑** is of the form **K**(***ψ***), for some kernel matrix **K** that is parameterized by ***ψ***. For *J* locations in a sample, **K**(***ψ***) is a *J ×J* matrix, where the (*j, j*^*′*^) entry is a function of the locations of the measurements with indices *j* and *j*^*′*^. Common choices for the kernel function include the exponential, Gaussian, and Matérn kernel functions; these functions have the advantage of isotropy, wherein they only depend on the distance between pairs of locations, as opposed to the locations themselves (Rasmussen and Williams, 2006, Chapter 4). We use the Matérn kernel as the default for `TESSERA`. Along with spatial variance (partial sill), range, and nugget parameters, this kernel includes a smoothness parameter that provides the flexibility to represent a wide range of spatial processes. A smoothness parameter of 0.5 yields the exponential kernel and the Gaussian kernel is recovered in the infinite limit.

While Gaussian process models offer extreme flexibility in capturing spatial structure, they also lead to computational challenges. In particular, the need to consider all pairs of locations leads to a dense matrix for **∑**. The authors in Vecchia (1988) noted that considering only the *k* nearest neighbors for each location yields a sufficiently accurate approximation to **∑** for a user-specified *k* as small as 10–20. This suggests using a sparse nearest-neighbor Gaussian process, which assumes **∑** = **P**^−1^, where **P** is a sparse version of **K**(***ψ***), with entries of **K**(***ψ***) set to 0 outside of the *k* nearest neighbors (Datta et al., 2016). This choice enables scalable fitting and inference for our model, as in the lattice model setting. In spatial transcriptomics, the distances between cell centroids would be provided, **K**(***ψ***) would be calculated, and the sparse version **P** would be based on a user-specified parameter *k* for the number of neighbors to consider.

### 2.3. Fitting the TESSERA Model

Fitting the multi-sample `TESSERA` model necessitates novel estimation and algorithmic procedures. A natural framework for solving this problem is maximum likelihood estimation. However, a direct maximization of the likelihood function with respect to ***ν*** would involve a high-dimensional integral over the ***ϕ***_*i*_ and would not scale well. A Monte Carlo approach could approximate the integral, but would also require large amounts of sampling and computation. Instead, the `TESSERA` method is based on the expectation-conditional-maximization (ECM) algorithm, which is an iterative algorithmic framework for estimating parameters where the data-generating model depends on unobserved latent variables (Meng and Rubin, 1993).

Briefly, the ECM fitting procedure modifies the maximization step in the expectation-maximization (EM) algorithm to alternate between maximizing over different portions of the parameter vector ***ν*** separately, resulting in the following steps for the `TESSERA` model.

- **Expectation (E) step**. Compute the expected value of the joint log-likelihood *L*(***ν***; **Z, *η***) of the observed counts **Z** and unobserved variables ***η***, conditional on the observed counts, the covariates **X**, and current estimates 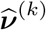of the unknown parameters ***ν***,

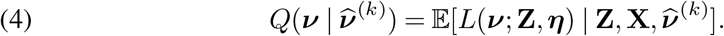
- **Conditional maximization (CM) step**. Sequentially maximize 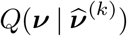 with respect to ***ν*** for a single element (or subset of elements) of ***ν*** at a time (Dempster, Laird and Rubin, 1977; Meng and Rubin, 1993).

Generally, the E-step simplifies to calculating 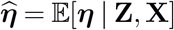 or other moments of ***η*** | **Z, X**, and the ‘conditioning’ here results in sequentially maximizing over one parameter from ***ν*** while holding the other parameters in the optimization constant. ECM is useful when a global maximization of 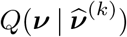 is slow, but conditional updates can be done rapidly (or analytically). The results of M eng and Rubin (1993) show that maximizing the conditional likelihood is an approximation for the standard M-step.

Recall that for `TESSERA` the set of model parameters ***ν*** are the shared regression parameter ***β*** and the sample-specific parameters ***ψ***_*i*_ for the covariance structure of the random effects ***ϕ***_*i*_. Due to the intractable log-likelihood, we use a Gaussian approximation proposed by Clayton and Kaldor (1987) for the distribution of ***η***| **Z, X** in the E-step (see Section S3). For the CM-step, we fix ***β*** and sequentially optimize in each sample’s spatial correlation parameters ***ψ***_*i*_, then optimize in ***β*** while holding the sample-specific correlation parameters fixed; this is iterated until convergence of 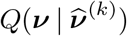. EM algorithms are commonly used for single-sample spatial models (Clayton and Kaldor, 1987). The conditional maximization step of `TESSERA`, where each sample’s spatial correlation parameters are optimized separately per sample, results in an algorithm that is similar structurally to an EM algorithm per sample, but with the added optimization of the shared ***β*** parameter globally over all of the samples. The details of the `TESSERA` fitting algorithm are presented in Section S3 (see pseudocode in Algorithm 1).

The `TESSERA` algorithm can work with arbitrary covariance structures, but we focus on the lattice and spNNGP models, both of which have a sparse precision matrix **P**. This sparsity enables an efficient, scalable implementation of the ECM framework. Relative to a method based on Markov chain Monte Carlo (MCMC) that must generate thousands of posterior draws, `TESSERA` is able to achieve faster computational runtimes (Supplementary Figure S-1), with the caveat that, unlike exact MCMC methods, an ECM algorithm only guarantees convergence to a local optimum of the log-likelihood. Moreover, while approximations like the integrated nested Laplace approximation (INLA) can replace MCMC samplers and offer dramatic speedups at minimal loss of accuracy, INLA scales exponentially in the number of hyper-parameters, in this case, the spatial covariance parameters (Rue, Martino and Chopin, 2009; Opitz, 2017). The ECM framework, on the other hand, maintains a linear scaling in the number of parameters. Hence, the `TESSERA` method, through its ECM fitting procedure, can directly model count data while scaling to large numbers of samples and measurements, all with low computational overhead and fast time-to-results.

#### Algorithm 1

The `TESSERA` algorithm.

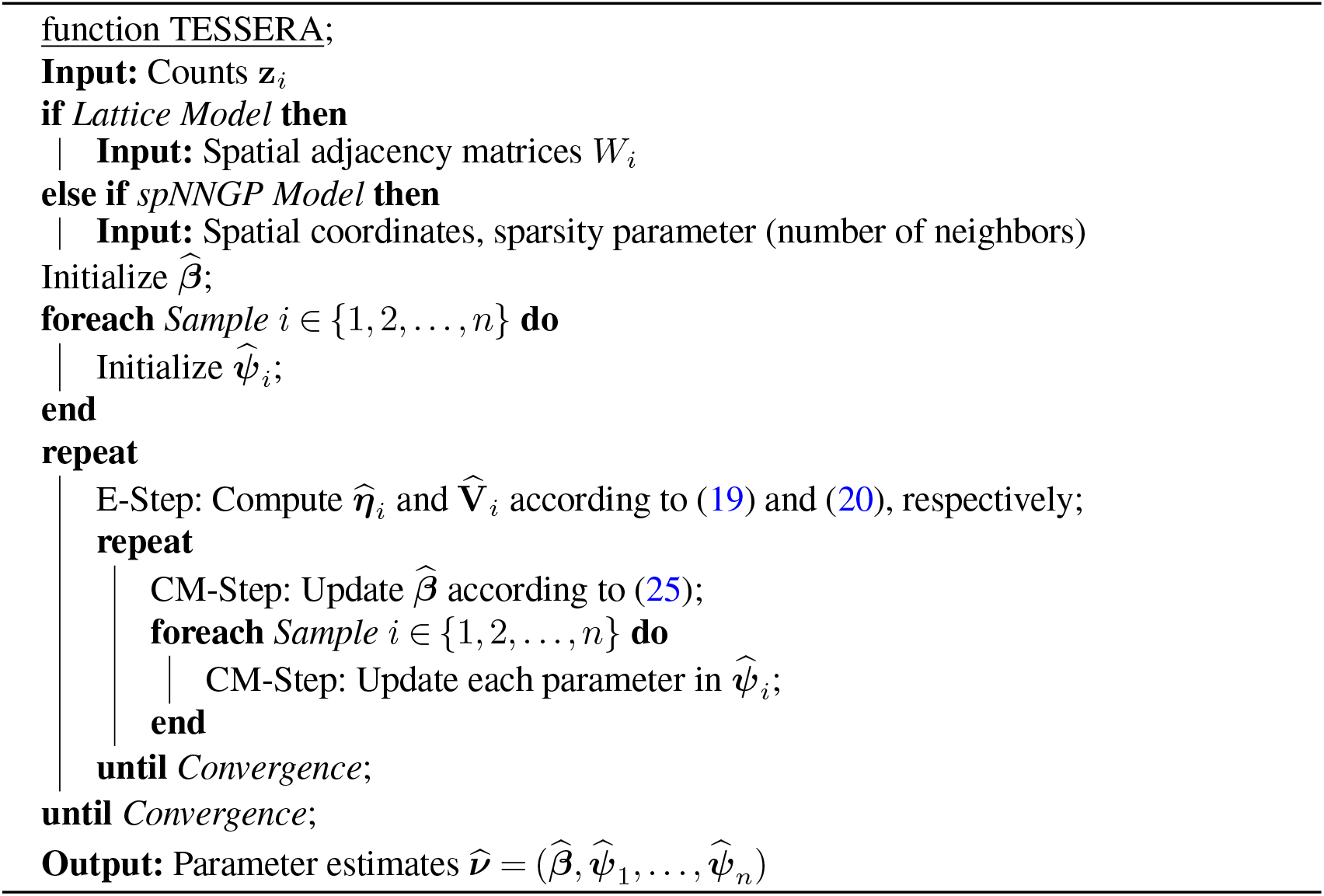

### 2.4. Hypothesis Testing

Given an appropriate design matrix, we can use the fitted 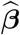 from `TESSERA` to perform differential expression analyses, where the precise parameterization of ***β*** depends on the scientific question. We distinguish between two primary classes of covariates: within-sample (observation-level) covariates, which vary across cells or spots (e.g., cell type or tissue structure), and across-sample (sample-level) covariates, which remain constant within an individual sample but vary across samples (e.g., disease condition or treatment group). Testing the associated elements of 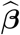 allows us to address different types of DE questions. We may identify genes differentially expressed between cell types or tissue structures, or test for interaction effects with across-sample covariates like disease condition. By accounting for these interactions, we can detect differences specific to a given cell type or structure, thereby revealing how disease effects vary across different tissue contexts. More generally, we can perform hypothesis testing for any covariate that can be encoded in a design matrix by testing relevant contrasts of the elements of 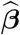. Details for the design matrices and associated ***β*** parameterizations used in our simulated and real data analyses (Sections 3 and 4) are provided in Section S1.2.

To perform statistical hypothesis testing for the differential expression questions we consider, we may invoke the asymptotic normality of the maximum likelihood estimator, even in the case of dependent data as in spatial transcriptomics—see Crowder (1976) and Lehmann and Casella (2006, Section 6.7, p480-481). Indeed, the likelihood function for the `TESSERA` model herein is smooth and sufficiently regular. Using the fact that the maximum likelihood estimator is asymptotically normal, with mean equal to the true underlying parameter value and covariance equal to the inverse of the Fisher information, we may specify a Wald test statistic for contrasts of the parameters, e.g., of the fixed effects ***β*** and of the spatial correlation parameters *γ*_*i*_. Indeed, consider a general parameter vector ***ξ*** *∈* ℝ^*d*^, *q* linear contrasts encoded in a matrix **C** *∈* ℝ^*q×d*^, and a vector of constants **c** *∈* ℝ^*q*^. Define the null hypothesis *H*_0_ and the alternative hypothesis *H*_1_ as follows

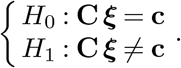

Given the maximum likelihood estimator 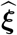 of ***ξ***, with Fisher information **V** and observed information 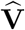, the Wald test statistic is defined as

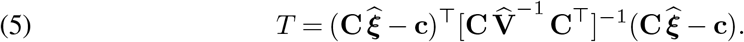

Classically, under appropriate assumptions and regularity conditions, the Wald test statistic *T* converges in distribution to a 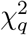 random variable (Wald, 1943). However, the asymptotics of Wald statistics are poorly understood for generalized linear mixed models, and the standard approaches of degree-of-freedom estimation are not applicable to our ECM procedure (Stroup, 2013, 2015; Bell and Grunwald, 2011). Moreover, in an ECM procedure, it is unclear how to obtain standard errors. Hence, we propose the following Wald test procedure, inspired by the empirical null distribution estimation procedure from Efron (2004).

From the Hessian matrix of the expected log-likelihood, we may obtain estimated standard errors for 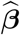 (Meilijson, 1989). Given these standard errors, we may form Wald test statistics in the usual manner, as in Equation (5). We might then take inspiration from the methodology in the `limma` package (Smyth, 2004) and note that, while we do not know the exact distribution of the Wald statistics and the asymptotics of the Wald statistics are measured in terms of the number of samples as opposed to the number of cells/locations, it is plausible that the Wald statistic for a single contrast behaves like a scaled 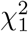 random variable. Moreover, for finite samples, the bias in the Hessian and in the estimator of ***β*** might be non-trivial; allowing for a non-centrality parameter could account for this bias. The problem is that we do not know, a priori, what the scale and non-centrality parameters are of this scaled non-central 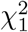 distribution. If we had some threshold below which the set of Wald statistics was comprised mostly of null cases, we could use it to estimate the 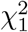 parameters.

We propose the following scheme to automatically choose the threshold. Under the null hypothesis, the *p*-value for a test statistic with a continuous density function should have a uniform distribution on [0, 1], 𝒰 (0, 1). For a given contrast, let *T*_*g*_ denote the Wald test statistic for gene *g* = 1, 2, …, *G*. Assume that the *T*_*g*_ corresponding to true null hypotheses follow a scaled non-central chi-squared distribution, 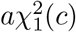. To try to isolate those test statistics, for a given threshold value *t*, define the set 𝒯 (*t*) ={*T*_*g*_ : *T*_*g*_ ≤*t*} from which to estimate *a* and *c*. However, since we are conditioning on *T*_*g*_ being less than *t*, we will need to estimate *a* and *c* for a 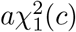 distribution conditioned to be less than *t*. We estimate the parameters by numerically maximizing the conditional likelihood in *a* and *c*.

To determine the best choice of cutoff *t*, we compute the *p*-values *p*_*g*_(*t*) of the *T*_*g*_ *∈* 𝒯 (*t*) assuming their null distribution is 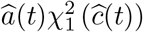, and compare these *p*-values to a 𝒰 (0, 1) distribution. Let *p*_*g*_ (*t*) = ℙ (*X*≥ *T*_*g*_), for 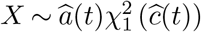, where 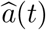 and 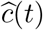 are estimates of the scale and non-centrality parameters. Given the *p*-values *p*_*g*_(*t*), compute the squared *L*^2^ error between their empirical CDF and the CDF of the uniform distribution on [0, 1]:

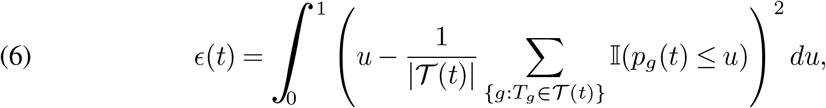

where 𝕀 is the indicator function. Hence, for each threshold *t*, we obtain an error *ϵ*(*t*); we choose the threshold *t* that minimizes *ϵ*(*t*).

As in other genomic settings, and as for virtually all differential expression methods, probability statements related to *p*-values rely on hard-to-verify assumptions and often do not account for prior preprocessing steps. We therefore refrain from attaching strong probabilistic interpretations to *p*-values and use these mostly as useful numerical summaries to determine which genes exhibit differential expression. Empirical evidence for Type I error control and power can be obtained from simulation studies or by reference to known biology, as in Sections 3 and 4 below.

### 2.5. Comparison with Prior Work

While spatial count models, such as Poisson models with spatially-varying random effects and non-spatially-varying fixed effects, have been studied extensively (Glaser, 2017; Dass, Lim and Maiti, 2010), existing models can only handle a single sample at a time. Here, we have multiple disconnected samples with no common coordinate system and want to estimate a single set of fixed effects across samples, while estimating the spatial random effects within each sample. In this section, using our notation, we describe the evolution of non-spatial to spatial models as well as the extension of single-sample to multi-sample settings, and describe the limitations of existing methods.

#### 2.5.1. Non-Spatial Models

The simplest applicable model for our data is a non-spatial generalized linear model based on a Poisson distribution applied to a single sample. In our notation, we would have observations *Z*_*i,j*_ where

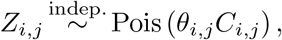

and

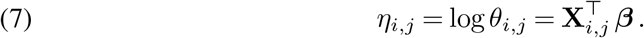

Note that this model can apply to a single sample, in which case we would drop the sample index *i*. Importantly, this model treats all cells or measurements as independent, even if they come from the same sample. Thus, if applied to multiple samples, this model considers all the cells to be thousands of independent observations.

An alternative model would be to consider a normal distribution, instead of a Poisson. Though we observe counts *Z*_*i,j*_, we might consider logarithmically transforming them as log(*c* + *Z*_*i,j*_), for some constant *c*, typically 1 or 1*/*2 (Booeshaghi and Pachter, 2021; Lun, 2018). Then, the GLM for the normal distribution is equivalent to a simple linear model,

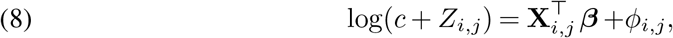

where 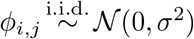 for some variance parameter *σ*^2^ *>* 0. It should be noted that the log-transformed counts are not truly continuous and treating them as such does not capture the essence of the data. In particular, in spatial transcriptomics, the observed data are quite sparse and, for many genes, the values *Z*_*i,j*_ range mostly between 0 and 3. This will be a concern for all methods discussed here that use Gaussian models on log-transformed counts, rather than modeling the data with a count distribution.

Though not typically considered as a spatial model, a generalized additive model (GAM) can account for spatial information using functions of the spatial coordinates (*x*_*j*_, *y*_*j*_) of the measurements as covariates (Wood, 2017). Models of this form have been considered for spatial transcriptomics data in Cable et al. (2022). A common form for these functions is a smoothing spline, e.g., a thin-plate spline, and these models can be fit via maximum likelihood or restricted maximum likelihood approaches. It is also possible to fit a model with a per-sample spatial spline component and a common set of fixed effects via the `mgcv` package (Wood, 2017).

#### 2.5.2. Single-Sample Spatial Models

As already mentioned, our `TESSERA` model is a generalization of models described previously in the literature for a single spatial sample, which account for spatial dependency between locations by adding a random effect to the Poisson GLM of (7),

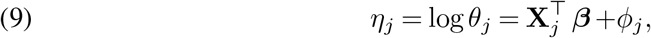

where *ϕ*_*j*_ are potentially correlated zero-mean Gaussian random variables. The CAR, SAR, Leroux, and spNNGP models correspond to different ways to describe the correlation structure of the *ϕ*_*j*_.

Other methods have instead extended the Gaussian model of Equation (8) by assigning a spatial correlation structure to the *ϕ*_*j*_. Of particular interest in the Gaussian setting is the `BRISC` method (Saha and Datta, 2018), which assumes continuous observations and applies a spNNGP structure to *ϕ*_*j*_ of Equation (8). The algorithm in `BRISC` builds on recent work on sparsification via nearest-neighbor approximations to accelerate the fitting of Gaussian processes. As a point of interest, the nnSVG method proposed for spatial transcriptomics uses the `BRISC` algorithm to test a single sample for the presence of spatial patterning, i.e., testing for the existence of spatial dependencies (Weber et al., 2023).

#### 2.5.3. Block-Wise Spatial Models

Another class of single-sample methods supports disjoint or disconnected partitions of the sample, wherein each partition corresponds to a distinct connected component. This results in block covariance matrices, with the blocks corresponding to the different partitions. This is similar to our setting, where separate samples correspond to disjoint partitions. Indeed, the `TESSERA` model could be equivalently written with random effects having a block covariance matrix, with blocks defined by **∑**_*i*_.

One such model is the hierarchical spatial autoregressive (HSAR) model from Dong and Harris (2015), which builds on the Gaussian model in (8) with a SAR covariance structure for *ϕ*_*i,j*_, but allows for a block-wise structure of the SAR adjacency matrix, i.e., an adjacency matrix **W** that is not fully connected. However, the SAR parameters are shared across all samples, that is, there is only a single *τ* ^2^ and *γ*. If we expect the spatial structure to change across samples, this model is clearly not the right fit for our data. Estimation of HSAR is accomplished via MCMC, though, at the time of writing, the relevant HSAR package is no longer in the `R` CRAN package repository. It should be noted, also, that HSAR assumes a Gaussian distribution, thus must be applied to the log-counts, with the ensuing problems mentioned above.

#### 2.5.4. Multivariate Models

Another related model is the *multivariate* CAR (MCAR) model from Jin, Carlin and Banerjee (2005). This model assumes a single set of spatial locations indexed by *j* and allows for multivariate data **Z**_*j*_ ∈ ℝ ^*K*^ to be collected at each *j*, e.g., expression measures for different genes for cell *j*. This gives a model of the form

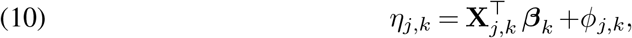

where the counts *Z*_*j,k*_ are generated according to a Poisson distribution with parameter *θ*_*j,k*_ = exp *η*_*j,k*_. Here, each outcome *k* has a corresponding covariate matrix **X**_*k*_ and its own fixed effect vector ***β***_*k*_ . The covariance structure of the random effects *ϕ*_*j,k*_ allows for dependence both spatially for each outcome and across outcomes, with acommon CAR structure for each outcome. The CAR structure has a correlation parameter *γ* that is shared across outcomes, but the scale parameter 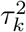 can vary between outcomes, as can the strength of correlation between pairs of outcomes.

In principle, this model has similar components to our `TESSERA` model, if we let the outcomes *k* be instead samples *k*. However, the very differing contexts for the two models result in different structural constraints: First, we wish to estimate a single set of fixed effects whereas the MCAR ***β***_*k*_ vary across samples/outcomes; second, we do not want to restrict ourselves to the setting of identical spatial locations across samples; and third, we want to allow for the values of the correlation parameter to vary across samples.

#### 2.5.5. Model Fitting

In the single-sample setting, MCMC methods for fitting spatial Poisson models have been used successfully for small datasets (e.g., a few hundred measurements) (Zhang, 2002; Christensen, 2004). However, these methods often have problems for larger datasets, including slow mixing times due to correlation of the observations (here, the counts *Z*_*i,j*_), expensive calculations and evaluations of the complicated likelihood function, and the need to repeatedly sample a high-dimensional vector of random effects (Guan and Haran, 2018, 2019). Beyond the issues with MCMC, many of these models involve complex covariance structures, e.g., Matérn or Gaussian covariances, for the spatial random effects, and for large numbers of measurements (cells, in this case), these covariance structures lead to large dense matrices and high computational demands (Sun, Zhu and Zhou, 2020; Zhu, Sun and Zhou, 2021; Dupont, Wood and Augustin, 2022; Anderson et al., 2022). While sparse approximations as in `BRISC` (Saha and Datta, 2018) help, they do not solve the underlying issues of the MCMC: speed and slow mixing. Moreover, off-the-shelf implementations of MCMC methods for the Poisson spatial model are relatively few, and a non-trivial degree of expertise is required to implement them from scratch—the same goes for alternatives such as Laplace approximations like INLA (integrated nested Laplace approximation) (Rue, Martino and Chopin, 2009). The INLA approximation replaces the MCMC sampler and is generally faster; examples of INLA for single-sample spatial models include the work in Vicente et al. (2023). Of particular note is the fact that, unlike existing off-the-shelf MCMC implementations in `R`, the `R-INLA` implementation can be used to specify and fit a multi-sample Poisson spatial model with distinct spatial correlation structures for each sample. However, as the timing results in Supplementary Figure S-1 indicate, this procedure is not scalable to realistically sized spatial transcriptomics datasets. For INLA, each additional sample requires estimating additional hyper-parameters (spatial covariance parameters) but the computational scaling is exponential in the number of hyper-parameters (Rue, Martino and Chopin, 2009; Opitz, 2017). Simpler than both MCMC and INLA is the EM algorithm applied to a single sample in Clayton and Kaldor (1987). At the cost of some convergence guarantees (global vs. local optima) and a slight loss of accuracy, the ECM algorithm offers simplicity and speed (see Supplementary Figure S-1).

## 3. Simulations Studies

### 3.1. Synthetic Data Based on Real Data

To evaluate the performance of `TESSERA`, we conduct a series of simulation studies based on the model defined in Equation (3). We generate data using two distinct spatial covariance models, Leroux and spNNGP, to assess the method across different types of spatial dependence. To ensure biological realism, we choose the parameters ***ν*** for these simulations by applying `TESSERA` to the kidney spatial transcriptomics dataset described in Section 4 and retain the original spatial coordinates and covariates (cell type and disease condition). Within a given simulation study, the datasets are generated independently across trials from the specified Poisson-Leroux or Poisson-spNNGP distribution for the *Z*_*i,j*_, conditioned on these fixed parameters. We benchmark `TESSERA` against a Poisson and a negative binomial (NB) GLM, as well as a generalized additive model. Where applicable, we also compare our method to pseudobulk methods `DESeq2` and `limma-voom`, which aggregate expression measures within each sample and cell type (Lun and Marioni, 2017). We assess performance in terms of accuracy for estimating model parameters, model fit, power to detect differential expression, and robustness to model misspecification.

### 3.2. Performance of TESSERA on Multi-Sample Data

We focus on two genes, the ones with the highest and lowest variances of the raw counts across all cells and samples, thereby representing two extremes of the data. For each gene and data-generating scenario, we conduct 100 simulation trials and, for each trial, apply every method to be evaluated.

#### 3.2.1. Estimation Accuracy

When data are generated from a Poisson-Leroux distribution, the results demonstrate that `TESSERA` applied using this model consistently outperforms the other methods in estimating the fixed effects ***β*** in terms of the relative squared error, as shown in the left panel of Figure 2. `TESSERA` also produces accurate estimates of the parameters *γ* and *τ* ^2^ for the spatial covariance matrix. We see that the error in *γ* ∈ [0, 1) is generally low (Supplementary Figure S-2). For *τ* ^2^, the absolute error is low for genes with true values near 0 (Supplementary Figure S-3). The relative error is consistent across genes and reflects a slight bias toward underestimating *τ* ^2^ (Supplementary Figure S-4).

**Figure 2:**
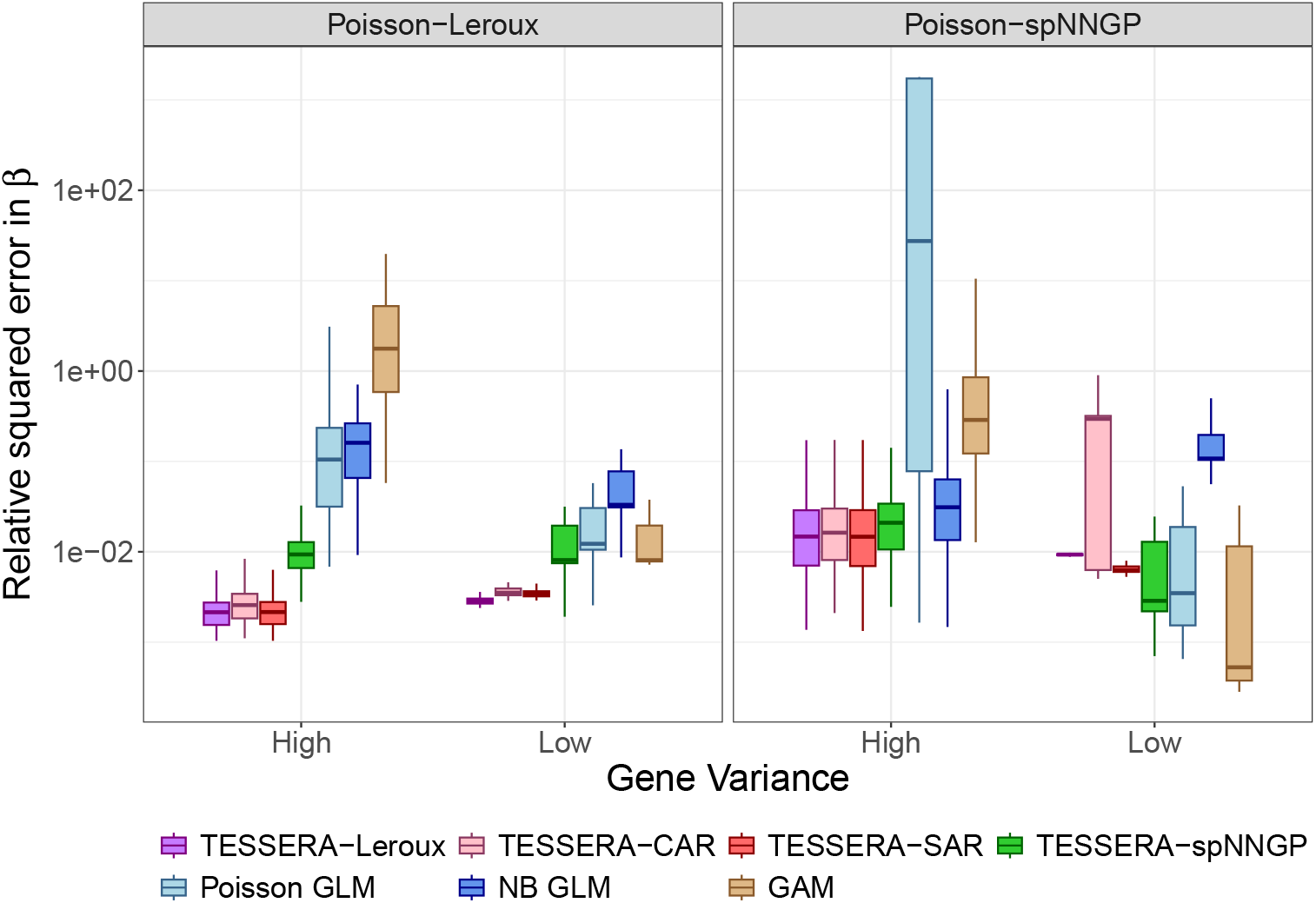
Simulation study: Relative error in estimating the shared regression coefficient *β*. Shown are boxplots of the relative squared error in estimating ***β***, 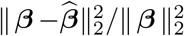, across 100 trials of simulated data. Different colored boxplots represent different methods for estimating the parameter ***β***: `TESSERA` with different choices of spatial covariance structure, Poisson GLM, negative binomial GLM, and GAM. We generate synthetic data based on the kidney dataset from Abedini et al. (2024), by sampling from both the multi-sample Poisson-Leroux (left) and Poisson-spNNGP (right) distributions with parameters derived from fitting the model to the original, real data. We show results for two sets of parameters, those corresponding to the genes with the maximum (“High”) and minimum (“Low”) variances of the raw counts across all cells and samples.

Notably, even when the model is misspecified, with data generated from a Poisson-spNNGP model, `TESSERA` (applied with a Poisson-Leroux model) still performs better than other methods for high-variance genes, as seen in the right panel of Figure 2. For low-variance genes, `TESSERA` maintains low relative squared error in ***β***, indicating the method’s robustness even in the presence of model misspecification. For genes with variances falling between the extremes, the results are comparable, as demonstrated in Supplementary Figure S-5.

#### 3.2.2. Capturing Spatial Autocorrelation

To compare methods in terms of their ability to account for spatial autocorrelation, we compute the Moran’s *I* (see Definition 4) of their residuals and contrast it to the Moran’s *I* of residuals for data generated using an independent baseline model (with *γ* = 0). Use of the independent baseline allows us to differentiate between capturing genuine spatial structure and potential artifacts of model complexity. Across both the Poisson-Leroux and Poisson-spNNGP data-generating scenarios, `TESSERA` fit with lattice-based models consistently outperforms `TESSERA` fit with a spNNGP model and the non-spatial methods, by more effectively accounting for spatial autocorrelation. While other models frequently leave behind residual autocorrelation or show high variability in performance, the lattice-based models yield residuals that more closely align with spatial independence regardless of the underlying generative process (Supplementary Figures S-6 and S-7).

#### 3.2.3. Model Fit

`TESSERA` fit with the multi-sample lattice models provides the best fitted counts in terms of mean squared error (MSE), with errors an order of magnitude smaller than those from the GLM and the GAM; `TESSERA` fit with the multi-sample spNNGP model shows errors typically falling in between the two (Supplementary Figure S-8). This indicates that explicitly modeling spatial dependence within samples can substantially improve model fit. Mirroring the analysis in Section 3.2.2, the benefit of explicitly accounting for spatial dependence is made even more apparent when we examine the difference in normalized MSE for spatially correlated data relative to the independent (*γ* = 0) baseline. While the Poisson GLM, NB GLM, and GAM exhibit inconsistent and often substantial shifts in MSE when spatial dependence is introduced, indicating that the fit of non-spatial models is highly sensitive to gene-specific spatial patterns, `TESSERA` variants maintain a comparatively stable error profile. Across all genes and data-generating scenarios, `TESSERA` shows at most relatively small levels of increased MSE in response to the more complex setting of spatial dependence (Supplementary Figure S-9).

### 3.2.4. Recommended Default Covariance Model for TESSERA

In summary, these results reinforce that the `TESSERA` method, particularly with lattice-based models, not only provides accurate estimates of fixed effects but also excels in minimizing residuals and accounting for spatial autocorrelation, outperforming conventional methods like GLM and GAM. In what follows, for analyses where we only fit `TESSERA` for a single model, we use the Poisson-Leroux lattice model as our default; its covariance structure incorporates both spatial and non-spatial components—unlike the Poisson-CAR and Poisson-SAR models—and its spatial parameters are identifiable, in contrast to the Poisson-spNNGP model.

### 3.3. Performance of TESSERA on Single-Sample Data

Although `TESSERA` is designed for a multi-sample setting, we validate the method on single-sample data to benchmark it against existing single-sample spatial methods. For this purpose, we generate data using the estimated parameters from a representative sample, *HK3035_ST*, within the kidney dataset (Abedini et al., 2024). We again focus on the two genes with the highest and lowest variances of the raw counts across all cells and samples in the entire kidney dataset. We conduct 100 simulation trials for each combination of gene and data-generating scenario and, for each trial, apply every method to be evaluated. We assess model performance by examining the relative squared error for the fixed effects ***β*** within a single-sample framework in Supplementary Figure S-10. `TESSERA` fit with lattice models consistently demonstrates estimation performance comparable to the MCMC and INLA implementations that are commonly used in spatial statistics. This indicates that `TESSERA`’s computational efficiency does not come at the cost of accuracy. `TESSERA` fit with the Poisson-spNNGP model exhibits superior performance relative to BRISC, though the `BRISC` algorithm fits a Gaussian model to log-transformed counts and thus has a different underlying model. Notably, non-spatial Poisson and NB GLM exhibit significantly higher error rates, by several orders of magnitude, highlighting the necessity of accounting for spatial autocorrelation.

In the aforementioned analysis, we dropped four cells from the *HK3035_ST* sample, corresponding to two cell types: one with three cells and another with only one cell. While the corresponding entries of ***β*** are technically estimable, leaving these cells in the dataset might lead to worse results. To examine the effect of these small cell groups or of high variance elements in 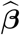, we present the relative squared error of the fixed effects when we do not drop these four cells in Supplementary Figure S-11. The MCMC results show a substantial degradation in performance, characterized by much larger variances and error values. This pattern also appears when using INLA for genes with low variance. In contrast, the ECM-based `TESSERA` results remain stable, leading us to conclude that `TESSERA` is robust to the presence of these small subgroups.

### 3.4. TESSERA Power and Type I Error Control

#### 3.4.1. Simulation Model

The power simulations remain grounded in the kidney spatial transcriptomics dataset (Abedini et al., 2024), where the elements of ***β*** correspond to interactions between cell type and disease condition and to sample-specific main effects. For these simulations, however, we focus on a single gene and modify the specification of its ***β*** vector to concentrate on the detection of differences across disease conditions for a given cell type. Concretely, we begin with the 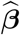 from either a Poisson-Leroux or Poisson-spNNGP model fit to a high-variance gene (*IGHG2*). For a given cell type, we set the coefficient corresponding to the control condition to the average of all coefficients in 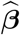. We then vary the coefficient corresponding to the diabetic kidney disease (DKD) condition by adding a value in [−2, 2] to the value of the control coefficient to create an effect size for the contrast. We do this separately for each of four cell types, with on average approximately 40, 75, 165, and 600 cells per sample, to cover a wide range of realistic scenarios (see Supplementary Figure S-12 for the range of numbers of cells per sample). To evaluate power, we perform 100 simulation trials for each combination of contrast value and data-generating scenario and, for each trial, apply every method to be evaluated. Power is then calculated as the proportion of trials in which we reject the null hypothesis that the control-cell type and DKD-cell type interaction coefficients are equal.

To test for differential expression with `TESSERA`, we perform Wald tests following the procedure in Section 2.4. The parameters of the scaled non-central chi-squared null distribution are estimated using Wald statistics derived from the real data. Specifically, we fit a Poisson-Leroux model to all 3,000 genes and compute Wald statistics for all within-cell-type, across-condition contrasts. This collection of (3,000 genes × 3 condition pairs × 26 cell types) statistics yields a 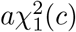 null distribution, which is then used to compute *p*-values for the Wald statistics generated in our power simulations. As detailed in Section S3.2, the covariance of 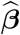 is used to compute these Wald statistics.

We benchmark the performance of the `TESSERA` method against the GLM and GAM alternatives as well as pseudobulk methods `DESeq2` and `limma-voom`. To accommodate the multi-gene dispersion estimation required by pseudobulk methods, we fit their models using all available genes but replace a single gene with synthetic data generated by modifying 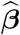 as described above.

While these simulations focus on a single gene, practical applications typically involve simultaneous testing across thousands of genes, bringing additional considerations such as multiple testing adjustment. Additionally, the parameters of the 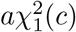 null distribution were estimated once based on a single realization of the data (specifically, that of the real data), thus missing in the simulations the variability in that estimation process. To capture this variability, these parameters would have to be re-estimated across a full set of simulated genes for every realization of the data, which is computationally prohibitive.

#### 3.4.2. Characterizing the Power and Type I Error Rate Trade-off

We now consider the relative performance of `TESSERA` in detecting differentially expressed genes. We generated receiver operating characteristic (ROC) curves across a broad range of effect sizes and cell types for the DKD vs. Control condition comparison. Representative results are shown in Figure 3 for the *C_TAL* cell type (∼600 cells per sample) for two effect sizes; ROC curves for a wider range of values are provided in Supplementary Figures S-13 and S-14. The ROC curves illustrate that `TESSERA` consistently outperforms or matches the performance of other methods. Furthermore, `TESSERA` has consistently high performance, unlike other methods whose performance can vary greatly for different simulation models or effect sizes.

**Figure 3:**
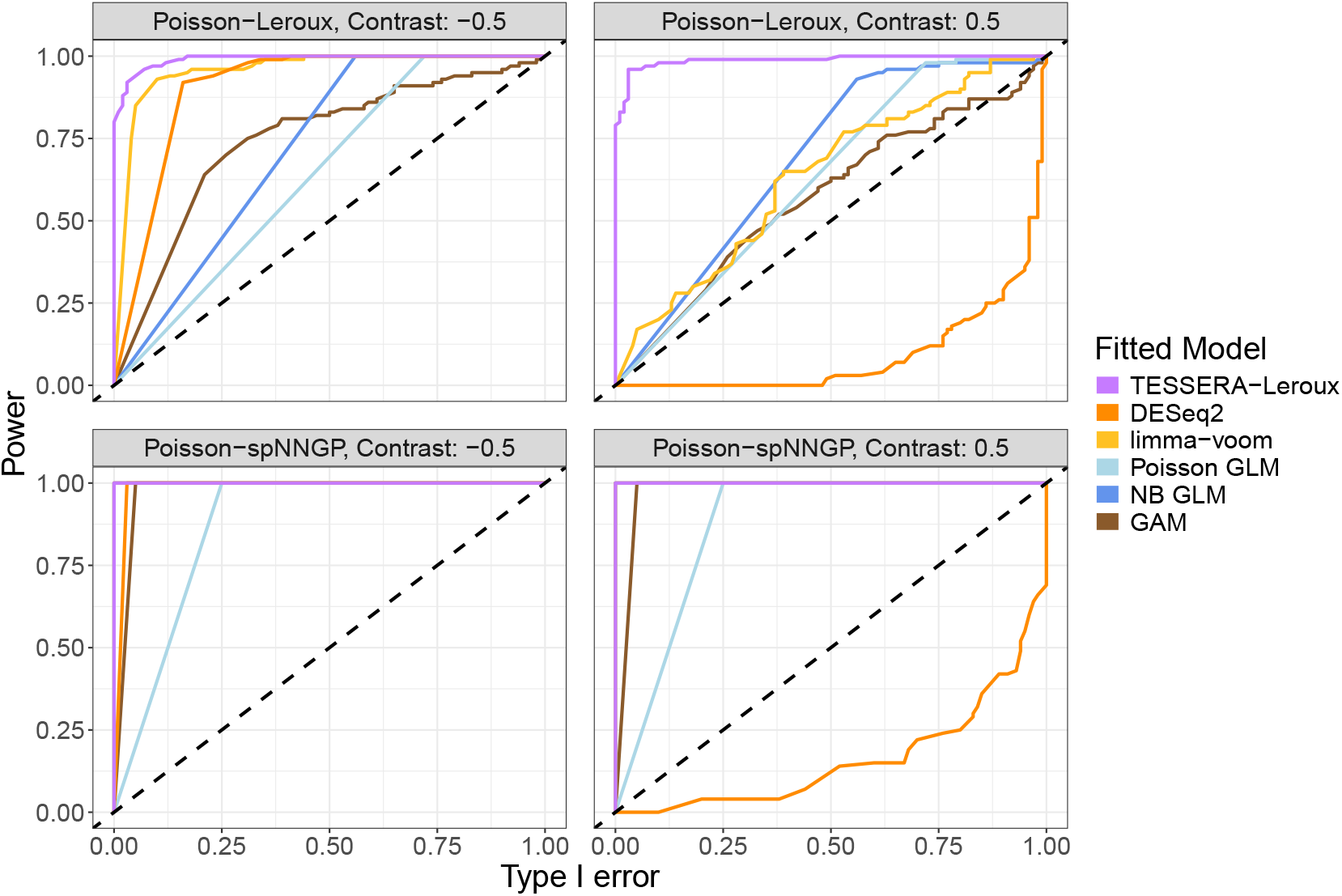
Simulation study: ROC curves. Receiver operating characteristic curves illustrate the trade-off between power (y-axis) and Type I error rate (x-axis) for detecting DE between conditions within a cell type for one gene. The dotted diagonal line represents the *y* = *x* identity line, where the power is exactly equal to the Type I error rate. Curves that rise more steeply toward the top-left corner indicate superior method performance. Different colored lines correspond to different methods, as indicated in the legend. Synthetic data are generated by sampling from either a multi-sample Poisson-Leroux (top row) or Poisson-spNNGP (bottom row) distribution with parameters derived from fitting the model to the *IGHG2* gene in the kidney dataset from (Abedini et al., 2024), except for entries of ***β*** which are modified to create the contrast values represented in the individual plot headers (see details in Section 3.4). Results are displayed for the *C_TAL* cell type and the DKD vs. Control condition comparison.

We can examine the overall behavior demonstrated in the ROC curves more closely by considering the power and Type I error explicitly (Figure 4). The right-hand column of Figure 4 shows the observed Type I error curve in simulation for different *p*-value thresholds. `TESSERA` is the only method that controls the Type I error across both settings. In contrast, the curves for all other methods (except `limma-voom` in the spNNGP setting) lie above the 45-degree line, indicating that they inflate the Type I error and produce more false positives than the reported nominal level.

**Figure 4:**
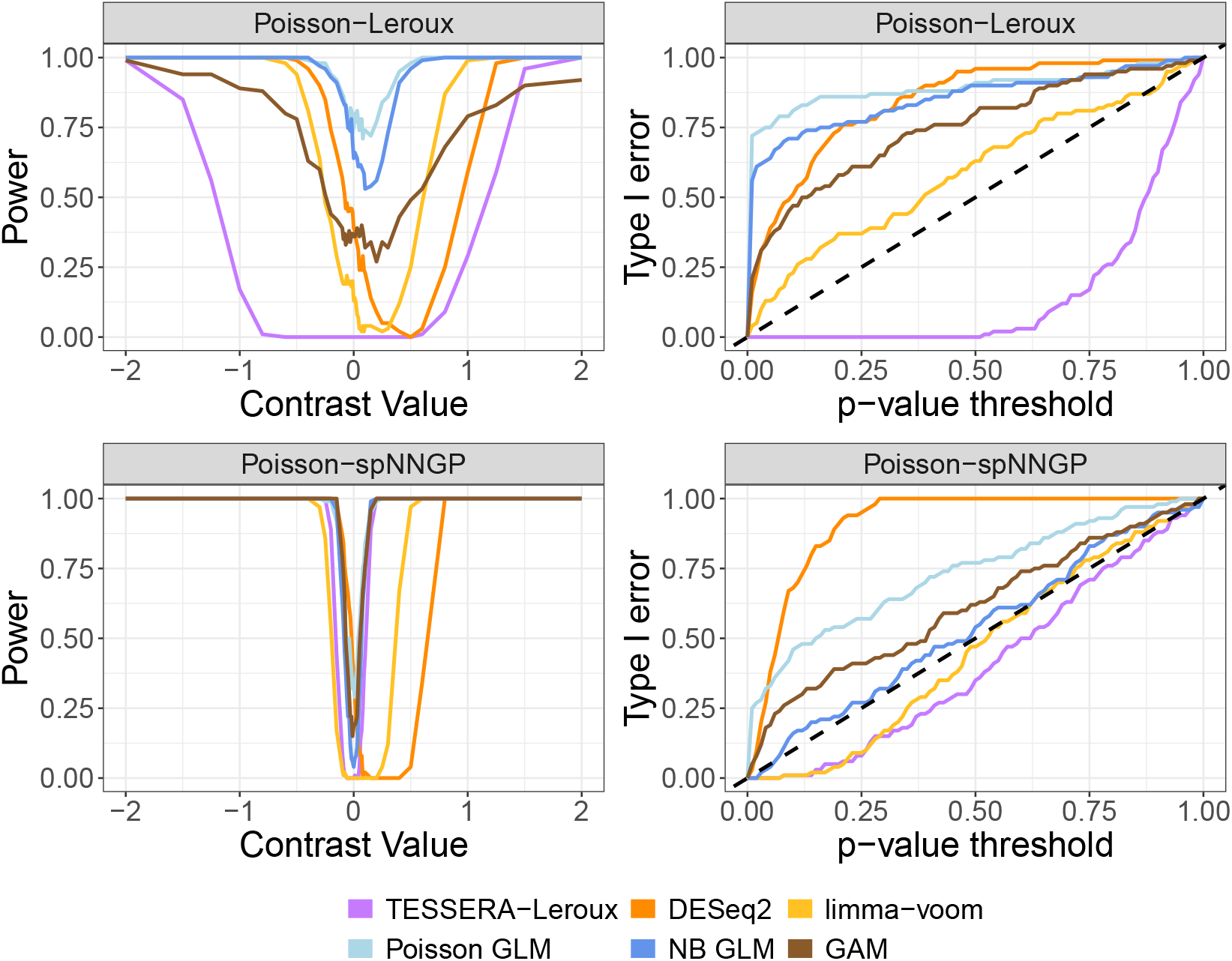
Simulation study: Power and Type I error rate. This figure examines power and Type I error for detecting DE between conditions within a cell type for one gene. In the left-hand column, we present plots of power (y-axis) as a function of the contrast value (x-axis), where power is estimated by the proportion of simulations in which the null hypothesis of no DE is rejected at nominal Type I error rate of 0.05. At an effect size of 0, this value represents the Type I error rate. In the right-hand column, we present curves of the Type I error rate (y-axis) as a function of the *p*-value threshold (x-axis), where exact control of the Type I error rate is represented by the dotted line *y* = *x* and methods with curves above this line do not effectively control the Type I error. Different colored lines correspond to different methods, as indicated in the legend. Synthetic data are generated by sampling from either a multi-sample Poisson-Leroux (top row) or Poisson-spNNGP (bottom row) distribution with parameters derived from fitting the model to the *IGHG2* gene in the kidney dataset from Abedini et al. (2024), except for entries of ***β*** which are modified to create the effect sizes represented on the x-axis for the power curves (see details in Section 3.4). Results are displayed for the *C_TAL* cell type and the DKD vs. Control condition comparison.

Turning to the power analysis, the curve shown in the left-hand column of Figure 4 shows that all methods exhibit the expected V-shaped power curve, where power increases with the magnitude of the effect size (see Supplementary Figures S-15 and S-16 for additional cell types). Of particular note is a distinct rightward shift in the power curves of all methods other than `TESSERA`; while this is observed for all comparison methods in the Leroux setting, it persists for pseudobulk methods `limma` and `DESeq2` across both settings. This shift also helps explain the lack of Type I error control for pseudobulk methods. Notably, when the effect size is zero, the plotted power value represents the empirical Type I error rate, and typically this is also the minimum of the power curve. The pseudobulk methods exhibit curves whose minimum power values are similar to `TESSERA`, but are shifted to the right. Because of this shift, these minima are not located at the point of zero effect size, but at a point where the methods should have power to detect an effect. This shift is also reflected in the behavior of the pseudobulk methods in the ROC curves of Figure 3, which dip below the 45-degree diagonal at small positive contrast values where the Type I error rate exceeds the power to detect the true signal. In both covariance settings, `TESSERA` is the only approach that achieves its power while strictly maintaining the Type I error rate at a nominal level of 0.05.

Beyond the differences between methods, we also observed a discrepancy in absolute power between the two covariance settings. While `TESSERA` has lower power in the Leroux setting than in the misspecified spNNGP setting, this difference is driven by the magnitude of the standard errors of the estimated contrasts. As seen in Figure S-17, the standard errors of the estimated contrasts in the spNNGP setting are less than half of those in the Leroux setting, while the estimated contrast values are similar (though we note that the spNNGP contrast estimates tend to be slightly larger in magnitude). The smaller standard errors lead to notably larger Wald statistics, and hence likely explain the higher power for the spNNGP setting relative to the Leroux setting.

#### 3.4.3. Impact of Cell Abundance

The extent of Type I error inflation is strongly influenced by cell abundance. We examined performance across four cell types ranging from ∼40 to ∼600 average cells per sample. For the GAM, Poisson GLM, and NB GLM methods, the deviation from the nominal Type I error rate becomes progressively more severe as the number of cells per sample increases (Supplementary Figures S-18 and S-19). In the case of the Poisson and NB GLM, GAM, and `DESeq2`, the Wald statistic is modeled as having a *χ*^2^ distribution, where the standard errors are estimated from the observed information matrix, which corresponds to the Hessian of a misspecified likelihood function that treats the counts as independent. Here, treating the counts as independent leads to standard errors that are systematically underestimated, as evidenced by standard errors that decrease with increasing cell abundances while the nominal Type I error simultaneously worsens (Supplementary Figures S-20 and S-21).

#### 3.4.4. Robustness via Empirical Null Distribution Estimation

While `limma-voom` does not completely control the Type I error rate, its deviations are notably less pronounced than those of the other benchmarked methods. This superior performance likely stems from its structural flexibility; by employing a Gaussian likelihood, `limma-voom` decouples the mean and variance, whereas Poisson and NB models are constrained by rigid, predetermined mean-variance dependencies. This decoupling allows `limma-voom` to more accurately approximate empirical distributions that do not strictly conform to the fixed dispersion structures inherent to Poisson and NB models.

`TESSERA` provides a different path toward robustness by addressing the inherent limitations of relying on a theoretical null distribution. While `limma-voom` and `DESeq2` both share information across genes to stabilize specific model parameters such as variance or dispersion, `TESSERA` instead shares information across genes for the test statistics themselves to characterize the null distribution. By employing a scaled non-central 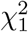 distribution to learn the empirical null distribution directly from the data, `TESSERA` seeks to ensure valid inference even under model misspecification. Consequently, while `limma-voom` provides more flexibility than NB or Poisson-based models, it remains constrained by a theoretical null distribution, a limitation shared by `DESeq2` and the cell-level Poisson and NB-based GLM or GAM models. In contrast, `TESSERA` maintains robust Type I error control by adapting its null distribution to the unique characteristics of the observed test statistics. This empirical approach is essential for accurate Type I error control; as demonstrated in Supplementary Figures S-22 and S-23, applying a standard chi-squared null distribution with one degree of freedom fails to maintain the nominal Type I error rate.

We compared `TESSERA` against `fdrtool` (Strimmer, 2008), a widely-used implementation of the empirical null framework that builds upon the local FDR approach established by Efron (2004). Notably, `TESSERA`’s estimation procedure (Section 2.4) does not require the null distribution to be centered or symmetric, providing a more versatile framework than methods like `fdrtool`, which rely on such assumptions. While performance on the simulated data is virtually identical (Supplementary Figures S-22, S-23, S-24, and S-25), `TESSERA` generalizes to settings where skewed or shifted null distributions would otherwise violate the symmetry and centering assumptions of conventional empirical models. As in our application of `TESSERA`’s thresholding procedure for the null distribution, we applied `fdrtool` to a single realization (the real data with 3,000 genes).

#### 3.4.5. Robustness to Spatial Covariance Structure

Interestingly, non-spatial models show higher Type I error inflation in the Leroux data-generating setting than in the spNNGP setting. The Leroux model generates local, discrete clusters of spatial dependency (Cramb et al., 2020; Wall, 2004), whereas the spNNGP model produces global, smooth gradients characteristic of the Matérn covariance family (Datta et al., 2016). The Leroux localized structure represents a more severe departure from the independence assumption, sharply reducing the effective sample size (Griffith, 2005) and making these “pockets” of expression more likely to be misinterpreted as biological signal by non-spatial models. `TESSERA` is less prone to spurious discoveries as it partitions the total variance into components related to the fixed effects and to a structured spatial random effect, thereby preventing random spatial autocorrelation from being misattributed to the contrast of interest.

## 4. Real Data Analysis: Visium Kidney Tissue Samples

We analyze a spatial transcriptomics dataset generated using the Visium platform on formalin-fixed, paraffin-embedded (FFPE) human kidney tissue samples (Abedini et al., 2024). The dataset comprises 14 samples from distinct patients, representing three conditions: three healthy controls, six patients with diabetic kidney disease (DKD), and five patients with hypertensive kidney disease (HKD). Cell type annotations were provided by the original authors, using `Cell2Location` (Kleshchevnikov et al., 2022) and `CellTrek` (Wei et al., 2022) to identify 26 distinct cell types, with varying numbers of cells per cell type–disease combination (Supplementary Figure S-26). We further filtered the dataset to retain the most highly variable genes and computed library sizes to serve as normalization factors in our analyses (see Section S1.1).

For all analyses, we applied the `TESSERA` method to fit a multi-sample Poisson-Leroux model and compared its results with a Poisson GLM, a negative binomial GLM, a generalized additive model, and pseudobulk approaches (`DESeq2` and `limma-voom`).

All models were fit using a fully-crossed design between cell type and disease and include a fixed effect for sample (see Section S1.2 for details). Differential expression analyses were performed both between cell types and between conditions within an individual cell type. For the `TESSERA` model, we obtained *p*-values using our Wald test thresholding procedure (Section 2.4). For each method and analysis, we computed adjusted *p*-values using the Benjamini-Hochberg (BH) procedure (Benjamini and Hochberg, 1995) for control of the false discovery rate (FDR) over genes.

### 4.1. Model Fit

The results of applying `TESSERA` on this dataset demonstrate several important advantages of the underlying multi-sample spatial model. Firstly, in most settings, at least some cell types will be highly spatially clustered while also having quite differing gene expression levels; not including them in the model (via the design matrix in the model of Equation (3)) can result in estimates of spatial variation which are actually due to cell type distinctions (Walker et al., 2022; Yu and Luo, 2022; Gaspard-Boulinc et al., 2025; Shang, Wu and Zhou, 2025). Our model allows for the separation of cell type effects from spatial effects. The gene expression within each sample can be decomposed into estimated covariate (e.g., cell type) effects 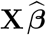 and estimated spatial random effects 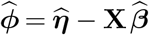, where 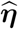 is estimated using the approximation in Equation (19) and the ECM algorithm. When we calculate the Moran’s *I* of 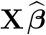 compared to the Moran’s *I* of 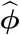, we see high values for 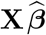 (Figure 5), indicating the importance of including cell type covariates to avoid spuriously categorizing cell type spatial patterning as autocorrelation. We also see that 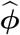 retains substantial spatial autocorrelation, distinct from the notably higher signal found in the covariate component. This confirms that while the spatial distribution of cell types accounts for much of the observed spatial structure, our model successfully identifies an underlying spatial dependence.

**Figure 5:**
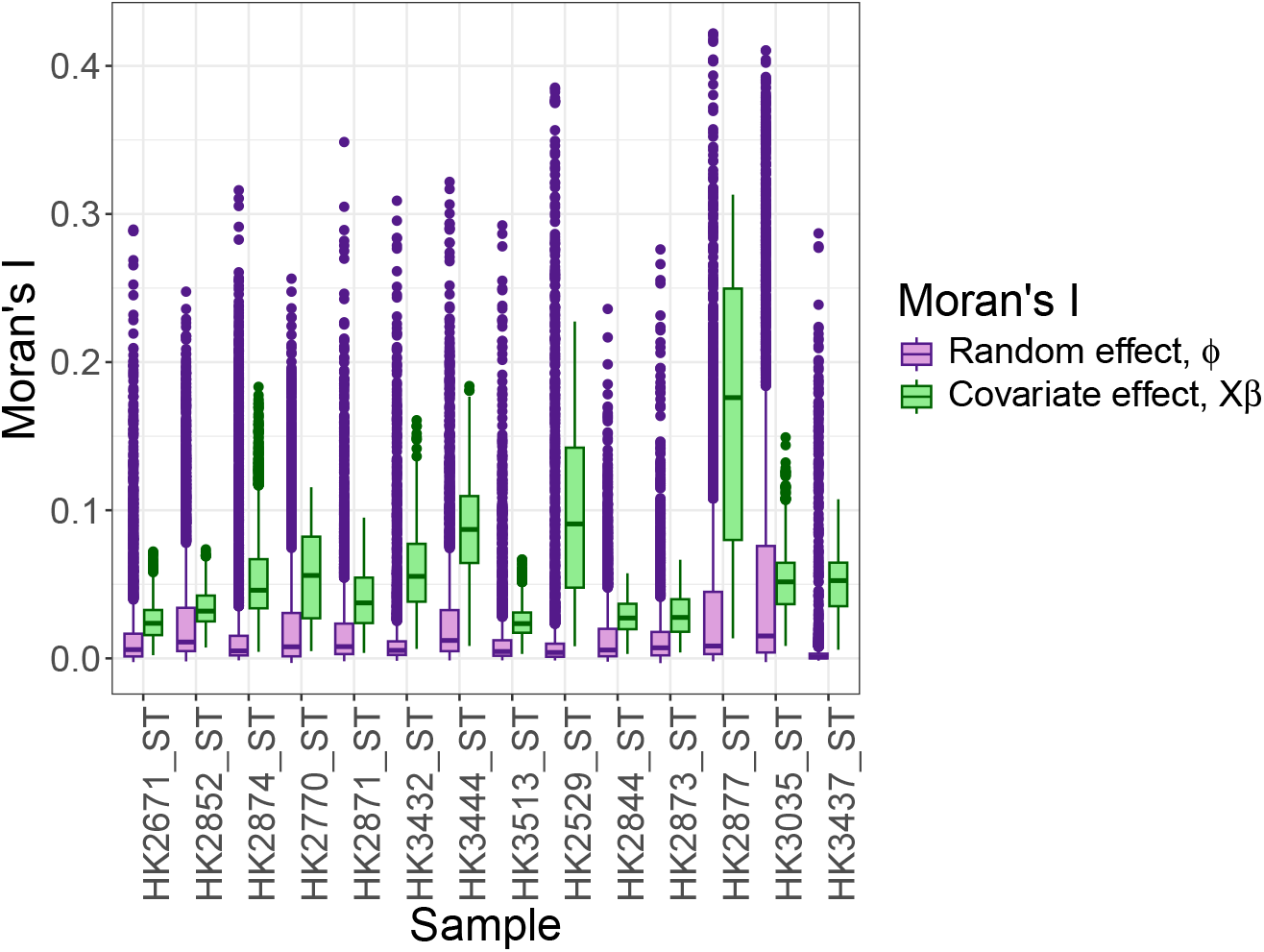
Kidney dataset: Moran’s *I* of spatial random effect and covariate effect. Box-plots show, per sample, the distribution across genes of Moran’s *I* calculated on the estimated spatial random effects 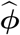 (purple) and covariate effects 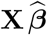 (green) (see Section 4 for details). Results are based on fitting a multi-sample Poisson-Leroux model with `TESSERA` to the kidney dataset (Abedini et al., 2024). The 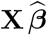 have large values of Moran’s *I*, demonstrating that the covariates induce significant spatial patterning, but the estimated random effects also capture additional, non-trivial spatial autocorrelation.

Moreover, we find that `TESSERA` fit with the Poisson–Leroux model generally yields smaller residuals, as measured by the normalized mean squared error, compared with the other methods (Supplementary Figure S-27). Compared with the Poisson and NB GLM, the `TESSERA` residuals exhibit a substantially stronger decrease in their spatial autocorrelation, as measured by Moran’s *I*, compared to the original counts (Supplementary Figure S-28). Given that on synthetically generated non-spatial, over-dispersed count data the decrease in Moran’s *I* is comparable for `TESSERA` and the Poisson and NB GLM (Supplementary Figure S-29), our results here indicate that a non-spatial model does not adequately explain all the variation inherent to the data.

Beyond the need for a spatial model, we note that the estimated spatial parameters *γ* and *τ* ^2^ vary widely across samples (Supplementary Figure S-30), demonstrating that there is substantial heterogeneity in the spatial correlation structure across samples necessitating separate spatial parameters per sample.

### 4.2. Spatial Modeling Recovers Localized Anatomical Structure

To further examine the behavior of `TESSERA` against a non-spatial model, we focus on the gene *ACTA2*, a canonical marker for vascular smooth muscle cells (*VSMC*), in sample *HK3035_ST*. This sample contains a prominent *VSMC* cluster where *ACTA2* exhibits a notable degree of spatial structure in 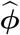, summarized by a Moran’s *I* value of 0.19. This choice of gene-sample pair illustrates `TESSERA`’s ability to identify a strong, spatially organized signal that is distinct from the cell type fixed effects.

The `TESSERA` method provides a superior fit for the vascular smooth muscle marker *ACTA2* for this sample compared to a non-spatial NB GLM. Both models include fixed effects for cell types (Figure 6a) to establish a baseline for gene expression; however, while the NB GLM produces residuals that cluster heavily along a renal artery (Figure 6h), `TESSERA` effectively reduces this structured error through a Poisson-Leroux spatial random effect 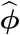 (Figures 6g and 6e). The spatial patterns in the NB GLM residuals stem from the assumption of spatial invariance, which forces a single, uniform average on every spot with the same cell type label regardless of its anatomical context. Because this cell type-specific average is calculated globally, it cannot capture the high-intensity signals of major vessels, causing the NB GLM fitted values (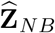, Figure 6d) to appear fragmented and systematically under-estimated compared to the raw counts (Figure 6b). Although the NB GLM uses an over-dispersion parameter to account for this remaining variance, it treats the variation as random rather than structured, leaving the model unable to resolve the biological signal along the tissue’s architecture. By incorporating both a covariate effect 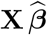 (Figure 6f) and Poisson-Leroux spatial random effect 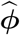 (Figures 6e), `TESSERA` more effectively approximates the architecture of the kidney’s arterial network in the final fitted counts (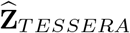, Figure 6c). Consequently, the `TESSERA` fitted counts are closer to the observed raw counts and are more biologically consistent with the known physical architecture of the kidney.

**Figure 6:**
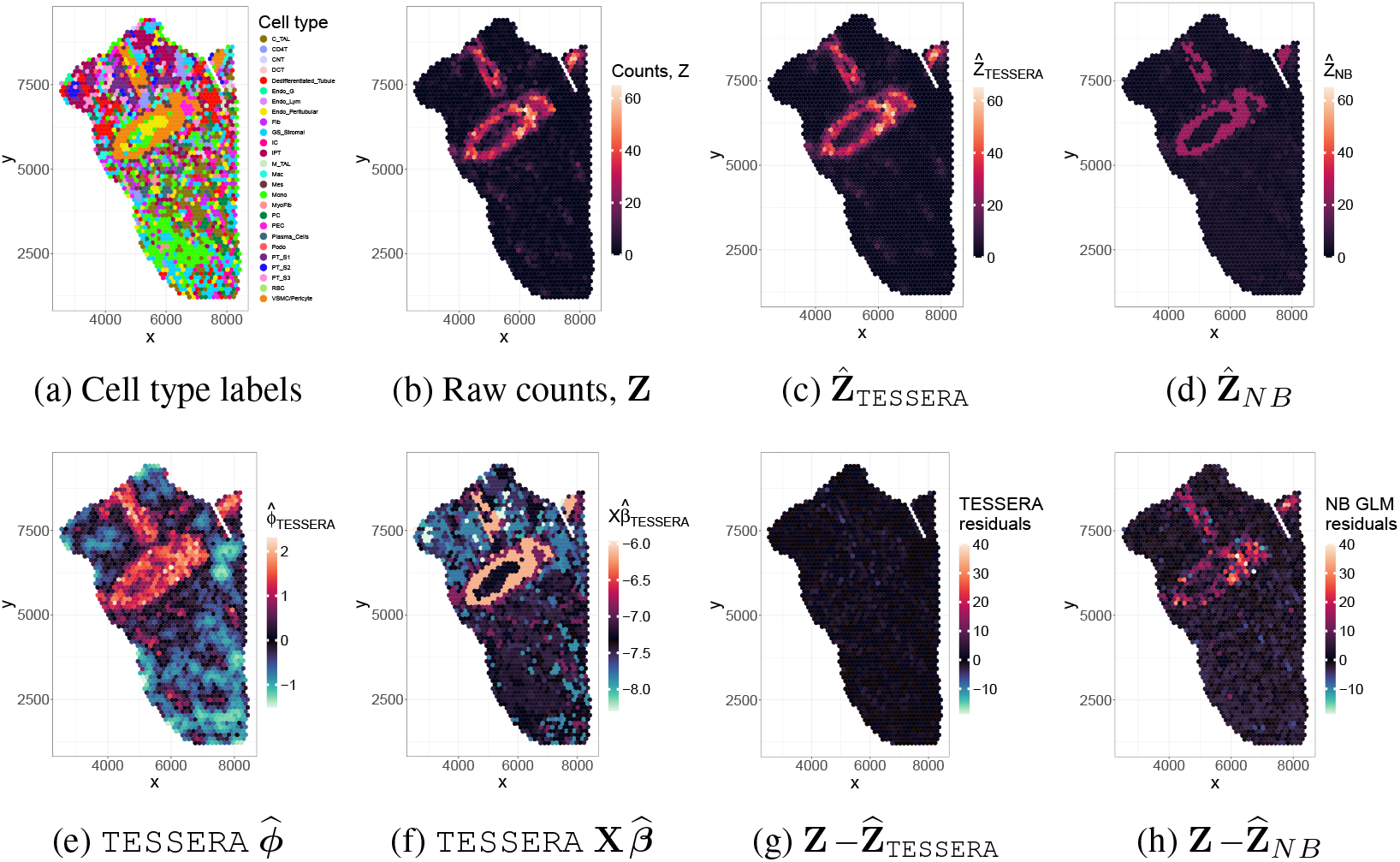
Kidney dataset: Capturing structured vascular expression. For the kidney dataset (Abedini et al., 2024), we examine the results of model fitting for both `TESSERA` with a Poisson-Leroux model and an NB GLM on the gene *ACTA2*. We focus on the results for sample *HK3035_ST*. **(a)** Categorical cell type annotations used as fixed effects in both models. (b) Observed raw counts, **Z**, for the vascular marker *ACTA2*. **(c)** `TESSERA` fitted counts, 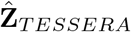. **(d)** NB GLM fitted counts, 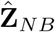 . **(e)** `TESSERA` estimated spatial random effects, 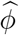. **(f)** `TESSERA` estimated covariate (cell type) effects, 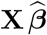. **(g)** `TESSERA` raw residuals, 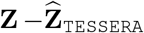. **(h)** NB GLM raw residuals, 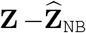.

### 4.3. Differential Expression Between Cell Type

Next, we examine whether applying `TESSERA` to identify genes that are DE between cell types yields known marker genes that are predominantly expressed in specific cell types and serve to characterize them or distinguish between them and other cell types. We use the kidney marker genes outlined in Lake et al. (2023, Supp. Table 5) mapped to the cell types present in Abedini et al. (2024) (Supplementary Figure S-31) and focus on the comparison of proximal tubule segment 1 (*PT_S1*) cells and cortical thick ascending limb (*C_TAL*) cells. The marker genes are specific to their respective designated cell type (Supplementary Figure S-32), making this a reasonable comparison.

Formally, we consider the following contrast

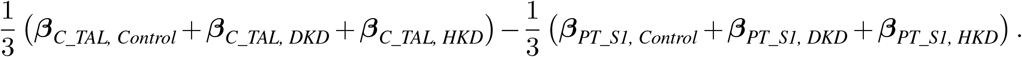

We test for DE between the two cell types using the Wald test of Section 2.4, where the parameters of the null distribution are estimated using all genes across all pairwise cell type contrasts (i.e., based on 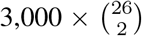 test statistics), and begin by ranking genes according to their Wald test statistics (Supplementary Figure S-33). We can see that all methods rank the marker genes relatively similarly and with a generally low rank, a strong indication of differential expression, as we would expect.

We next create volcano plots for each method (Supplementary Figure S-34), where we see that the marker genes have higher estimated effects and lower BH adjusted *p*-values than non-marker genes across all methods. If we then compare the number of genes that were found to be significantly DE (at a nominal FDR level of 0.05), we find that `TESSERA` fit with the multi-sample Poisson-Leroux model detects similar numbers of genes as the pseudobulk methods and less than half the number of genes as found by the GLM and the GAM (Supplementary Figure S-35). The simulation results in Section 3.4 show that these non-spatial methods fail to control the Type I error, therefore implying that most of their discoveries are likely false positives; indeed, the data exhibit significant spatial autocorrelation, making non-spatial models prone to inflated statistical significance and hence potential false discoveries.

Lastly, we examine the overlap between methods in the ranking of all genes, not just marker genes, and observe high Jaccard similarity across methods for the top-ranked genes (Supplementary Figure S-36). This consistency is expected given the distinct functional roles of proximal tubule and cortical thick ascending limb cells. Due to the pronounced differences in gene expression between these cell types, the primary markers are easily captured by all methods, leading to a high degree of consensus in the resulting rankings.

### 4.4. Differential Expression Between Conditions, Within Cell Types

We next study the more biologically interesting scenario of differential expression between conditions within a cell type, where, for cell type *c*, the contrasts of interest are

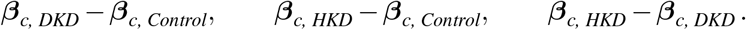

We examine the number of genes found to be DE (at a nominal FDR level of 0.05) by our `TESSERA` method, using the procedure described in Section 2.4, where the parameters of the null distribution are estimated using the Wald statistics from all within-cell type pairwise contrasts between conditions (Control, DKD, and HKD) for all genes (i.e., 3,000 ×3 ×26 test statistics). Diagnostics of the Wald test statistics for the `TESSERA` method indicate that our thresholding procedure identifies a single optimal threshold that minimizes the *L*^2^ error. Applying the resulting threshold produces a *p*-value distribution for sub-threshold statistics that closely follows a 𝒰 (0, 1) distribution (Supplementary Figure S-37), with the corresponding scale and shift parameters for these test statistics shown in Supplementary Figure S-38.

We compare the number of differentially expressed genes identified by `TESSERA` to those found by Poisson GLM, NB GLM, GAM, and pseudobulk methods `limma-voom` (Ritchie et al., 2015) and `DESeq2` (Love, Huber and Anders, 2014). All DE methods are consistent in that many more genes are identified as DE between DKD and the other two conditions than between Control and HKD, as seen in Supplementary Figure S-39. This result is consistent with previous work and with the underlying pathology of DKD and HKD (Barrios et al., 2025). However, the number of genes found DE for a particular comparison varies considerably from method to method as seen in Figure 7 and Supplementary Figure S-40.

**Figure 7:**
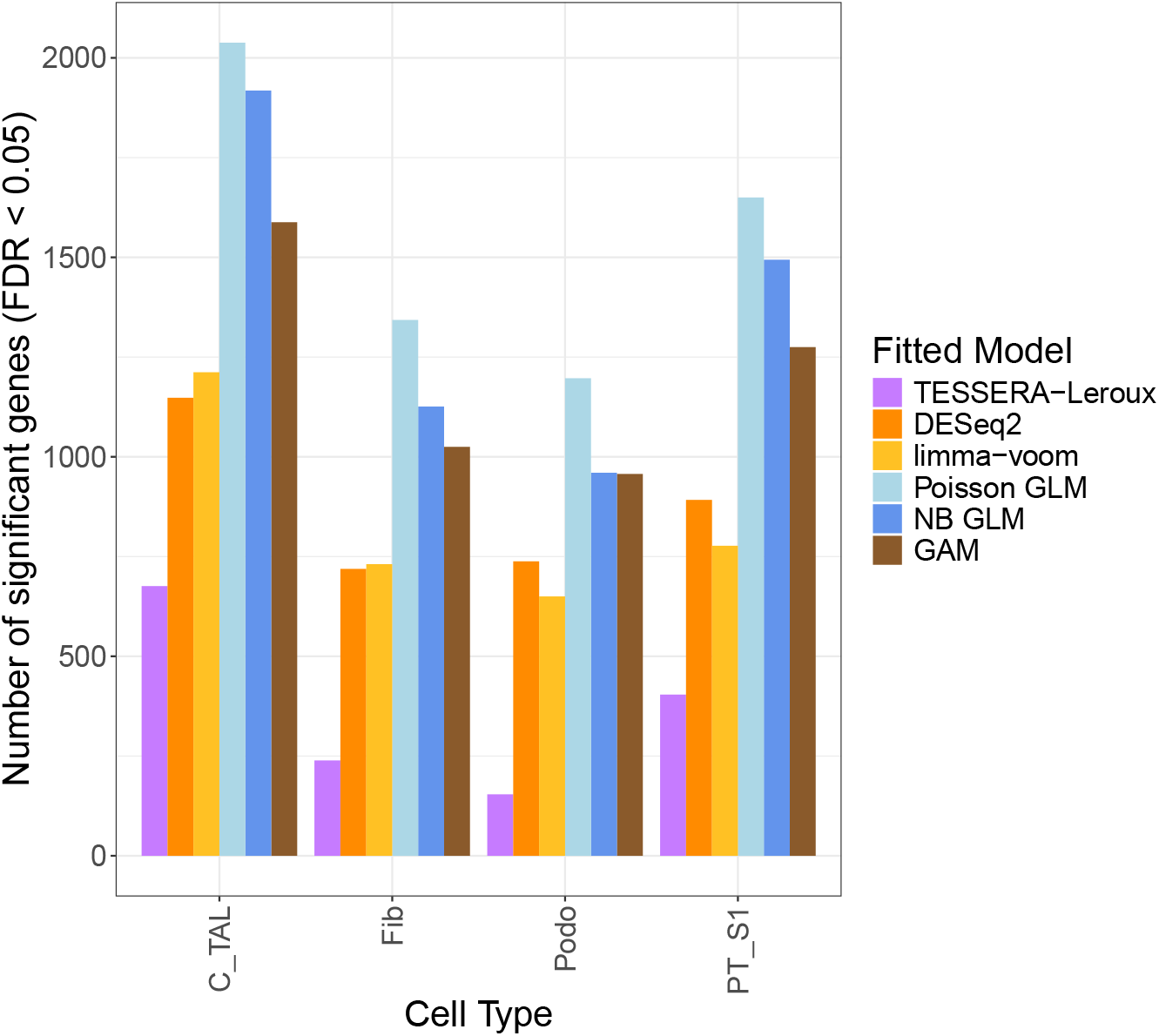
Kidney dataset: Number of differentially expressed genes for the Control vs. DKD contrast. The height of each bar indicates the number of genes found to be differentially expressed between the Control and DKD conditions in four cell types representing different structured compartments of the kidney, for the dataset of Abedini et al. (2024): Cortical thick ascending limb (*C_TAL*), fibroblasts (*Fib*), podocytes (*Podo*), and proximal tubule segment 1 (*PT_S1*). For all methods, significance is based on Benjamini-Hochberg-adjusted *p*-values with a nominal FDR level of 0.05. For `TESSERA`, *p*-values are derived from the Wald test procedure described in Section 2.4. Different colors correspond to the results from the different DE methods (see legend).

Specifically, in Figure 7, `TESSERA` identifies the fewest DE genes among the methods compared, followed by the pseudobulk approaches. Notably, the GAM and GLM methods identify more than twice as many DE genes as `TESSERA`, a finding consistent with the results seen in the *C_TAL* vs. *PT_S1* comparison (Supplementary Figure S-35). This suggests that accounting for spatial correlation in our model reduces our effective sample size appropriately and, taken together with the results on Type I error in Section 3.4, demonstrates that accounting for spatial correlation leads to fewer spurious discoveries. Furthermore, `TESSERA` exhibits higher Jaccard similarity for top-ranked genes with pseudobulk methods, which treat the sample as the experimental unit, than with GLM or GAM methods, which treat cells as independent observations (Supplementary Figure S-41).

## 5. Discussion

In this paper, motivated by recent advances in spatial transcriptomics technology and its growing adoption, we have proposed the `TESSERA` method for analyzing large, multi-sample datasets. Key elements of our method are that it is based on a Poisson model that operates directly on raw counts, that it allows for distinct spatial covariance structures in each sample while estimating a single set of fixed effects across samples, and that it is based on a scalable ECM algorithmic framework. Additionally, we are able to present identifiability results for the fixed effects ***β*** as well as for the spatial covariance parameters in lattice models.

We have demonstrated the performance of `TESSERA` on realistically generated synthetic data. We showed that it accurately estimates the underlying fixed effects and the spatial covariance parameters of the lattice models, and that it accurately models the counts (measurements). Compared to non-spatial methods, our method leads to a substantial reduction in the spatial autocorrelation of the fitted residuals. Furthermore, `TESSERA` enables robust statistical inference, achieving superior or comparable power at any given Type I error rate across a broad range of effect sizes compared to simpler GLM and GAM alternatives and standard pseudobulk methods. Crucially, out of all compared methods, only ours was able to consistently control the Type I error rate regardless of the underlying spatial generative process. While non-spatial methods showed worsening Type I error control as cell counts increased and pseudobulk methods like `limma` and `DESeq2` exhibited biased power curves, `TESSERA` provided valid inference across all settings. These results underscore the necessity of models that both account for spatial autocorrelation and accommodate heterogeneity in spatial structures across samples.

We applied our method to a real spatial transcriptomics dataset, generated by the authors in Abedini et al. (2024). We find that our method finds far fewer differentially expressed genes than all other methods, both for comparisons between cell types and comparisons between conditions for a given cell type. Taken together with the power and Type I error results on the synthetic data, we conclude that it is likely that a substantial portion of the genes discovered by other methods are false discoveries.

Within spatial transciptomics, we believe that the `TESSERA` method is an extremely useful analytic framework that avoids the pitfalls incurred by not accounting for the spatial correlation between neighboring cells. Notably, our approach provides a hitherto nonexistent extension to the multi-sample setting, while remaining competitive with or outperforming existing algorithms in the single-sample setting, thereby filling a critical gap in current methodological capabilities. Moreover, compared to GAMs or existing spatial computational frameworks such as MCMC samplers and INLA, our ECM framework enables faster model fitting and inference under the increasing computational demands of multi-sample settings with numerous experimental units and spatial locations. While we have focused on spatial transcriptomics applications, we expect that `TESSERA` has application beyond genomics, e.g., in urban planning, ecology, or for geospatial analyses involving islands or other disconnected structures.

## Acknowledgments

We thank Iain Carmichael for helpful conversations during early stages of this project. Sandrine Dudoit and Elizabeth Purdom are joint corresponding authors.

## SUPPLEMENTARY MATERIAL

### S1. Model Implementation

This section provides implementation details for the application of `TESSERA` and other methods to both simulated and real data in Sections 3 and 4, respectively. Section S1.1 outlines the data preprocessing steps applied to the spatial transcriptomics kidney dataset from Abedini et al. (2024). Section S1.2 describes the design matrix which encodes the fully-crossed effects of cell type and disease along with sample-specific fixed effects, and which is used for all methods. Section S1.3 then summarizes the computational settings, preprocessing steps, and software used for both single-sample and multi-sample analyses. Table 2 provides a complete overview of the models, input data, and key algorithmic parameters.

**Table 2:**
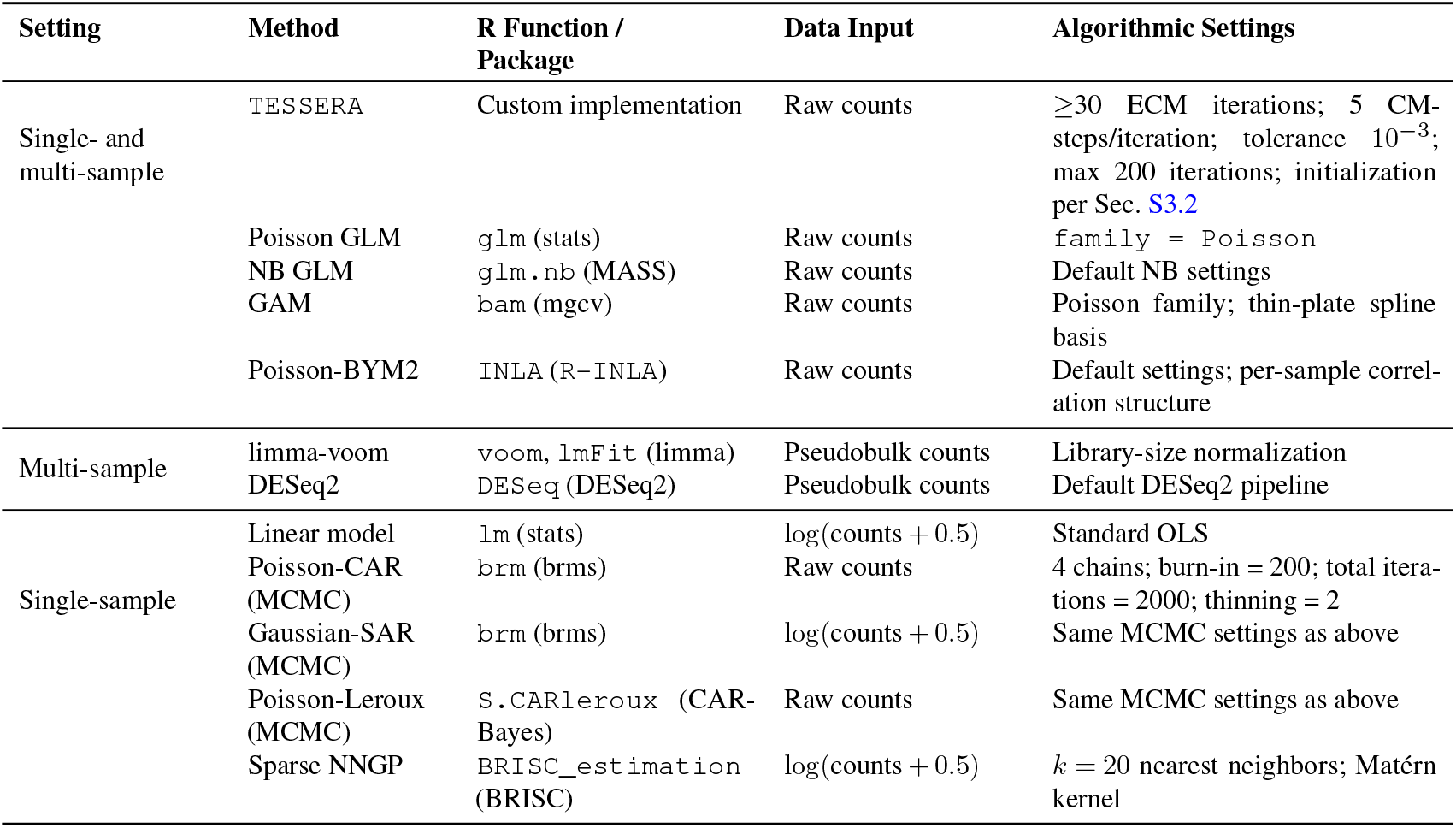
Methods evaluated and implementation specifications.

| Setting | Method | R Function / Package | Data Input | Algorithmic Settings |
| --- | --- | --- | --- | --- |
| Single- and multi-sample | TESSERA | Custom implementation | Raw counts | $\geq 30$ ECM iterations; 5 CM-steps/iteration; tolerance $10^{-3}$; max 200 iterations; initialization per Sec. S3.2 |
|  | Poisson GLM | glm (stats) | Raw counts | family = Poisson |
|  | NB GLM | glm.nb (MASS) | Raw counts | Default NB settings |
|  | GAM | bam (mgcv) | Raw counts | Poisson family; thin-plate spline basis |
|  | Poisson-BYM2 | INLA (R-INLA) | Raw counts | Default settings; per-sample correlation structure |
| Multi-sample | limma-voom | voom, lmFit (limma) | Pseudobulk counts | Library-size normalization |
|  | DESeq2 | DESeq (DESeq2) | Pseudobulk counts | Default DESeq2 pipeline |
| Single-sample | Linear model | lm (stats) | $\log(\text{counts} + 0.5)$ | Standard OLS |
|  | Poisson-CAR (MCMC) | brm (brms) | Raw counts | 4 chains; burn-in = 200; total iterations = 2000; thinning = 2 |
| | Gaussian-SAR (MCMC) | brm (brms) | $\log(\text{counts} + 0.5)$ | Same MCMC settings as above |
|  | Poisson-Leroux (MCMC) | S.CARleroux (CAR-Bayes) | Raw counts | Same MCMC settings as above |
| | Sparse NNGP | BRISC_estimation (BRISC) | $\log(\text{counts} + 0.5)$ | $k = 20$ nearest neighbors; Matérn kernel |

#### S1.1. Data Preprocessing

The dataset from Abedini et al. (2024) was preprocessed by the authors to remove low-quality spots and low-expression genes, resulting in 12,511 genes measured across 37,143 cells. To account for variation in sequencing depth across cells, we computed a library size for each cell as the sum of its counts across all genes. This quantity, *C*_*i,j*_, serves as a standard proxy for sequencing depth and was included as a normalization factor in all downstream models. Following common practice in high-dimensional gene expression analyses, to reduce noise and improve power, we retained only the most variable genes. Inspection of the variance–rank curve for log_10_-transformed gene variances (Supplementary Figure S-42) revealed a clear inflection point at approximately 3,000 genes, and we therefore restricted our analyses to this subset of genes.

The spNNGP model depends on pairwise distances between cells and the lattice models on cell-cell adjacency matrices. For the spNNGP model, pairwise Euclidean distance matrices are calculated between the cell centroids, i.e., their spatial coordinates. To derive cell-cell adjacency matrices for the lattice models, we threshold the pairwise distances between cell centroids. In particular, an analysis of these pairwise distances revealed that, for the vast majority of cells, there is a large gap between the sixth-smallest and seventh-smallest distance values. This gap in distances corresponds to the hexagonal pattern in the 10x Genomics Visium platform, so that we expect all cells that are not on the edge of the tissue to have exactly six neighbors (Crowell et al., 2025; 10x Genomics, 2025b). Were we given data from an imaging platform (e.g., CosMx SMI), we would instead use the results of the cell segmentation on the tissue images and define neighboring cells as those whose minimum distance between boundaries is under some threshold or whose boundaries are in contact.

#### S1.2. Model Design Matrix

For both the real and synthetic data analyses, we use a fully-crossed design between cell type and disease for all models. The design includes no intercept and no cell type and no disease main effects, but does include a fixed effect for sample. That is, the design matrix consists of binary indicators, with one column for each possible combination of cell type and disease (e.g., *C_TAL* cells in a Control sample) and one column for each sample. Concretely, our model has the form

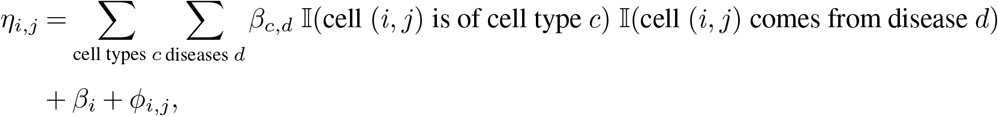

where 𝕀 is the indicator function. The corresponding design matrix has full column rank and no zero rows.

An equivalent model is a properly-constrained model that includes an intercept and main effects

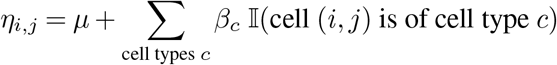

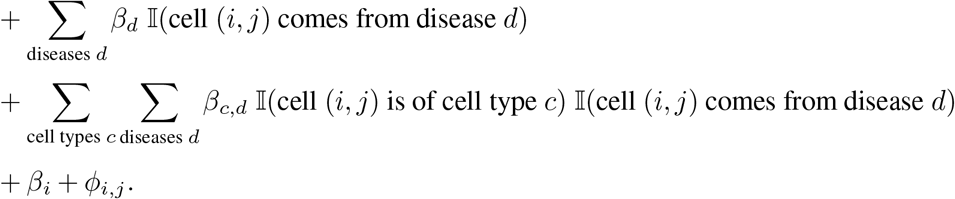

It is important to note that, for identifiability, either some constraints must be imposed on the parameters *β* (e.g., sum-to-zero) or the summations must not range over all levels. That is, there must be a reference level for each of the factors. To see that the two models are equivalent, note that every cell must belong to some combination of disease and cell type and both design matrices are capable of uniquely representing all such combinations. Thus the design matrices of both the intercept-and-main-effects model and the interaction-only model are simply different linear reparameterizations of one another. That is, any group mean that can be expressed as a sum of a baseline and offsets can also be expressed directly as a standalone mean. It follows that the design matrices have the same range and that the models are equivalent.

The advantage of the intercept-free model is that the contrasts of interest, differences in cell types within a disease or differences between diseases for a given cell type, are simply differences of two coefficients, whereas in the model with main effects and an intercept, the formulation is slightly more complex.

For our analysis of the kidney dataset, the column corresponding to the Endo-Lym cell type interaction with the HKD condition is dropped, as there are no Endo-Lym cells in the HKD samples.

#### S1.3. Implementation Details

We implemented a range of models to assess both spatial and non-spatial approaches in single-sample and multi-sample settings. Alongside the `TESSERA` method, these include standard Poisson and negative binomial GLM, generalized additive models (GAM), pseudobulk methods (`DESeq2` and `limma-voom`), as well as spatial models including CAR, Gaussian-SAR, Leroux, and sparse NNGP.

For the Poisson and negative binomial models, raw counts were analyzed directly. For linear and Gaussian-based models, counts were log-transformed with a 0.5 offset to handle zero values. For pseudobulk methods, counts were aggregated for each cell type-sample combination. MCMC-based methods were run with four chains, 200 burn-in iterations, 2,000 total iterations, and thinning by 2. The sparse NNGP model used *k* = 15 nearest neighbors and an exponential covariance kernel.

All analyses were performed in `R` (version 4.5.1). Table 2 summarizes the models and methods, software functions, input data, and key algorithmic settings for both single- and multi-sample methods.

### S2. Covariance Model Specification

In this section, we discuss the details of the models used by `TESSERA` to define the covariance matrix **∑**_*i*_ of observations within a given sample *i* (i.e., the specification of the random effect distribution). Each **∑**_*i*_ is defined by a set of parameters ***ψ***_*i*_ which are specific to each sample, i.e., not shared across samples. For notational simplicity, in the remainder of this section, we suppress the sample index *i*, with the understanding that all parameters introduced should be considered sample-specific.

#### S2.1. Lattice Spatial Random Effect Specification: SAR, CAR, and Leroux

For the lattice models, there are two parameters that make up ***ψ, ψ*** = (*τ* ^2^, *γ*), and we can express the precision matrix of the random effect ***ϕ*** (and thus of ***η***) as **P**(***ψ***) = **P**^∗^(*γ*)*/τ* ^2^, where **P**^∗^ depends on a known adjacency matrix and a typically unknown attraction or correlation parameter *γ*, with *γ ≈* (™ 1, 1) for the SAR and CAR models and *γ ∈* [0, 1) for the Leroux model (Table 1). We use the same notation *γ* in each model to emphasize the parameter’s role in defining the level of spatial attraction between neighboring observations, but it should be recognized that *γ* has different meanings in the different models and thus values of *γ* from different models cannot be directly compared.

Let **W ≈** {0, 1} ^*J×J*^ denote the known measurement-site (cell-cell) adjacency matrix of the *J* observations (cells) within a sample. That is, *W*_*j,j*_*′* = *W*_*j*_*′*_,*j*_ = 1 if locations *j* and *j*^*′*^ are adjacent and 0 otherwise; the diagonals are also set to zero. As a technical condition, we require **W** to have no isolated points; connecting an isolated location to its nearest location solves this problem. Moreover, for the SAR and CAR models, a fully-connected adjacency matrix avoids a non-invertible precision matrix.

Define **D ∈** ℕ^*J×J*^ to be the diagonal degree matrix, where *D*_*j,j*_ is the number of neighbors that location *j* has. Since **W** has no isolated points, **D** is invertible. Furthermore, a given location likely has at most a handful of neighbors (or nearby locations deemed to be neighbors), so that **W** is mostly zero. Thus, importantly, **W** and **D** can be stored and treated as sparse matrices.

##### S2.1.1. SAR Model

Analogous to an autoregressive process in time series data analysis, the spatial autoregressive (SAR) model for *ϕ*_*j*_, *j* = 1, …, *J*, is:

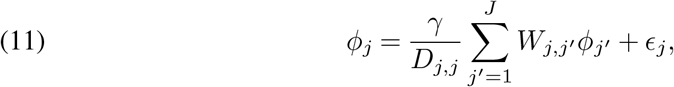

where *ϵ*_*j*_ *∼* 𝒩 (0, *τ* ^2^*/D*_*j,j*_) (Besag and Kooperberg, 1995; Besag, 1974). It follows that the vector ***ϕ*** *∈* ℝ^*J*^ of the *ϕ*_*j*_ satisfies

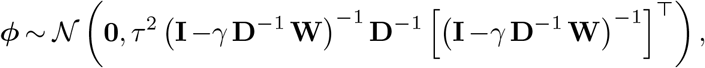

or that the precision matrix **P** *≈* ℝ^*J×J*^ satisfies

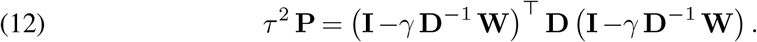

To see this, we note that

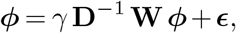

so that

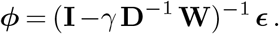

For **P**^−1^ to be a proper covariance matrix, we require *τ* ^2^ *>* 0 and *γ* ≠ ±1. Importantly, due to the structure of **P** in the SAR model, **W** is not required to be symmetric for **P** to be symmetric.

##### S2.1.2. CAR Model

The conditional autoregressive (CAR) model for *ϕ*_*j*_, *j* = 1, …, *J*, is:

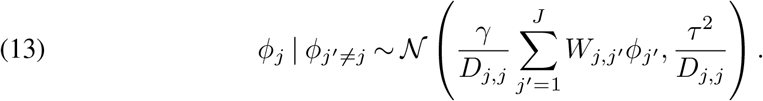

We recall Brook’s Lemma, wherein for a random variable **X ∈** ℝ^*n*^ with two sets of values **x** and **y** and density *f* with conditionals *f*_*j*_, we have that (Brook, 1964)

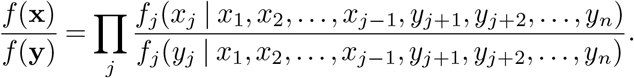

When applying the lemma to the CAR model, take **x** to be the vector ***ϕ*** *∈* ℝ^*J*^ of *ϕ*_*j*_’s and set **y** = **0** *∈* ℝ^*J*^, so that

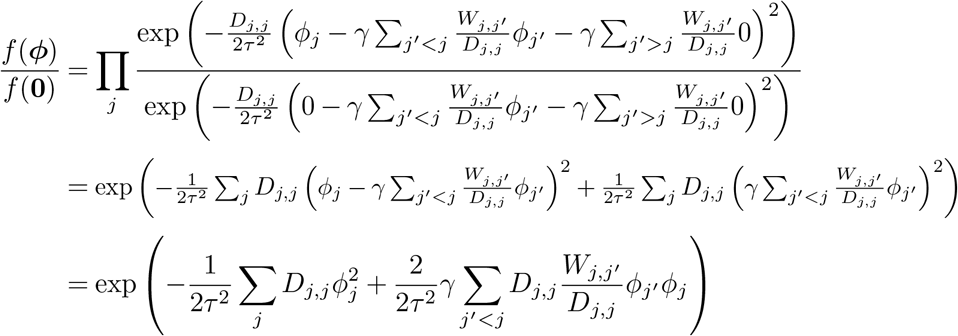

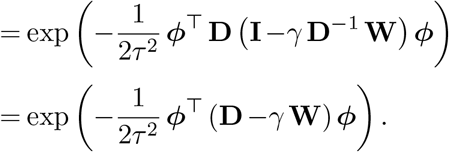

Noting that *f* (**0**) is a constant (i.e., it does not depend on ***ϕ*** but only on the constants **W, D**, *γ*, and *τ* ^2^), we have that *f* (***ϕ***) ∝ *f* (***ϕ***)*/f* (**0**). It follows that, after normalizing appropriately,

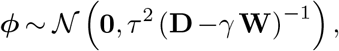

or that (14)

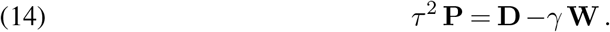

As in the SAR model, for **P**^−1^ to be a proper covariance matrix, we require *τ* ^2^ *>* 0 and *γ* ≠ ±1. Importantly, unlike the SAR model, we require **W** to be symmetric for **P** to be symmetric in the CAR model. We note that the CAR and SAR models are related, in that it is possible to choose a **W**, *γ*, and *τ* ^2^ for one model so that the resulting distribution is the same as that of the other, though practically we usually consider **W** as fixed and determined by the problem (Ver Hoef, Hanks and Hooten, 2018).

##### S2.1.3. Leroux Model

The Leroux model is in the same style as the CAR model, where we have (Leroux, Lei and Breslow, 2000)

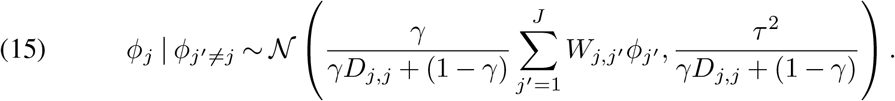

Essentially, relative to the CAR model, the random effect is decomposed as having both a spatially-correlated component as well as a non-spatially-correlated component, wherein *γ ∈* [0, 1) defines the tradeoff between the two: at *γ* = 0, we have independence between the measurements, while as *γ →*1, we recover the intrinsic CAR model (i.e., the improper CAR model with *γ* = 1) (Leroux, Lei and Breslow, 2000). The matrix **P** is then

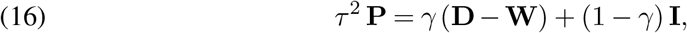

and

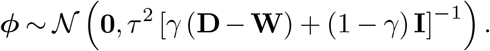

As in the CAR model, for **P**^−1^ to be a proper covariance matrix, we require *τ* ^2^ *>* 0 and *γ* cannot be 1. Importantly, like the CAR model, we require **W** to be symmetric for **P** to be symmetric.

Each of these three models is a natural fit for settings in which neighboring cells might interact or communicate, and cells that are not adjacent are less likely to directly communicate (Song et al., 2019). That is, in a spatial transcriptomics dataset, it is entirely natural to model a cell’s expression as a function of its direct neighbors, or nearby cells, and less so to (directly) include the effects from further-away cells (as a Gaussian or Matérn covariance would entail). That this selection leads to a sparse covariance structure is a boon that will enable the scalability of the model we propose.

#### S2.2. spNNGP Random Effect Specification

For a Gaussian process model, **∑** would be parameterized as **∑**(***ψ***) = **K**(***ψ***), for some kernel matrix **K ∈** ℝ^*J×J*^ that is parameterized by ***ψ***. Often, the structure of **K** can be expressed as

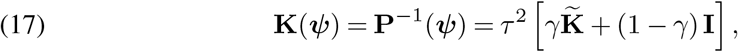

where *τ* ^2^ *>* 0 is an overall measure of scale (analogous to the lattice models) and *γ* trades off between the spatial and non-spatial components of the covariance (exactly analogous to the Leroux model). In this formulation, *γτ* ^2^ and (1 − *γ*)*τ* ^2^ correspond to the spatial variance (partial sill) and nugget parameters, respectively. The matrix 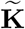 captures the pairwise relationship between locations. If we denote the measurement locations (coordinates) as **s**_*j*_, for isotropic kernels like we consider herein, the (*j, j*^*′*^) entry of 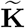 and hence **K**(***ψ***) are functions of ∥**s**_j_ ™ **s**_*j*_*′* ∥_2_. Beyond the measures of scale for the spatial and non-spatial components of the covariance, kernel functions such as the exponential, Gaussian, and Matérn include a range parameter; the Matérn kernel also includes a smoothness parameter (Rasmussen and Williams, 2006, Chapter 4).

The problem with inference for **P**^−1^ as defined here is that it is a dense matrix with a dense inverse (precision matrix). We make use of the approximation from Vecchia (1988), wherein only the nearest *k* neighbors are considered when forming the precision matrix **P**, where values of *k* of around 10 to 20 are reasonable. In particular, we make use of the findings of Datta et al. (2016), wherein we may write

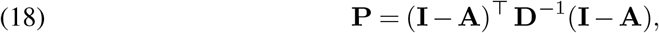

where **A** is a strictly lower-triangular matrix with at most *k* non-zero entries per row and **D** is diagonal—we refer the interested reader to the pseudocode in Finley et al. (2019, Section 2) for the details in forming the matrix **A**. Hence, we have found a sparse approximation for the precision matrix **P** and, similar to the lattice models, enabled rapid inference of the parameter ***β***. We might think of this class of models as related to the lattice models in that only a few neighbors per measurement are considered directly, but with the added flexibility of allowing different kernel functions to model the spatial dependencies.

### S3. Fitting the TESSERA Model

Recall that ***β*** denotes the vector of regression coefficients shared across samples and ***ψ***_*i*_ denotes the parameters for the sample-specific spatial covariance matrices **∑**_*i*_, *i* = 1, …, *n*; for the lattice models, ***ψ***_*i*_ can be partitioned into the parameters 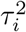 and *γ*_*i*_. Let ***ν*** = (***β, ψ***_1_,…, ***ψ***_*n*_) be the vector of all parameters to be estimated and ***ν***_*i*_ = (***β, ψ***_*i*_) the subset of parameters for the likelihood of sample *i*.

Let 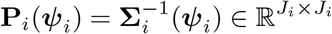 be the precision matrix for sample *i*. For the lattice models, one may write 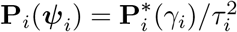, where the different versions of 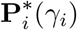 are provided in Table 1.

### S3.1. Approximation of the Distribution of η| Z, X

In both estimating and testing hypothesis for the parameters ***ν***, we rely on a normal approximation to the conditional likelihood of ***η***_*i*_ | **Z**_*i*_, **X**_*i*_, so as to make likelihood computations feasible. In this and the following sections, we draw from the development in Cressie (2015, Section 7.5.2, p. 544– 548) that is based on the work in Clayton and Kaldor (1987).

Specifically, from Clayton and Kaldor (1987), we have that the distribution of ***η***_*i*_ given the observed counts **Z**_*i*_ and covariates **X**_*i*_ (for a single sample) can be approximated with a normal distribution, ***η***_*i*_ | **Z**_*i*_,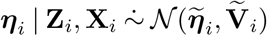, with

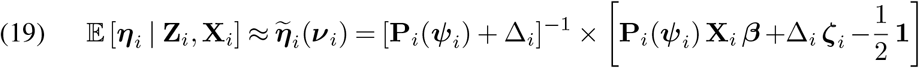

and

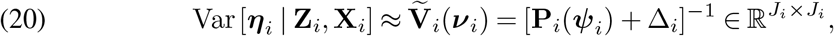

where **1** denotes the vector of all 1, Δ_i_ the diagonal matrix with entries *Z*_*i,j*_ + (1*/*2), and ***ζ***_*i*_ the vector of log((*Z*_*i,j*_ + 1*/*2)*/C*_*i,j*_).

#### S3.2. The Expectation-Conditional-Maximization Algorithm

For the (*k* + 1)st iteration of the expectation-conditional-maximization (ECM) algorithm, we perform the following two steps.

##### S3.2.1. E-Step

Compute the conditional expected value of the joint log-likelihood *L*(***ν***; **Z, *η***) of the observed counts **Z** and the unobserved ***η***, given the observed counts **Z** and covariates **X**

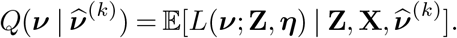

For the lattice models, the joint log-likelihood (up to a constant in ***ν***) is given by

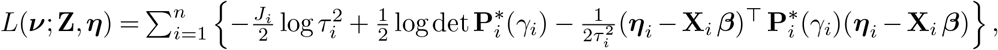

and for the spNNGP model,

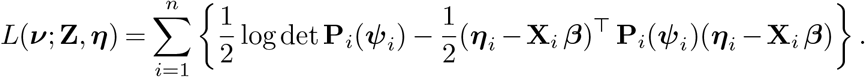

Note that finding 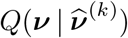 is not equivalent to simply replacing ***η*** with 𝔼 [***η*** | **Z**_*i*_, **X**_*i*_] in the appropriate likelihood equations above, as the expected value of a quadratic form involves also the covariance matrix (cf. Equation (38)), here Var[***η***_*i*_ | **Z**_*i*_, **X**_*i*_]. The calculation of the conditional means and covariance matrices of ***η***_*i*_ | **Z**_*i*_, **X**_*i*_ is intractable. Making use of the normal approximation of the distribution of ***η***_*i*_ | **Z**_*i*_, **X**_*i*_ given in Section S3.1, with 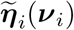 and 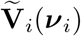 defined in Equations (19) and (20), respectively, we estimate the function *Q*(·) with 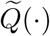 defined as follows. For the lattice models,

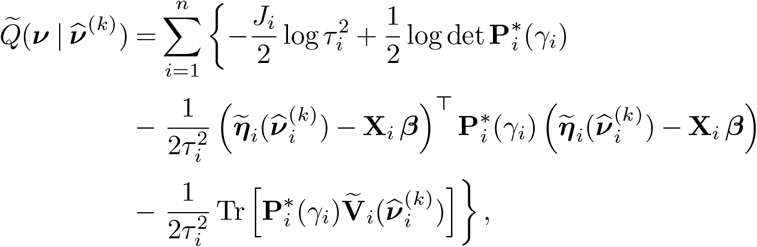

and for the spNNGP

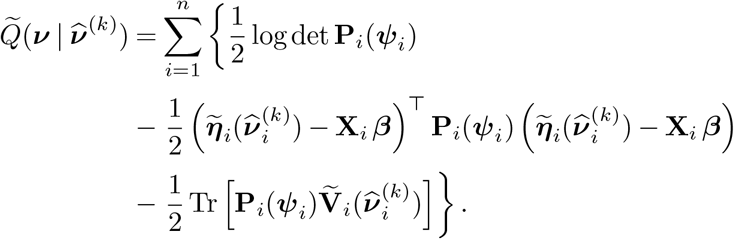

##### S3.2.2. CM-Step

In a traditional M-step, we would compute 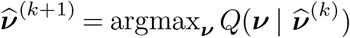 (or 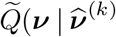 for our E-step approximation). However, as closed form solutions for a joint maximization in all parameters are not available and numerical approximations are computationally intractable, we instead perform a conditional maximization step, where we alternate between maximizing 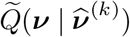 with respect to the sample-specific covariance parameters ***ψ***_*i*_ and the shared regression parameter ***β***.

Note that 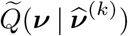 depends on 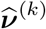 only via the conditional means 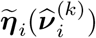 and covariances 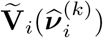. These are constants in the CM-step and are only updated after completion of the CM-step, when 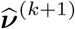 is determined and the next E-Step is performed. Thus, for simplicity in describing the CM-step, we will use the short-hand notation 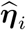 and 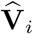 for 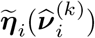 and 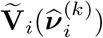, respectively, and write

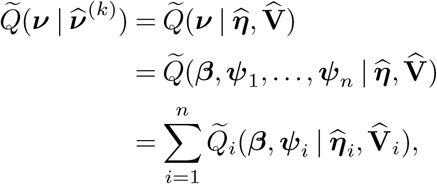

i.e., the sum of the approximate expected conditional log-likelihoods 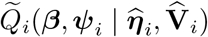 of each sample *i*. Thus, maximizing 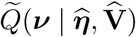 with respect to only the ***ψ***_1_, …, ***ψ***_*n*_ is equivalent to individually maximizing each 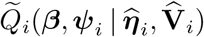 with respect to ***ψ***_*i*_.

The (*k*^*′*^ + 1)st iteration of the CM-step, nested within an iteration *k* of the ECM algorithm, is then as follows.

1. For each sample *i*, maximize 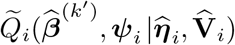 with respect to ***ψ***_*i*_, resulting in an update

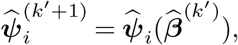

where the notation 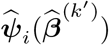 indicates the dependence on the prior CM iteration’s estimate of ***β***. The maximization in ***ψ*** is performed one element at a time, holding the others constant, e.g., in *τ* ^2^ holding *γ* constant, and then in *γ* holding *τ* ^2^ constant, for the lattice models. We derive 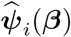 for each covariance model in Section S3.3 below. Note that there are further approximations needed for the maximization of 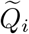 for the spNNGP covariance model, discussed in Section S3.3.6.
2. Maximize 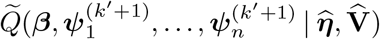with respect to ***β***, resulting in an update

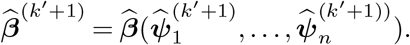

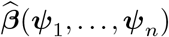 is derived in Section S3.3, Equation (25), and is the same for all covariance models.

These two steps are iterated until the expected log-likelihood 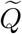 stabilizes, at which point 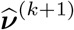 (corresponding to the (*k* + 1)st step of the overall ECM algorithm) is set using the final values,

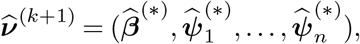

and the algorithm returns to the E-step.

##### S3.2.3. Initialization

For *k* = 0, we initialize ***ν*** with simple estimators. For ***β***, we use a Poisson GLM. For the spatial parameters ***ψ*** in a spNNGP model, we use the `BRISC` method applied to the log-transformed counts. In the lattice models, we initialize *τ* ^2^ with the variance of the log-transformed counts and *γ* with the Moran’s *I* of the counts in each sample. There is no initialization needed for the CM-step; the updates are either closed form (all parameters in the lattice-based covariance models) or rely on `BRISC` for the covariance parameters of the spNNGP covariance model.

##### S3.2.4. Variance Estimation

In addition to the detailed derivations for the CM-step updates, the Hessians for each parameter are given in Section S3.3. The Hessian derivations are used to verify that the estimated parameters actually correspond to local maxima. As the Hessian is equal to the negative of the observed information, it yields an estimate of the variance of the estimators. Due to the lack of identifiability in the spNNGP model (discussed in Section S4), as well as the slightly different procedure for maximizing 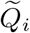 with respect to ***ψ***_*i*_, we do not provide Hessian matrices for the spatial parameters in this model.

##### S3.2.5. Convergence to Global Optima

As noted, the `TESSERA` algorithm relies on an approximation of the expected log-likelihood in the E-Step. While this approximation is generally good (and improves for larger counts), it is still an approximation and a potential source of error (Clayton and Kaldor, 1987). Moreover, the ECM algorithm, like the EM algorithm, can only guarantee convergence to a local optimum of the log-likelihood, as opposed to a global optimum (Meng and Rubin, 1993).

However, we may offer the following weaker guarantee. As detailed above, each of the CM-steps optimizes in a single parameter while holding the others fixed (where we treat ***β*** as a single parameter in this discussion). As long as the precision matrices **P**_*i*_ are invertible, the single-parameter Hessians of the approximated expected log-likelihood 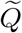 are all negative definite, so that each CM-step update has a unique solution. Hence, the sequence of CM-steps does not decrease 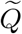.

##### S3.2.6. Prediction

Given estimated parameter values for ***ν***, we can perform prediction of the counts. Using the fact that the estimator that minimizes risk for the squared error loss is the conditional (posterior) mean, we may estimate *θ*_*i,j*_ with

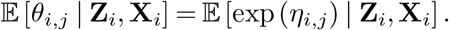

Using the approximate normality of *η*_*i,j*_ **Z**_*i*_, |**X**_*i*_, we have that 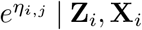 follows a log-normal distribution, giving us

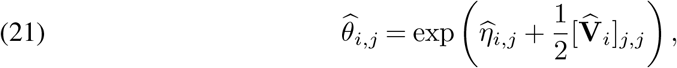

where 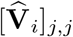 denotes the diagonal elements of the matrix 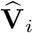 and 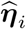 and 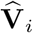 are obtained by plugging in 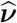 in Equations (19) and (20), respectively. It follows that the fitted values of the counts *Z*_*i,j*_ are given by

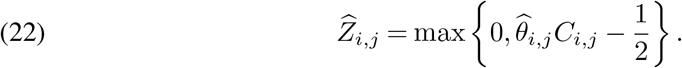

### S3.3. Derivation of TESSERA CM-step Updates

In the CM-step, we seek to maximize

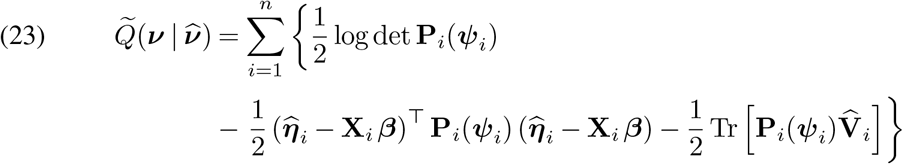

with respect to ***ν*** = (***β, ψ***_1_, …, ***ψ***_*n*_), by alternating maximization steps over the sample-specific covariance parameters ***ψ*** and the common regression parameter ***β***, where 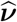 denote the previous iteration’s estimated parameters and 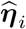 and 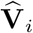 the corresponding estimated ***η***_*i*_ computed according to Equations (19) and (20), conditional means and covariances for respectively.

We first present the CM-step update for ***β***, which applies to all covariance models. For lattice models, 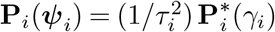 and we present below the CM-step updates for *τ* ^2^ and *γ*. The update for *τ* ^2^ is the same for all lattice models, whereas the update for *γ* is different for each of the CAR, SAR, and Leroux parameterizations of **P**^∗^.

#### S3.3.1. All Covariance Models: β

Assume that we have a precision matrix **P**_*i*_(***ψ***_*i*_) for sample *i*. Assume further that all spatial parameters ***ψ***_*i*_ are held fixed and, for notational simplicity, write **P**_*i*_ instead of **P**_*i*_(***ψ***_*i*_). Differentiating 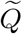 from Equation (23) with respect to ***β*** yields

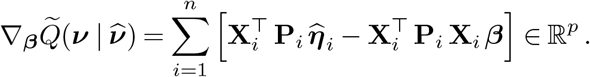

If we define

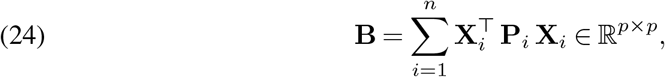

setting the gradient equal to zero yields that, for fixed ***ψ***_*i*_, 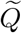 is maximized by

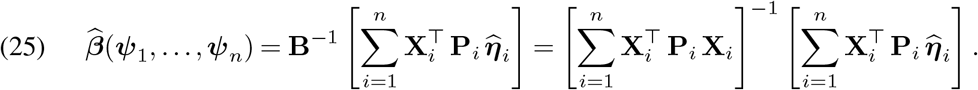

We note that the form of this estimator is that of a generalized least squares estimator. Defining a block-diagonal matrix of the **P**_*i*_ and stacking the rows of **X**_*i*_ accordingly yields the conclusion. Differentiating 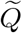 again yields the negative of the observed information equal to the matrix **B**. Note that **B** depends on the spatial covariance parameters ***ψ***_*i*_ via the **P**_*i*_. Assuming that the **P**_*i*_ have full rank (i.e., are positive-definite and proper precision matrices) and that the **X**_*i*_ have full column rank, 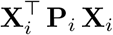 will have full rank and be positive-definite. Since **B** is the sum of positive-definite matrices, it is positive-definite, so that − **B** will be negative-definite and 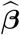 is a maximizer of 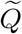.

#### S3.3.2. All Lattice Models: τ ^2^

We drop the sample index *i* for convenience, as differentiating in 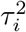 will only involve the terms with index *i*. Assume that ***β*** and *γ* are fixed; then **P**^∗^(*γ*) is also fixed and denoted simply by **P**^∗^. The gradient of 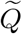 with respect to *τ* ^2^ is

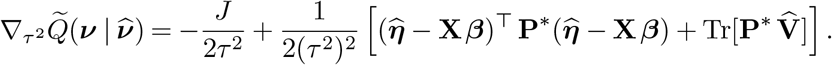

Setting the gradient equal to zero, it follows that the estimator for *τ* ^2^ is

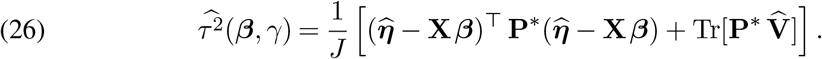

Differentiating again yields the negative of the observed information

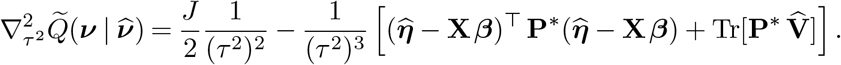

We note that if

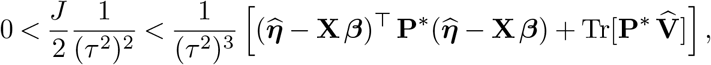

then the Hessian is strictly negative. Simplifying the above, we see that we need

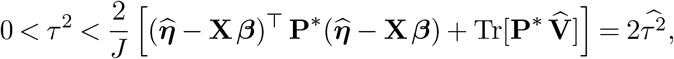

so that the Hessian is always negative at *τ* ^2^.

##### S3.3.3. γ in the CAR Model

In what follows, we omit the sample index *i* for convenience. Assume that ***β*** and *τ* ^2^ are fixed. In the CAR model, we have that **P**^∗^ = (**D** −*γ* **W**), so that the relevant portion of 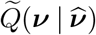 is

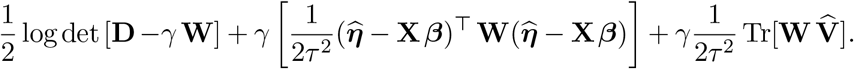

Differentiating the first term with respect to *γ* yields

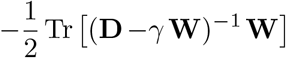

and then differentiating the second and third terms yields

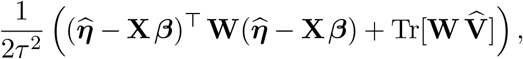

for an overall gradient of

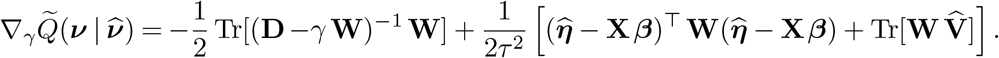

Using the invertibility of **D** and **W**, we may write

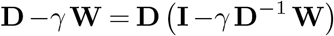

or

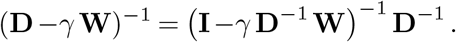

If we let **A** = **D**^−1^ **W**, we may write the first term of the gradient as

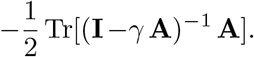

Assuming that **A** has an eigendecomposition **A** = **U Λ U**^−1^, where **Λ** has diagonal entries *λ*_*j*_, *j* = 1, …, *J*, we may write

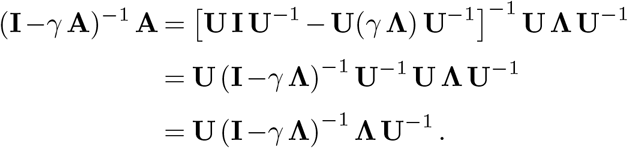

Taking the trace yields

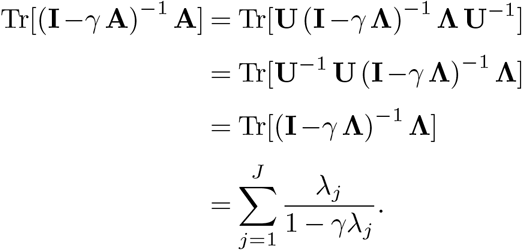

It follows that the gradient in *γ* is

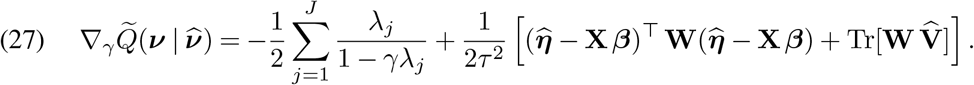

We may set the gradient equal to zero and numerically look for a root in [−1, 1] to find the 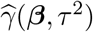 which maximizes 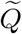 for fixed ***β*** and *τ* ^2^. Pre-computing **A** and its eigenvalues for each sample leads to an extremely rapid evaluation of the gradient, making the optimization more efficient.

Differentiating again and negating yields the observed information:

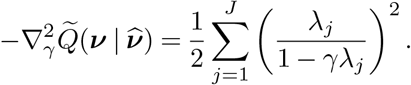

Note that the Hessian is strictly negative, as at least one of the *λ*_*j*_ is non-zero.

##### S3.3.4. γ in the SAR Model

In what follows, we omit the sample index *i* for convenience. Assume that ***β*** and *τ* ^2^ are fixed. Here, the relevant part of 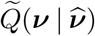 is

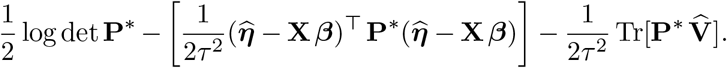

In the SAR model, we have that

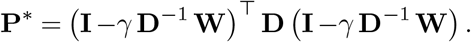

Reusing the definition of **A** = **D W**, we may expand **P**^∗^ as

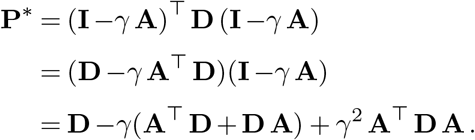

Note that since **A** = **D**^−1^ **W** and (**D A**)^⊤^ = **A**^⊤^ **D**, we may write **D A** = **D D**^−1^ **W** = **W**, so that **A**^⊤^ **D** + **D A** = 2 **W**. Similarly,

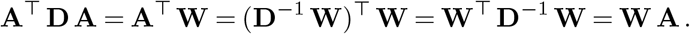

If we define

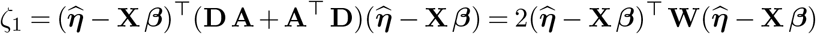

and

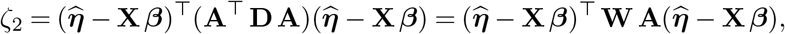

we may write the relevant part of the second term of 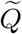 as

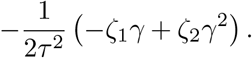

Differentiating the above with respect to *γ* yields

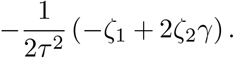

By a similar series of derivations, we may write the relevant part of the third term of 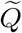 as

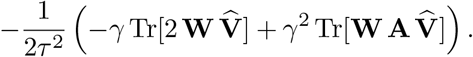

Differentiating in *γ* yields

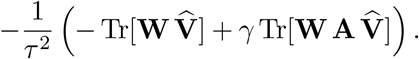

For the first term of 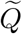, note that

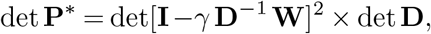

so that

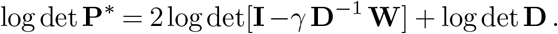

Differentiating this in *γ*, using Equation (39), yields

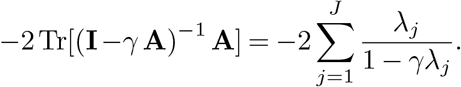

It follows that the overall gradient is (28)

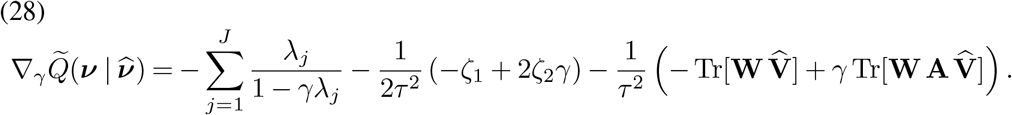

We may set the gradient equal to zero and numerically look for a root in [−1, 1] to find the 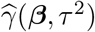 which maximizes 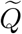 for fixed ***β*** and *τ* ^2^. Note that the overall form of the gradient is very similar to that in the CAR model.

If we differentiate again in *γ* and negate the result, we find an observed information of

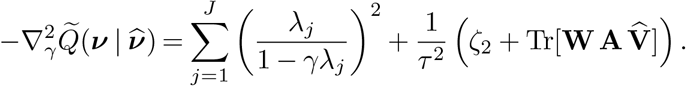

We note that the first term of the Hessian is strictly negative, *ζ*_2_ ≥ 0, and 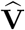 and **W** are positive-semidefinite; however, we cannot conclude that the Hessian is strictly negative unless the product **W A** 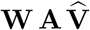 has a postive trace.

##### S3.3.5. γ in the Leroux Model

In what follows, we omit the sample index *i* for convenience. Assume that ***β*** and *τ* ^2^ are fixed. In the Leroux model, we have that

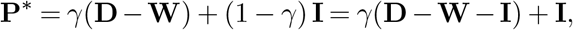

so that the relevant part of 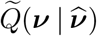 is

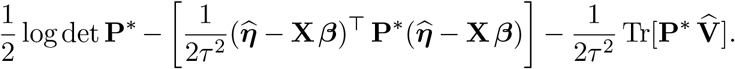

Considering the first (log-determinant) term, we may write

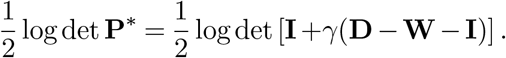

Differentiating in *γ* yields a gradient (of this term) of

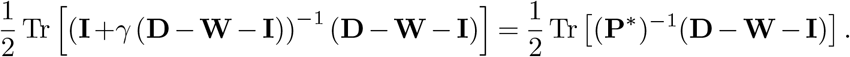

If we write the eigendecomposition of **D W** = **U Λ U**^−1^, where **Λ** has diagonal entries *λ*_*j*_, for *j* = 1, …, *J*, we have that

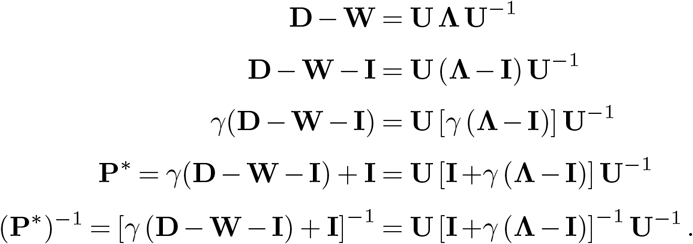

Then, we have that

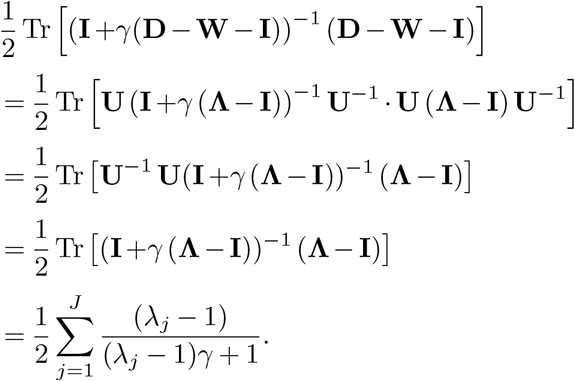

The second term of 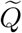 can be written as

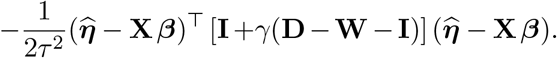

Isolating the part that contains *γ* yields

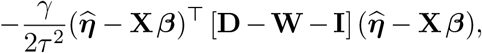

so that the gradient of this term is simply

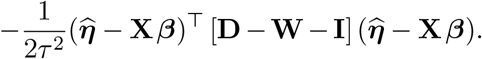

The relevant part of the final trace term is

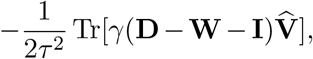

where differentiating yields

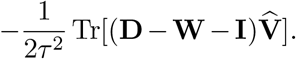

The overall gradient is then

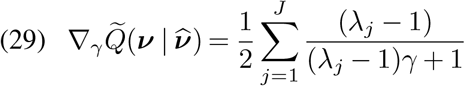

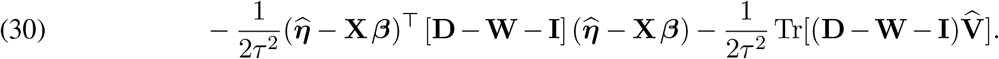

Again, we may set the gradient equal to zero and numerically look for a root in [0, 1] to find the 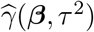 which maximizes 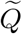 for fixed ***β*** and *τ* ^2^. Much like in the CAR and SAR cases, precomputing the eigenvalues of **D ™W** dramatically improves the ability to quickly fit these models.

Differentiating again yields a Hessian (negative of the observed information) of

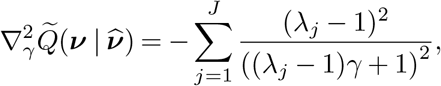

which is clearly negative.

##### S3.3.6. Spatial Correlation Parameters ψ in the spNNGP Model

Estimation of the parameters ***ψ***_*i*_ depends on the kernel function chosen, and performing a proper maximum likelihood optimization procedure has cubic computational complexity as a function of the number of cells in each sample (Saha and Datta, 2018). Instead, we use the `BRISC` algorithm applied to 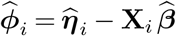. This procedure has quadratic time complexity and is much more scalable, in addition to being robust and accurate. Given the parameter estimates, we follow Datta et al. (2016) in forming the precision matrix **P**_*i*_ and the estimation of ***β*** proceeds as in the lattice models.

### S4. Identifiability of Parameters

In this section, we show that under relatively mild conditions, the fixed effects ***β*** are identifiable in all models we consider. Moreover, we show that the spatial parameters *γ* and *τ* ^2^ are identifiable in the lattice models. Due to the independence of the ***η*** across samples, it is sufficient to consider a single sample; we hence drop the sample index *i* where possible (though we include a verification that this single-sample focus is valid in Corollary 2). Moreover, as the counts **Z** only depend on the parameters ***ν*** via ***η***, we need not consider the joint distribution of **Z** and ***η***. That is, it is sufficient to consider the likelihood of ***η*** ∈ ℝ^*J*^ conditional on the covariates **X**. To prove identifiability, we show that if the difference in the log-likelihoods of ***η*** with two different parameter sets is zero, both sets of parameters must be the same. We first prove that the fixed effects ***β*** and the precision matrices **P** = **∑**^−1^ are identifiable in Theorem 1. We then prove in Theorem 3 that the spatial parameters *γ* and *τ* ^2^ are identifiable for each of the CAR, SAR, and Leroux models. We discuss the (lack of) identifiablilty of the spatial parameters in a spNNGP model at the end of this section.

#### Theorem 1.

*Consider the single-sample version of the* `TESSERA` *model of Equation* (3), *where* ***η*** | **X** *∼* 𝒩 (**X *β*, ∑**). *If we assume that*

1. *the covariate matrix* **X ∈** ℝ^*J×p*^ *has full column rank (so that p ≤ J)*,
2. *the spatial parameters are such that the precision matrix* **P** = **∑**^−1^ ∈ ℝ^*J×J*^ *of* ***η*** *is invertible*,
3. *the number of measurements J is at least* 2, *then we have that the fixed effects* ***β*** *∈* ℝ^*p*^ *and the precision matrix* **P** *∈* ℝ^*J×J*^ *are identifiable*.

Before proceeding, we return to the multi-sample setting.

#### Corollary 2.

*Consider the multi-sample* `TESSERA` *model of Equation* (3), *where* ***η***_*i*_ | **X**_*i*_ *∼* 𝔼 (**X**_*i*_ ***β*, ∑**_*i*_) *and the* ***η***_*i*_ *are independent across samples. If each precision matrix* **P**_*i*_ *is invertible, there are at least two measurements across all samples (*∑ _*i*_ *J*_*i*_ ≥ 2*), and the stacked covariate matrix* 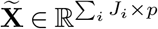 *of the* **X**_*i*_ *has full column rank, then* ***β*** *and the* **P**_*i*_ *are all simultaneously identifiable*.

Proof. If we let 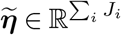 be the stacked vector of the ***η*** _*i*_ and 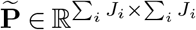 be the block-diagonal matrix of the **P**_*i*_, then 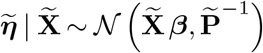 . We may then apply Theorem 1. That is, **P** is invertible if all of the **P**_*i*_ are, and, by assumption, 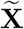 has full column rank. We may repeat our analysis with 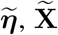, and 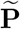 replacing ***η*, X**, and **P**. We conclude that ***β*** and 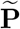 are identifiable. As 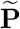 is block-diagonal with blocks **P**_*i*_, it follows that the **P**_*i*_ are identifiable.

We now consider the lattice models. In Theorem 1, we have shown that there must be a unique precision matrix, so it remains to show that only one set of parameters could generate a given **P**.

#### Theorem 3.

*In addition to the assumptions in Theorem 1, assume that:*

1. *(CAR, SAR) The fixed and known adjacency matrix* **W** *(and hence the degree matrix* **D***) has no rows or columns that are identically zero*.
2. *(Leroux) The fixed and known adjacency matrix* **W** *has at least one non-zero entry*.
3. *The value of τ* ^2^ *is strictly greater than zero*.
4. *The value of γ is such that the precision matrix* **P** *is invertible and positive-definite*.

*Then, we have that the parameters γ and τ* ^2^ *in the CAR, SAR, and Leroux models are identifiable*.

The first condition is necessary for the precision matrix **P** to be invertible in the CAR and SAR models. The second condition is a somewhat trivial technical condition that allows the Leroux model to have spatial correlation (otherwise the covariance would be diagonal and totally non-spatial). The third and fourth conditions apply to all three lattice models and ensure that the distribution of ***η*** is proper (i.e., integrates to 1).

#### S4.1. Conclusions for the Lattice Models

From Theorems 1 and 3 and Corollary 2, we conclude that the parameters 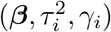 are identifiable in the lattice models under the set of mild assumptions specified above. Numerically, we see in our simulations that all spatial covariance parameters indeed appear to be identifiable and that ***β*** is identifiable in all models.

#### S4.2. The spNNGP Model

For the spNNGP model, we draw upon the Gaussian process literature and note that the kernel parameters ***ψ***_*i*_ are generally not identifiable (Arendt, Apley and Chen, 2012; Brynjarsdóttir and O’Hagan, 2014), though recent work has discovered some conditions under which identification is possible (Plumlee and Joseph, 2018; Kim and Lee, 2021; Tang, Zhang and Banerjee, 2021; Chen et al., 2023). However, Theorem 1 still applies, and thus ***β*** is still identifiable in this setting.

### S5. Identifiability Proofs

#### S5.1. Proof of Theorem 1

Proof. We consider two sets of parameter values for the regression coefficient ***β*** and the precision matrix **P**, namely ***β***_1_ and ***β***_2_ ∈ ℝ^*J*^ and **P**_1_ and **P**_2_ ∈ ℝ^*J×J*^ . The log-likelihood of ***η*** given **X** is given by

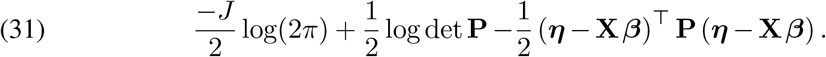

If we take the difference between the log-likelihoods at the two sets of parameter values and set it to zero, we find (after dropping a common factor of 1*/*2):

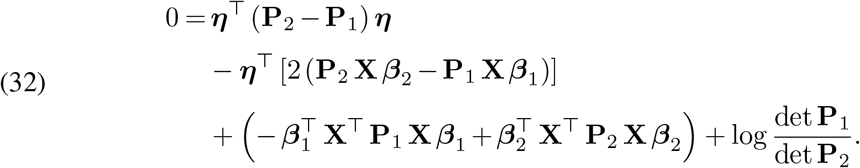

Importantly, Equation (32) must hold for all ***η*** and has the same form as Equation (42) in Lemma 5. We may apply the Lemma and conclude that:

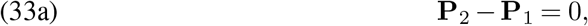

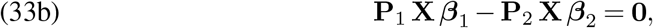

and,

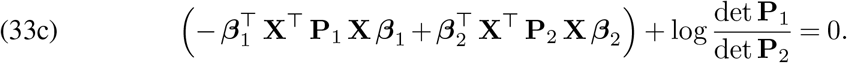

Via Equations (33a) and (33b), we conclude that **P**_1_ = **P**_2_, so that

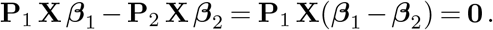

By assumption, **P**_1_ and **X** have full column rank, so that the nullspace of **P**_1_ **X** is only the zero vector. It follows that ***β***_1_ − ***β***_2_ = **0**.

#### S5.2. Proof of Theorem 3

We separate the proof into three parts, corresponding to each of the three lattice models. For each part, we consider two sets of parameters: *γ*_1_ and *γ*_2_, and 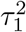 and 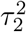. We consider the explicit forms of the precision matrices **P**_1_ = **P**_2_ and argue that the parameters must be identical.

##### S5.2.1. CAR

Proof. For a CAR model, we may write

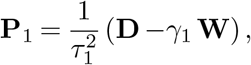

and similarly for **P**_2_. As **W** has no zero rows or columns, there exists a non-zero off-diagonal entry of **W**, denoted *W*_*j,j*_*′* . The corresponding entry of **P**_1_ has the form 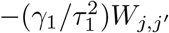 and that for **P**_2_ has the form 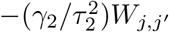 For these two values to be equal, we must have 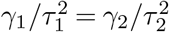. Now, we consider and equate a given diagonal entry of both precision matrices:

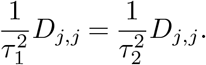

Since by assumption *D*_*j,j*_ is strictly positive, the only way for this statement to hold is if 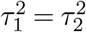. Finally, if 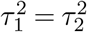 and 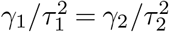, we can conclude that *γ*_1_ = *γ*_2_.

##### S5.2.2. Leroux

Proof. For a Leroux model, we may write

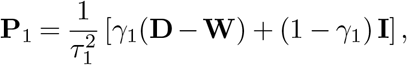

and similarly for **P**_2_. By assumption, there is at least one non-zero off-diagonal element of **W**, denoted *W*_*j,j*_*′* . The corresponding entry of **P**_1_ has the form 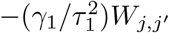 and that for **P**_2_ has the form 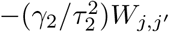 For these two values to be equal, we must have 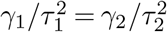. Now, considering and equating the diagonal entries of **P**_1_ and **P**_2_ yields

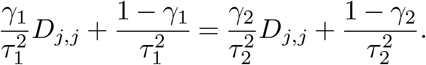

Simplifying and using the fact that 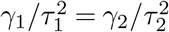 yields that 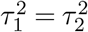. Finally, if 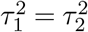 and 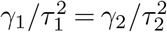, we can conclude that *γ*_1_ = *γ*_2_.

##### S5.2.3. SAR

Proof. For a SAR model, we may write

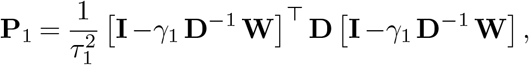

and similarly for **P**_2_. Expanding this definition and simplifying, we may write

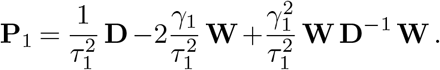

For **P**_1_ − **P**_2_ = 0 to hold, we require

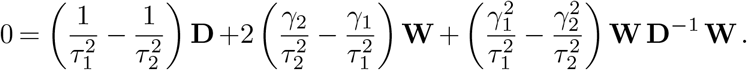

Equivalently, defining **A** = **D**^−1^ **W** and multiplying from the left with **D**^−1^ yields

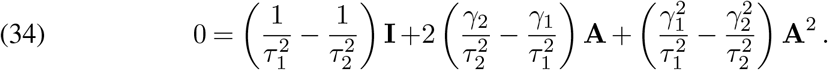

Equation (34) says that a polynomial of degree two annihilates **A**. Immediately, we see that if **A** has more than two distinct eigenvalues, all coefficients of this polynomial must be zero, as then a polynomial of degree two cannot annihilate **A** since all of the eigenvalues of **A** must be roots of any annihilating polynomial of **A** (see Hoffmann and Kunze (1971, Sec. 6.3) for the mathematical background). We would then conclude that every coefficient of Equation (34) is zero, so that 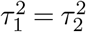 and *γ*_1_ = *γ*_2_.

If **A** has two or fewer distinct eigenvalues, the analysis is more complicated. Since **A** is a stochastic matrix (the rows sum to 1 and the entries are non-negative), we know the eigenvalues to be less than or equal to 1 in magnitude by the Gershgorin Circle Theorem—see Landau and Odlyzko (1981) for a discussion of the eigenvalues of matrices like **A**. Moreover, since **A** is stochastic, the leading eigenvalue is 1—consider that the vector of all 1s is an eigenvector with eigenvalue 1. We consider three cases for the eigenvalues of **A**: two distinct eigenvalues that have different magnitudes (1 and |*λ*| ≠ 1), two distinct eigenvalues that have the same magnitude (necessarily ±1), and a single distinct eigenvalue (necessarily *±* 1). In all three cases, we will assume that Equation (34) is not identically zero and derive a contradiction.

For the first case, assume that **A** has two distinct eigenvalues that have different magnitudes. To achieve this, we require both 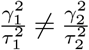 and 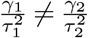 so that the linear and quadratic terms in Equation (34) are non-zero. There are then two possibilities for the eigenvalues:

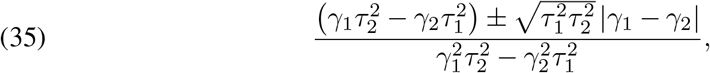

where we have applied the quadratic formula to Equation (34). The largest of the two eigen-values, corresponding to the addition, must be 1. Setting Equation (35) to 1 and solving yields the solution

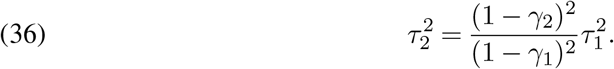

This solution holds for 0 *< γ*_1_ *< γ*_2_ *<* 1 and for −1 *< γ*_1_ ≤ 0 with 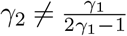. As long as *γ* ≠ *γ* or 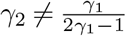 for *γ*≠ 1*/*2, the constraint that 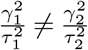 holds. We see that with this choice of 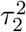, the other root has value

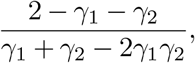

where we have substituted Equation (36) into (35) and taken the negative root. For some eigenvalue *λ*≠ *±*1 of **A**, we can solve for *γ*_2_ in terms of *γ*_1_ and *λ*, where we find

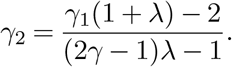

However, *γ*_2_ must be less than 1, which only holds for −1 *< γ*_1_ *<* 0 and 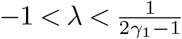. But *γ*_2_ must also be greater than −1, which only holds either for −1 *< γ*_1_ *<* 0 and 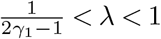 or for 0 *< γ*_1_ *<* 1 and |*λ*| < 1. That is, we cannot simultaneously bound *γ*_2_ from above and below. We conclude that there is no pair of distinct parameters that lead to a valid solution, or, that the SAR model is identifiable in this setting.

For the second case, assume that **A** has two distinct eigenvalues with the same magnitude. Since **A** is a stochastic matrix, these values must be *±*1. Looking at the quadratic formula, we require 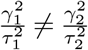 but that 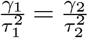 to have roots of identical magnitude and opposite sign. We would then conclude that the non-zero eigenvalues of **A** must be

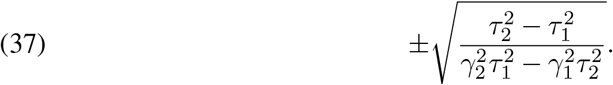

We find that setting

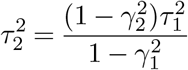

leads to an eigenvalue of 1. To satisfy the constraint 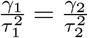, we find that we would need *γ*_1_ = *γ*_2_, but then 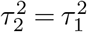 —a contradiction to our assumption. Hence, if **A** has eigenvalues *±*1, the SAR model is identifiable.

For the third case, assume that **A** has only a single distinct eigenvalue. As **A** is a stochastic matrix, this eigenvalue is 1. If we assume that the quadratic term in the polynomial in **A** is identically zero, or, that 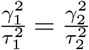 but that 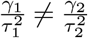, we would find that a linear polynomial annihilates **A**. That is, **A** would be diagonalizable (as the minimal polynomial has no repeated roots) and have eigenvalues equal to 1. However, the only such matrix is the identity matrix: **W** and hence **A** have zeros along the diagonal, so we cannot have a linear polynomial annihilate **A**. So, we then must assume that Equation (34) is proportional to (**A I**)^2^. If this condition holds, the quadratic and constant coefficients must be equal and the quadratic coefficient must be −1*/*2 times the linear coefficient:

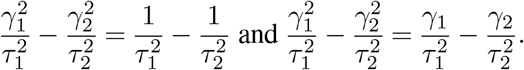

For both equations to hold, we would need *γ*_1_ = *γ*_2_ = 1, but then Equation (34) would only have a non-zero constant term—a contradiction that would lead to 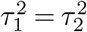. If we took *γ*_1_ = *γ*_2_≠ 1, we would find that we needed 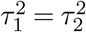 —a contradiction once again as Equation (34) is now identically zero. We conclude that this third case is also not possible and that the SAR model is identifiable in this setting.

### S6. Useful Results

#### S6.1. Useful Identities

1. For a random variable **X** with mean 𝔼 [**X**] = ***µ*** and covariance matrix Var[**X**] = **∑** and a constant matrix **A**,

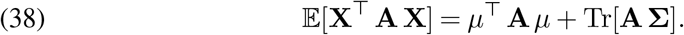
2. For matrices **X** and **Y** and a scalar *t*, we have (Dattorro, 2010, D.2.4)

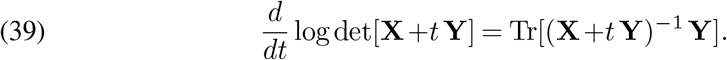
3. For a matrix **A** *∈* ℝ^*p×p*^, vectors **b** and **x** *∈* ℝ^*p*^, and a scalar *c*,

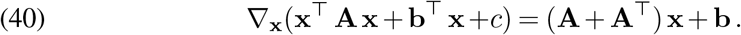

#### S6.2. Useful Definitions

##### Definition 4 (Moran’s *I*).

For a vector **z ∈** ℝ^*J*^ with entries *z*_*j*_, let 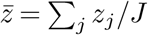. The Moran’s *I* is defined as

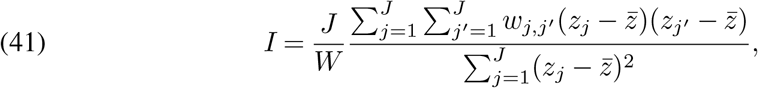

where the *w*_*j,j*_*′* are spatial weights, with *w*_*j,j*_ = 0 and *W* = ∑_*j,j*_*′ w*_*j,j*_*′* (Cressie, 2015, Sec. 6.5.1).

For a lattice model, the entries of the adjacency matrix **W** can be used in (41), or, if coordinates are available, the inverse of the Euclidean distance between pairs of points is another common choice.

#### S6.3. Useful Lemmas

##### Lemma 5.

*If for a square, symmetric matrix* **A ∈** ℝ^*J×J*^, *a vector* **b ∈** ℝ^*J*^, *and a scalar c we have that*

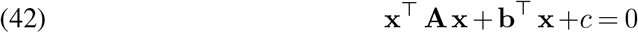

*for all vectors* **x** *∈* ℝ^*J*^, *then we must have that all of* **A, b**, *and c are zero*.

Proof. If Equation (42) holds for all **x**, then the gradient must also be identically zero for all **x**. Taking the gradient in **x**, we find that

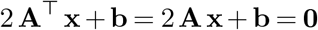

or that

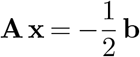

for all **x**. Assuming **A** is not identically zero, then there must exist a non-zero **x** that is not in the nullspace of **A**. Then, for any scalar *α*, we have that

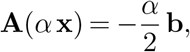

but we also know that, by assumption,

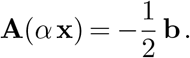

Thus, we must have

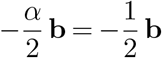

for all *α ∈* ℝ, so that **b** must be identically zero. But if **b** is identically zero, we have that **Ax** = 0 for all **x**. If this holds, we must have that the nullspace of **A** is all of ℝ^*J*^, so that **A** is entirely zero, which is a contradiction of our assumption that **A** is not identically zero. Then, taking **A** to be the zero matrix, we must have that

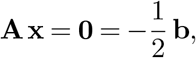

or that **b** must be identically zero. Returning to Equation (42), if **A** and **b** are zero, we require *c* = 0.

### S7. Supplementary Figures

#### S7.1. Synthetic Data: Method Runtime

**Figure S-1:**
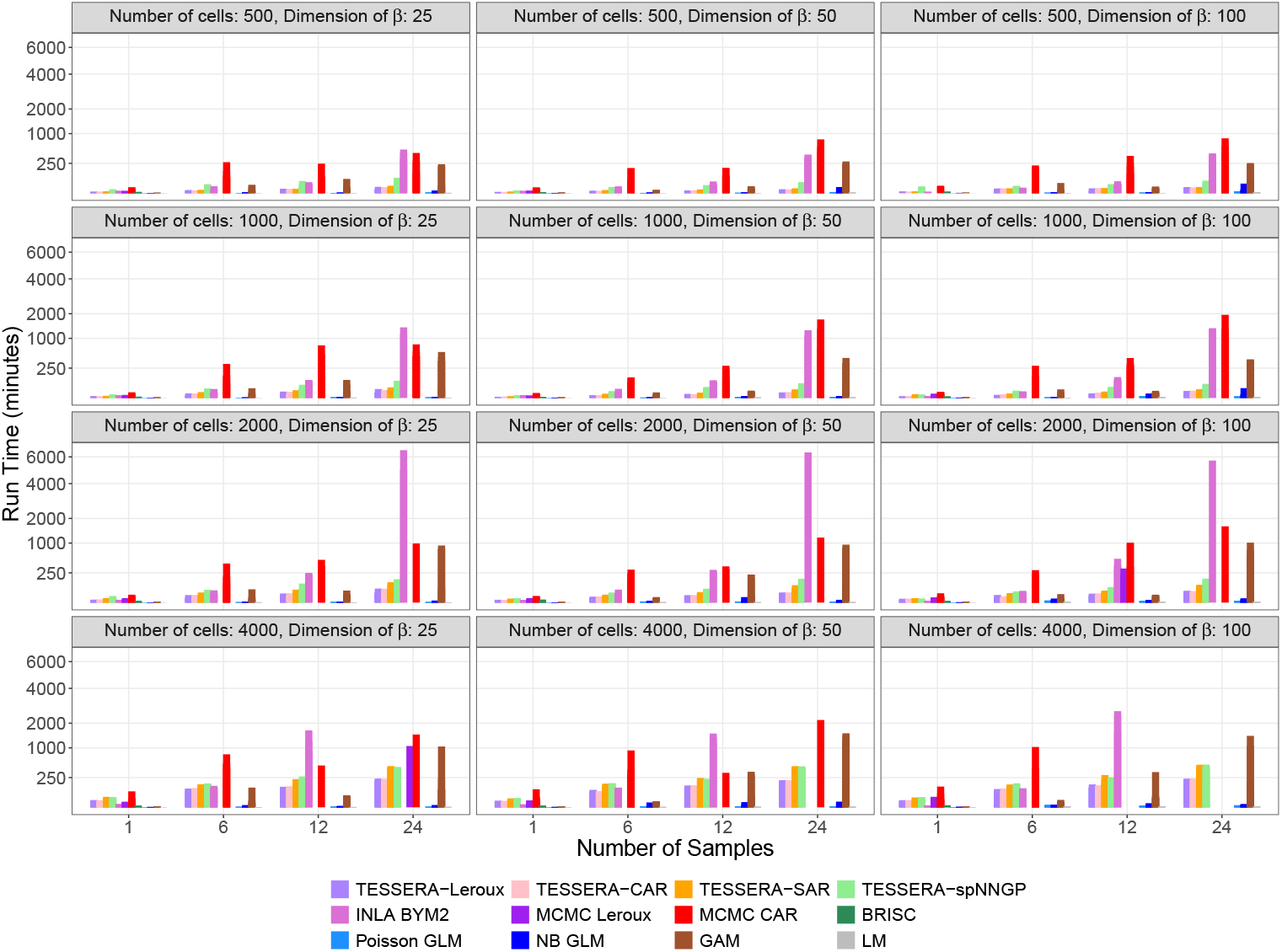
Computational scalability of TESSERA across different simulation scenarios. We evaluated method run time by simulating Poisson-Leroux-distributed data for two conditions with balanced sample sizes and varying numbers of cell types within condition. This simulation was designed exclusively to assess computational scalability of model fitting; it was not used for hypothesis testing or evaluating statistical power. We systematically varied the number of samples (*n* ∈ {1, 6, 12, 24}, where *n* = 1 represents a single-sample baseline and *n >* 1 represents two balanced groups of *n/*2 samples each), the number of cells per sample (*J*_*i*_ ranging from 500 to 4,000), and the fixed-effect dimensionality (dim(***β***) ≈ {25, 50, 100}). For *n* = 1, ***β*** consists solely of cell type effects; otherwise, it incorporates disease, sample, and cell type effects. The number of cell types is then chosen so as to achieve a given dim(***β***). Spatial parameters were derived from empirical fits to the `IGHG2` gene across 14 individual samples from a real-world kidney dataset (Abedini et al., 2024). Specifically, we obtained 14 pairs of spatial covariance parameters (*γ, τ* ^2^) from these fits. For each simulation, we selected covariance parameters by sampling at random, with replacement from this pool of 14 empirical pairs to achieve the desired sample size. Fixed effects ***β*** were drawn from a normal distribution matching the empirical mean and variance of the entries of the estimated ***β*** corresponding to each condition. Cell coordinates were sampled uniformly in the unit square and distance-thresholded to maintain an average of five neighbors per cell. Each simulation was run on a single thread of a 2.4 GHz Intel Xeon processor with 22 GB of RAM. Reported run times represent the average across five independent simulation trials. `TESSERA` was implemented with a 30-iteration limit, a threshold typically sufficient for convergence on the real-world data; implementation details for the compared methods are provided in Table 2. Facets represent unique combinations of cell count and fixed-effect dimensionality. Missing bars indicate simulation scenarios where a method (e.g., INLA or MCMC samplers) failed to terminate due to convergence issues, memory constraints, or time-limit exhaustion.

#### S7.2. Synthetic Data: Comparison of TESSERA with Multi-Sample Methods

**Figure S-2:**
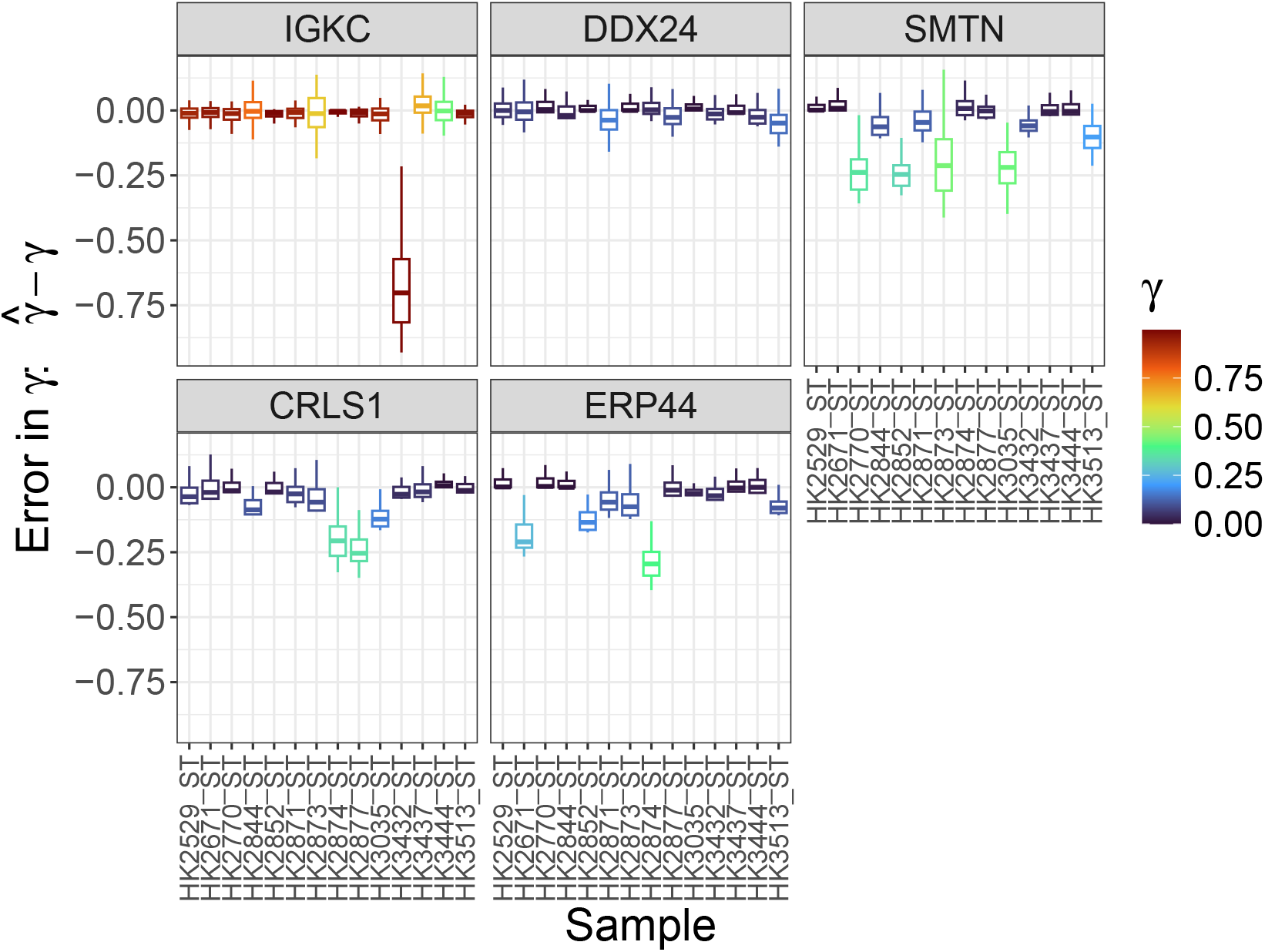
Error in estimating the sample-specific spatial covariance parameter *γ*. To evaluate how effectively `TESSERA` estimates the sample-specific spatial covariance parameter *γ ∈* [0, 1), we examine the estimation error 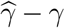 across 100 independent simulation trials. Synthetic data were generated from a Poisson-Leroux distribution with generative parameters estimated from the multi-sample kidney dataset (Abedini et al., 2024), for a representative subset of five genes selected based on their raw count variances across all cells and samples: *IGKC* (maximum variance), *DDX24* (third quartile), *SMTN* (median), *CRLS1* (first quartile), and *ERP44* (minimum). Each facet corresponds to a given gene, and facets are ordered row-wise and top-to-bottom by decreasing expression variance. Within each facet, each boxplot summarizes the estimation error over 100 trials for each of the 14 kidney samples and is color-coded by the true generative *γ* value used in the simulation (see legend). `TESSERA` fit with a Poisson-Leroux model yields consistently low estimation error for *γ* across a selection of genes exhibiting a wide range of variances in expression measures (with the exception of one sample for the highly variable gene *IGKC*).

**Figure S-3:**
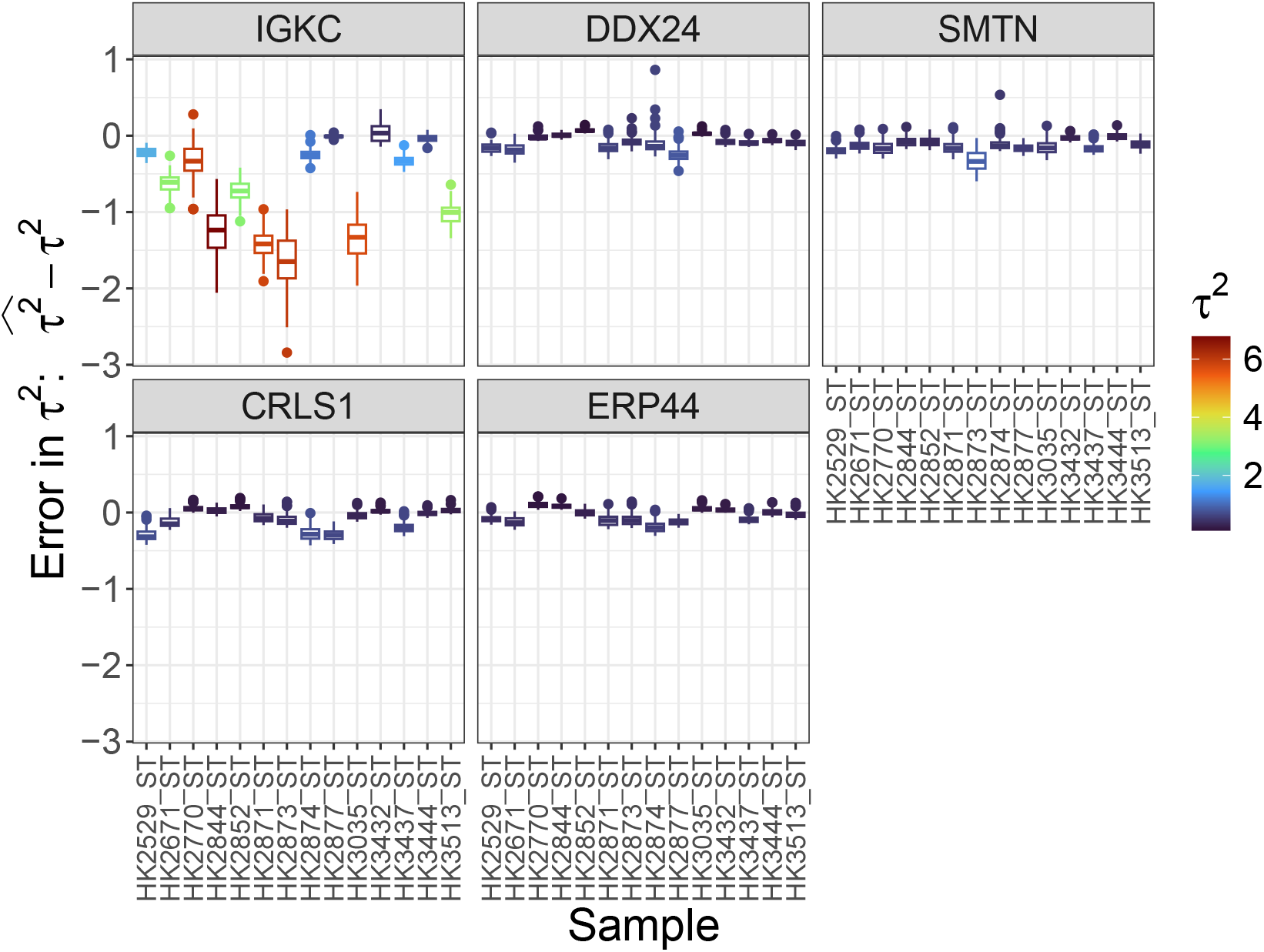
Error in estimating the sample-specific spatial covariance parameter *τ*^2^. To evaluate how effectively `TESSERA` estimates the sample-specific spatial covariance parameter *τ* ^2^ ∈ (0, *∞*), we examine the estimation error 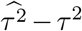 across 100 independent simulation trials. Synthetic data were generated from a Poisson-Leroux distribution with generative parameters estimated from the multi-sample kidney dataset (Abedini et al., 2024), for a representative subset of five genes selected based on their raw count variances across all cells and samples: *IGKC* (maximum variance), *DDX24* (third quartile), *SMTN* (median), *CRLS1* (first quartile), and *ERP44* (minimum). Each facet corresponds to a given gene, and facets are ordered row-wise and top-to-bottom by decreasing expression variance. Within each facet, each boxplot summarizes the estimation error over 100 trials for each of the 14 kidney samples and is color-coded by the true generative *τ* ^2^ value used in the simulation (see legend). `TESSERA` fit with a Poisson-Leroux model yields consistently low estimation error for *τ* ^2^ across a selection of genes exhibiting a wide range of variances in expression measures. Note that although the absolute estimation error is higher for *IGKC* than for the other genes, this gene also has higher true parameter value for *τ* ^2^. Thus, the absolute error remains small relative to the magnitude of the parameter.

**Figure S-4:**
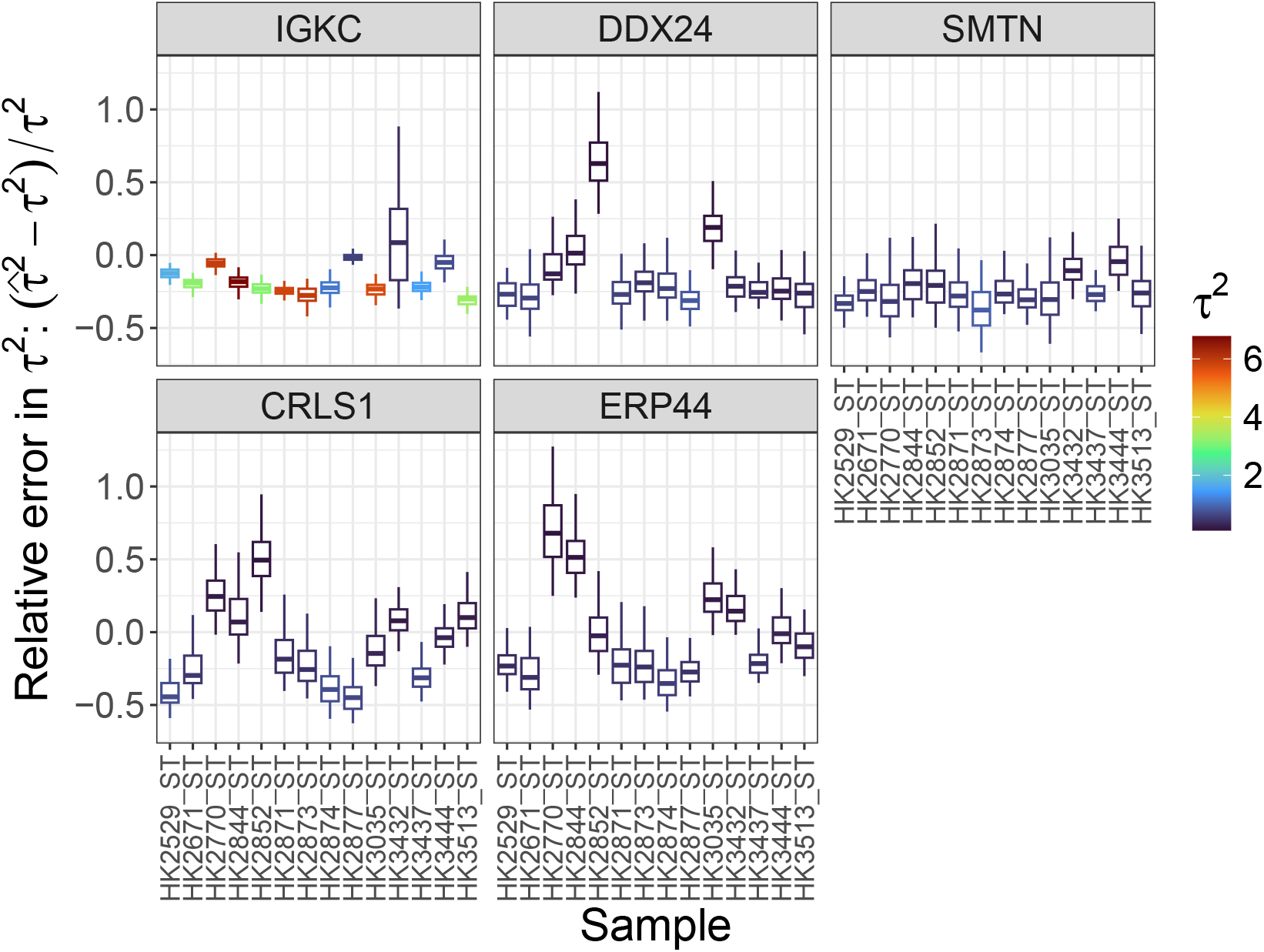
Error in estimating the sample-specific spatial covariance parameter *τ*^2^. To evaluate how effectively `TESSERA` estimates the sample-specific spatial covariance parameter *τ* ^2^ ∈ (0, *∞*), we examine the relative estimation error 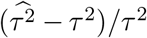 across 100 independent simulation trials. Synthetic data were generated from a Poisson-Leroux distribution with generative parameters estimated from the multi-sample kidney dataset (Abedini et al., 2024), for a representative subset of five genes selected based on their raw count variances across all cells and samples: *IGKC* (maximum variance), *DDX24* (third quartile), *SMTN* (median), *CRLS1* (first quartile), and *ERP44* (minimum). Each facet corresponds to a given gene, and facets are ordered row-wise and top-to-bottom by decreasing expression variance. Within each facet, each boxplot summarizes the estimation error over 100 trials for each of the 14 kidney samples and is color-coded by the true generative *τ* ^2^ value used in the simulation (see legend).

**Figure S-5:**
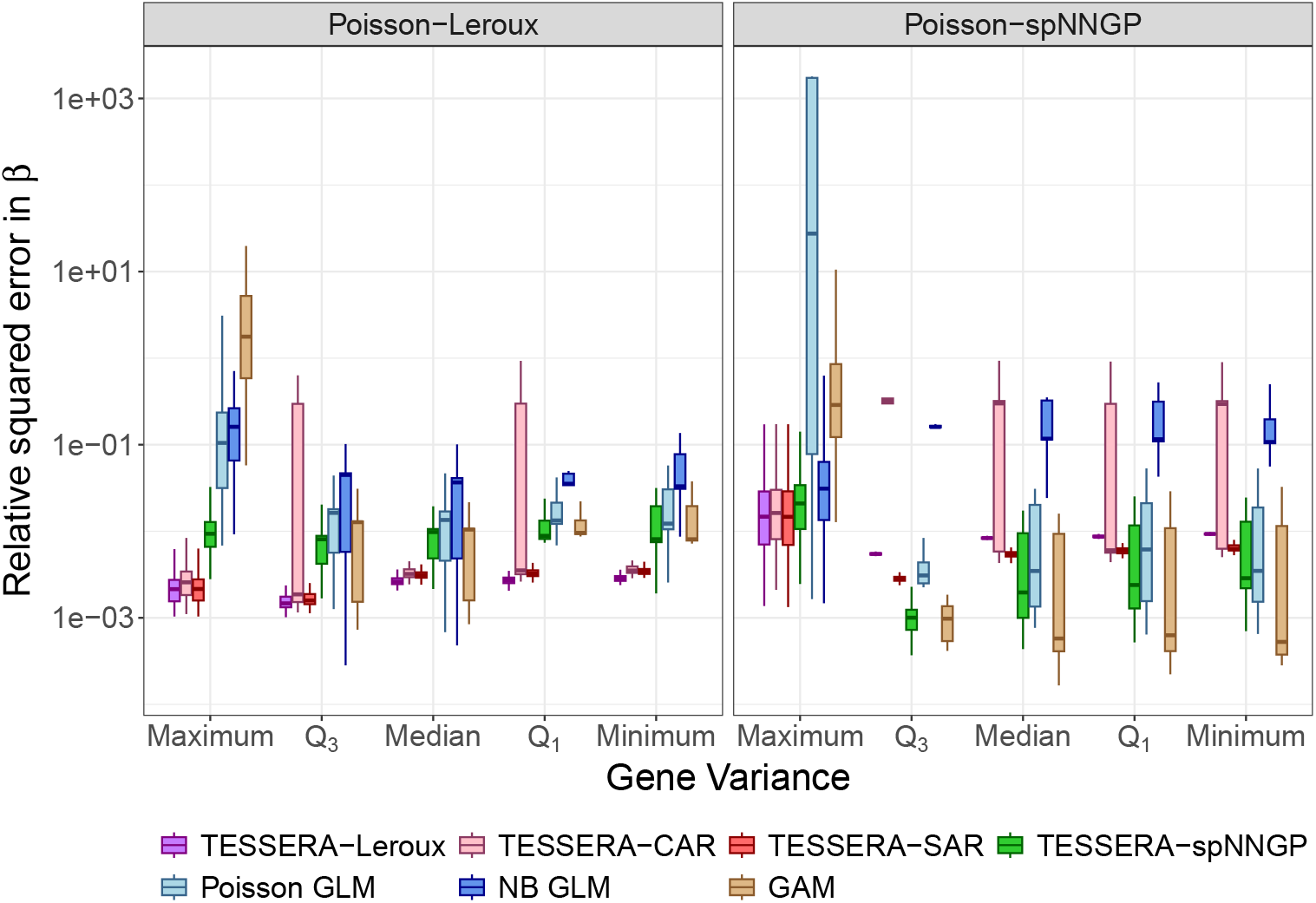
Relative squared error in estimating the shared fixed effect *β*. Synthetic data were generated using Poisson-Leroux and Poisson-spNNGP distributions, with generative parameters estimated from a multi-sample real-world kidney dataset (Abedini et al., 2024). We compared `TESSERA` fit with lattice and spNNGP models against a Poisson GLM, an NB GLM, and a GAM. Estimation accuracy of the fixed effects ***β*** was assessed via the relative squared error, 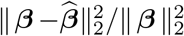, calculated across 100 independent simulation trials. Performance is displayed for a representative subset of genes covering the full distribution of observed expression variance; specifically, genes were selected corresponding to the maximum (*IGKC*), third quartile (*DDX24*), median (*SMTN*), first quartile (*CRLS1*), and minimum (*ERP44*) values of raw count variance across all cells and samples. Facets represent the underlying data-generating models.

**Figure S-6:**
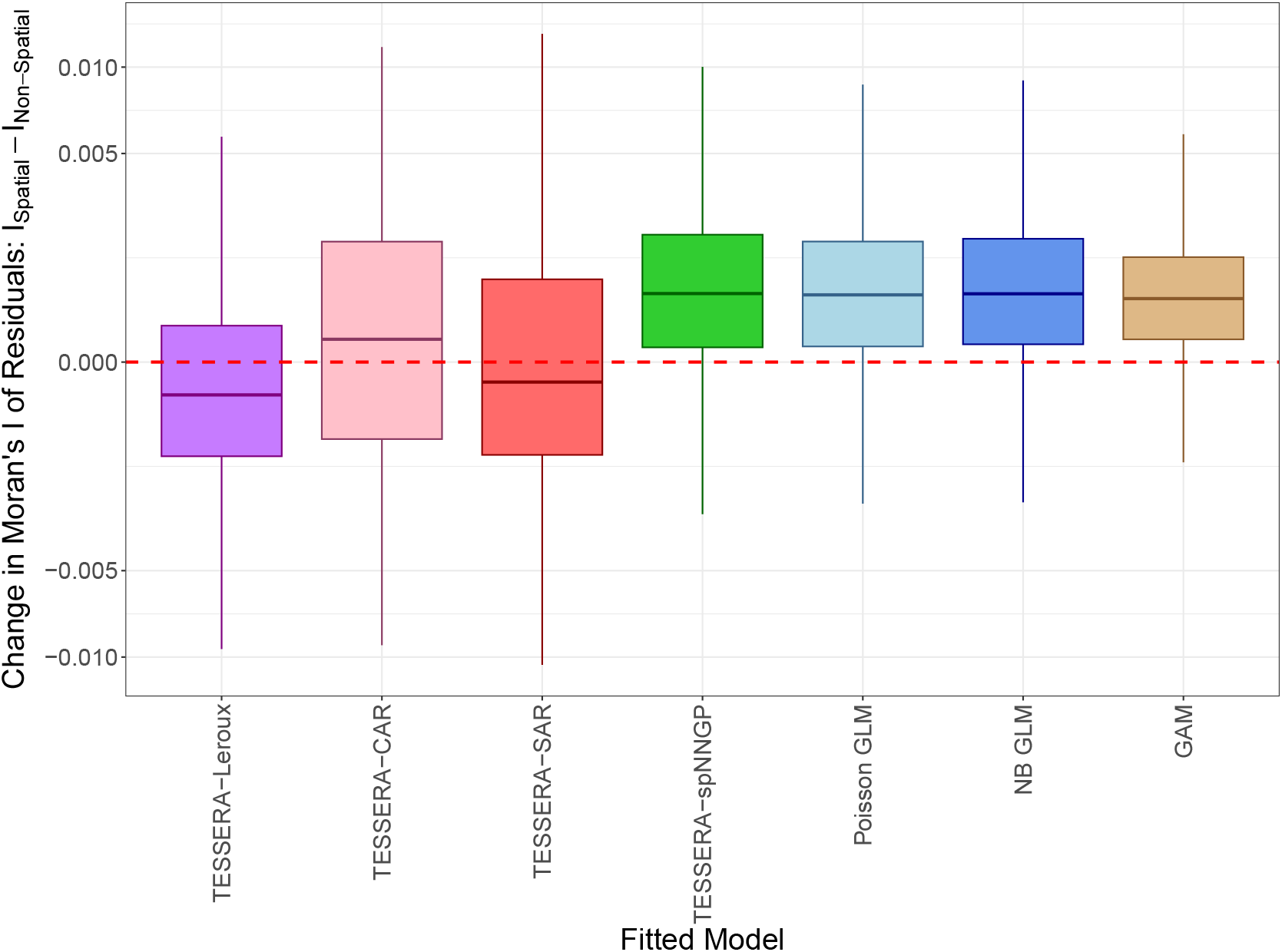
Residual spatial autocorrelation across fitted models (Poisson-Leroux-generated data). To evaluate how effectively each model accounts for spatial autocorrelation, we examine the change in the Moran’s *I* of residuals between spatial and non-spatial generative settings (*I*_*Spatial*_ −*I*_*Non*−*Spatial*_). This differencing accounts for the varying baseline capacities of models to reduce autocorrelation due to general model flexibility and complexity. The boxplots summarize differences in Moran’s *I* over 100 simulation trials, for each of seven methods: `TESSERA` (fit with lattice and spNNGP models), a Poisson GLM, an NB GLM, and a GAM. Values at or below zero (red dashed line) indicate that the method effectively accounts for the spatial autocorrelation present in the data. To generate these results, we produced 100 independent simulation trials for each of two synthetic datasets from a Poisson-Leroux distribution with generative parameters estimated from a multi-sample real-world kidney dataset (Abedini et al., 2024). To calculate *I*_*Spatial*_, we use the parameters as fitted; for *I*_*Non*−*Spatial*_, we set the spatial correlation parameter *γ* to zero so that the generated counts are independent.

**Figure S-7:**
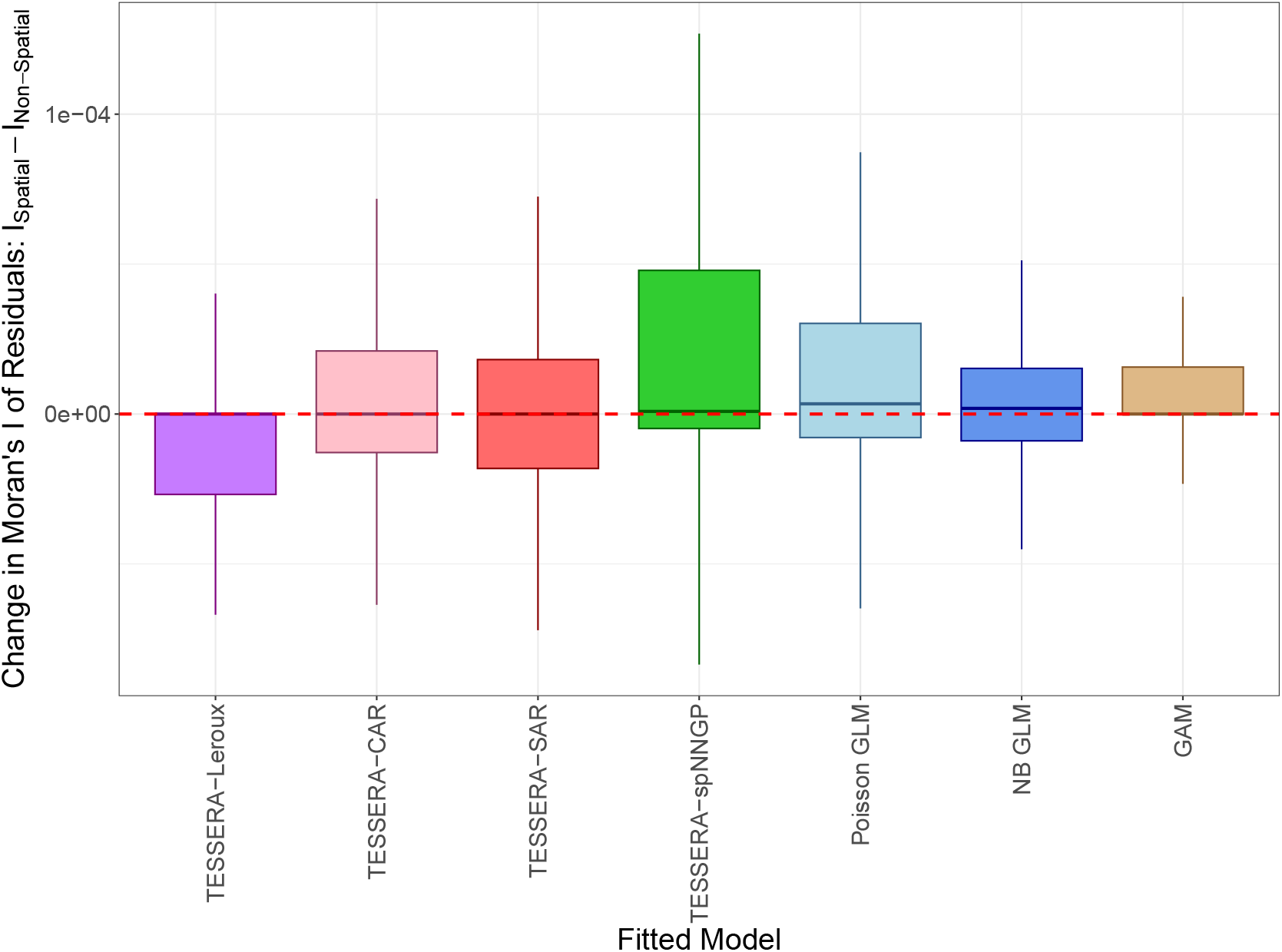
Residual spatial autocorrelation across fitted models (Poisson-spNNGP-generated data). To evaluate how effectively each model accounts for spatial autocorrelation, we examine the change in the Moran’s *I* of residuals between spatial and non-spatial generative settings (*I*_*Spatial*_ −*I*_*Non*−*Spatial*_). This differencing accounts for the varying baseline capacities of models to reduce autocorrelation due to general model flexibility and complexity. The boxplots summarize differences in Moran’s *I* over 100 simulation trials, for each of seven methods: `TESSERA` (fit with lattice and spNNGP models), a Poisson GLM, an NB GLM, and a GAM. Values at or below zero (red dashed line) indicate that the model effectively accounts for the spatial autocorrelation present in the data. To generate these results, we produced 100 independent simulation trials for each of two synthetic datasets from a Poisson-spNNGP distribution with generative parameters estimated from a multi-sample real-world kidney dataset (Abedini et al., 2024). To calculate *I*_*Spatial*_, we use the parameters as fitted; for *I*_*Non*−*Spatial*_, we set the spatial variance parameter (the partial sill) to zero so that the generated counts are independent.

**Figure S-8:**
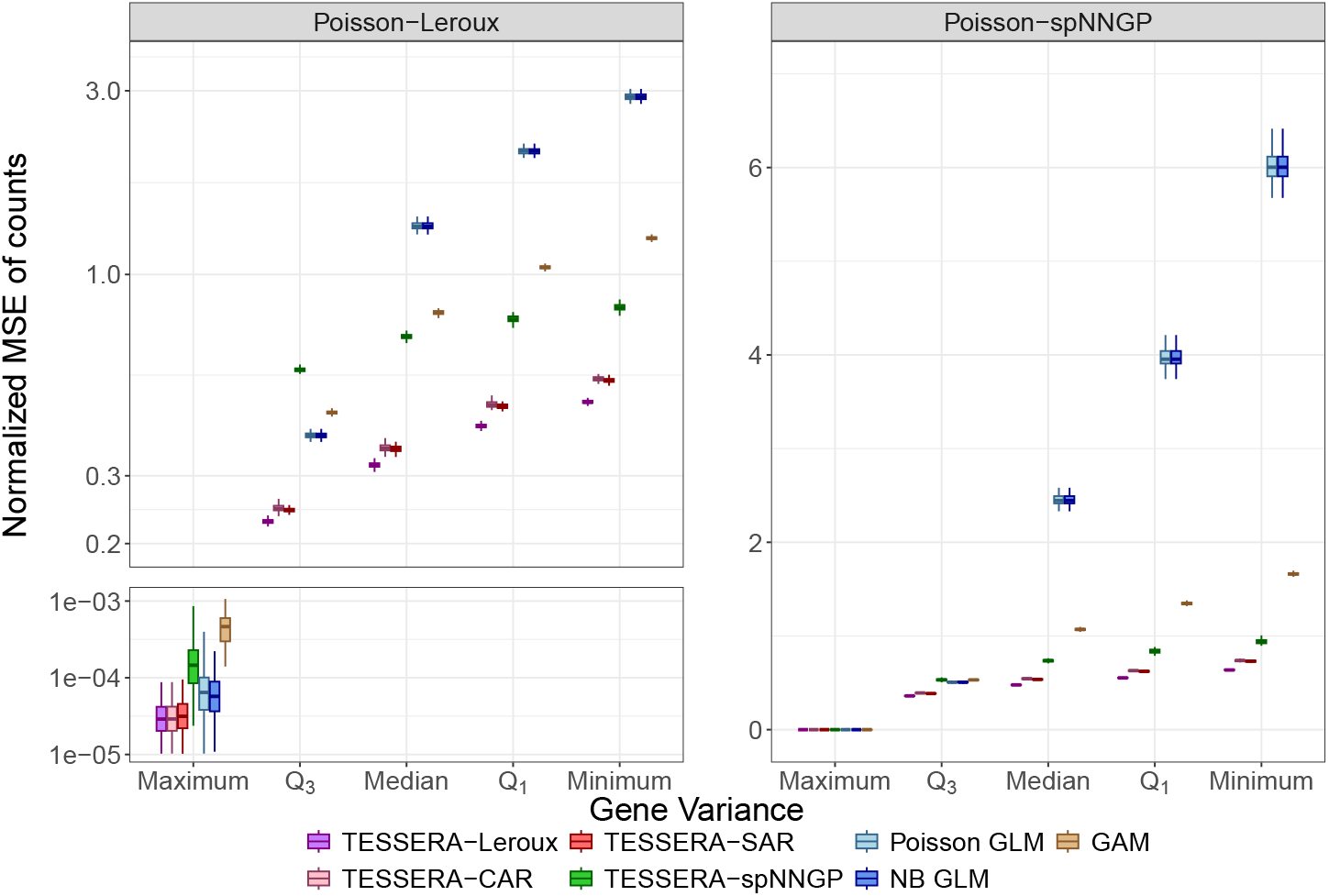
Normalized mean squared error for fitted counts 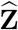. Synthetic data were generated using Poisson-Leroux and Poisson-spNNGP distributions, with generative parameters estimated from a multi-sample real-world kidney dataset (Abedini et al., 2024). We compared `TESSERA` fit with lattice and spNNGP models against a Poisson GLM, an NB GLM, and a GAM. The boxplots summarize the normalized mean squared error (MSE) for the fitted counts, 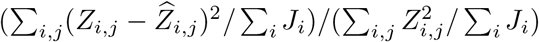, for each method across 100 independent simulation trials. Performance is displayed for a representative subset of genes covering the full distribution of observed expression variance; specifically, genes were selected corresponding to the maximum (*IGKC*), third quartile (*DDX24*), median (*SMTN*), first quartile (*CRLS1*), and minimum (*ERP44*) values of raw count variance across all cells and samples. Facets represent the underlying data-generating models.

**Figure S-9:**
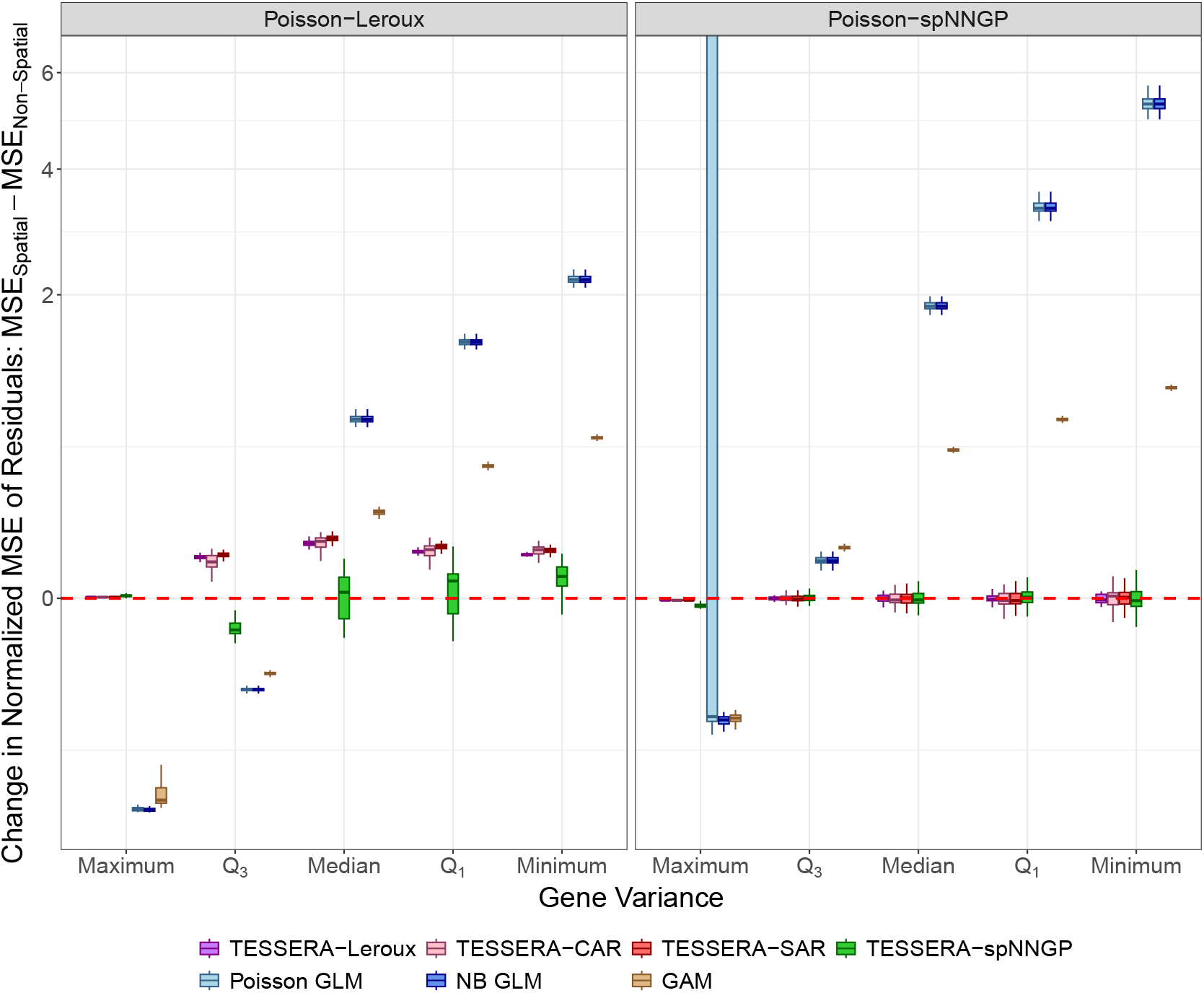
Change in normalized mean squared error for fitted counts 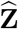. To evaluate how effectively each model accounts for spatial autocorrelation, we examine the change in the normalized MSE of the fitted counts between spatial and non-spatial generative settings (*MSE*_*Spatial*_ −*MSE*_*Non*−*Spatial*_). This differencing accounts for the varying baseline capacities of models to reduce normalized MSE due to general model flexibility and complexity. The boxplots summarize differences in normalized MSE over 100 simulation trials, for each of seven methods: `TESSERA` (fit with lattice and spNNGP models), a Poisson GLM, an NB GLM, and a GAM and for each of five representative genes covering the full distribution of observed expression variance. Specifically, genes were selected corresponding to the maximum (*IGKC*), third quartile (*DDX24*), median (*SMTN*), first quartile (*CRLS1*), and minimum (*ERP44*) values of raw count variance across all cells and samples. Values at or close to zero (red dashed line) indicate that the model is equally effective in both settings. To generate these results, we produced 100 independent simulation trials for each of two synthetic datasets from either a Poisson-Leroux or Poisson-spNNGP distribution with generative parameters estimated from a multi-sample real-world kidney dataset (Abedini et al., 2024). To calculate *MSE*_*Spatial*_, we use the parameters as fitted; for *MSE*_*Non*−*Spatial*_, we set the spatial component (*γ* for lattice models or the partial sill for an spNNGP model) to zero to ensure the generated counts are independent.

#### S7.3. Synthetic Data: Comparison of TESSERA with Single-Sample Methods

**Figure S-10:**
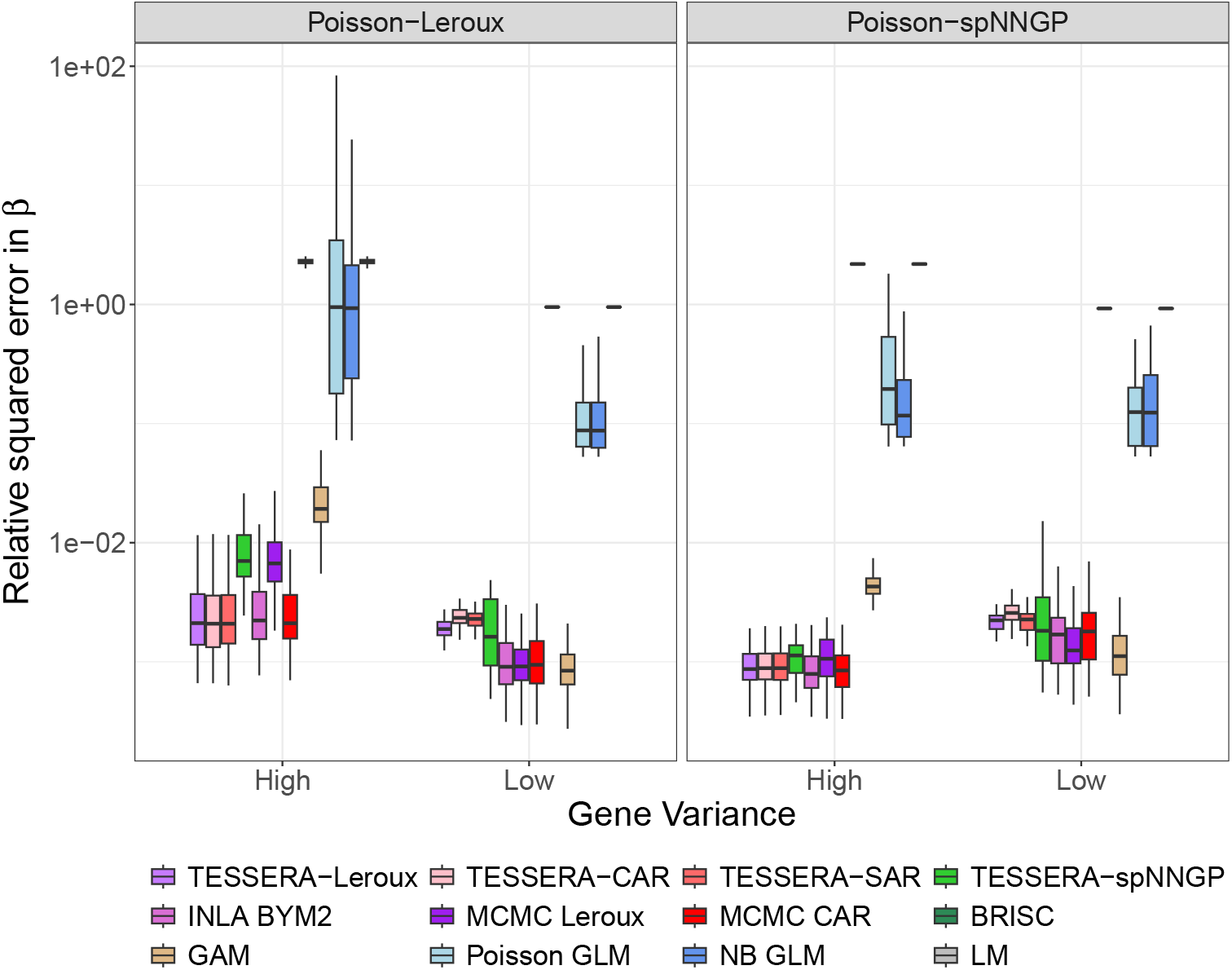
Relative squared error in *β* in single-sample setting (excluding low-abundance cell types). Synthetic data were generated using Poisson-Leroux and Poisson-spNNGP distributions with respective generative parameters estimated on a multi-sample real-world kidney dataset (Abedini et al., 2024). Estimation methods were evaluated on a single sample (corresponding to the *HK3035_ST* sample from the kidney dataset). Low-abundance cell types (those with ≤ 3 cells) were **excluded** in this set of simulations. We compared `TESSERA` fit with lattice and spNNGP models against existing spatial methods (MCMC-based Leroux/CAR, INLA BYM2, and BRISC), with a Poisson GLM, an NB GLM, a GAM, and a linear model (LM) serving as baselines. Estimation accuracy of the fixed effects ***β*** was assessed via the relative squared error, 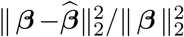, calculated across 100 independent simulation trials. Performance is shown for genes with the maximum and minimum variances of the raw counts across all cells and samples (denoted by ‘High’ and ‘Low’, respectively). Facets represent the underlying data-generating models.

**Figure S-11:**
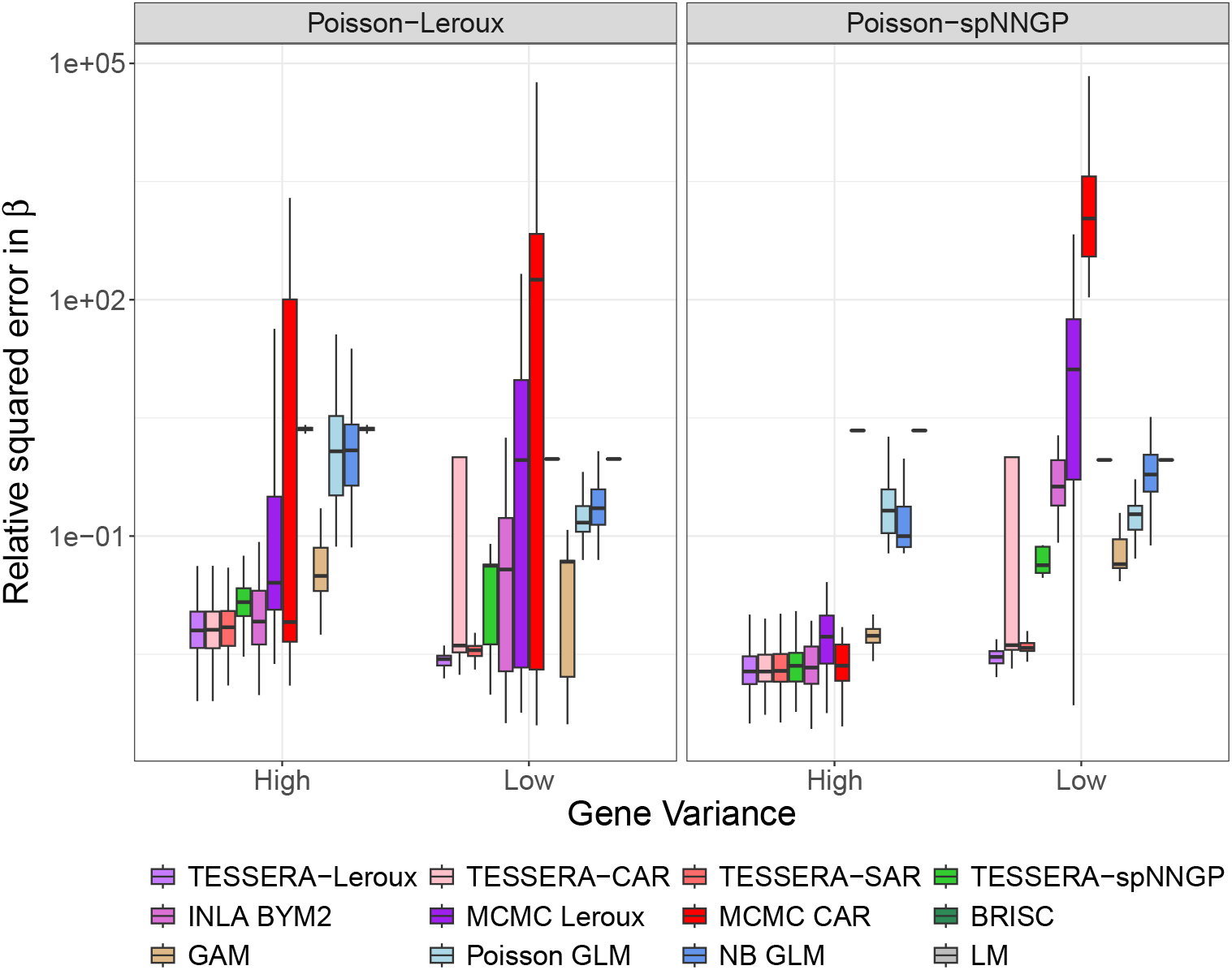
Relative squared error in *β* in single-sample setting (including low-abundance cell types). Synthetic data were generated using Poisson-Leroux and Poisson-spNNGP distributions, with generative parameters estimated from a multi-sample real-world kidney dataset (Abedini et al., 2024). Estimation methods were evaluated on a single sample (corresponding to sample *HK3035_ST* from the kidney dataset), where low-abundance cell types (≤3 cells) were **included** to assess model robustness to small cell groups representing high-variance elements of ***β***. We compared `TESSERA` with lattice and spNNGP models against existing spatial methods (MCMC-based Leroux/CAR, INLA BYM2, and BRISC), with a Poisson GLM, an NB GLM, a GAM, and a linear model (LM) serving as baselines. Estimation accuracy of the fixed effects ***β*** was assessed via the relative squared error, 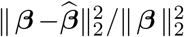, calculated across 100 independent simulation trials. Performance is shown for genes with the maximum and minimum variances of raw counts across all cells and samples (denoted as ‘High’ and ‘Low’, respectively). Facets represent the underlying data-generating models.

#### S7.4. Synthetic Data: Power Simulation

**Figure S-12:**
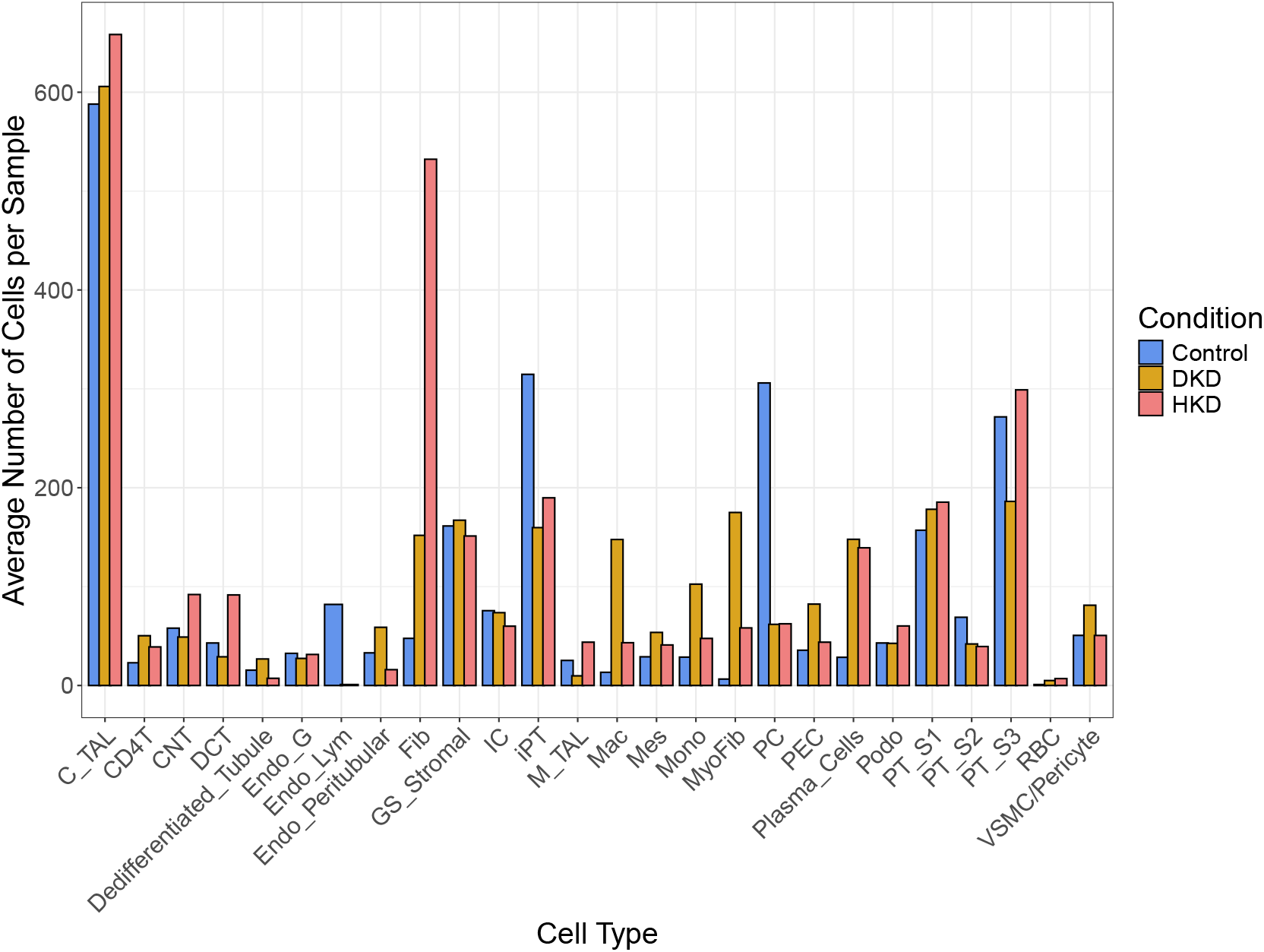
Average number of cells per sample by cell type and condition. Shown are average cell abundance by cell type and condition for a multi-sample real-world kidney dataset (Abedini et al., 2024). Bars represent the mean number of cells per sample for each cell type-condition pair.

**Figure S-13:**
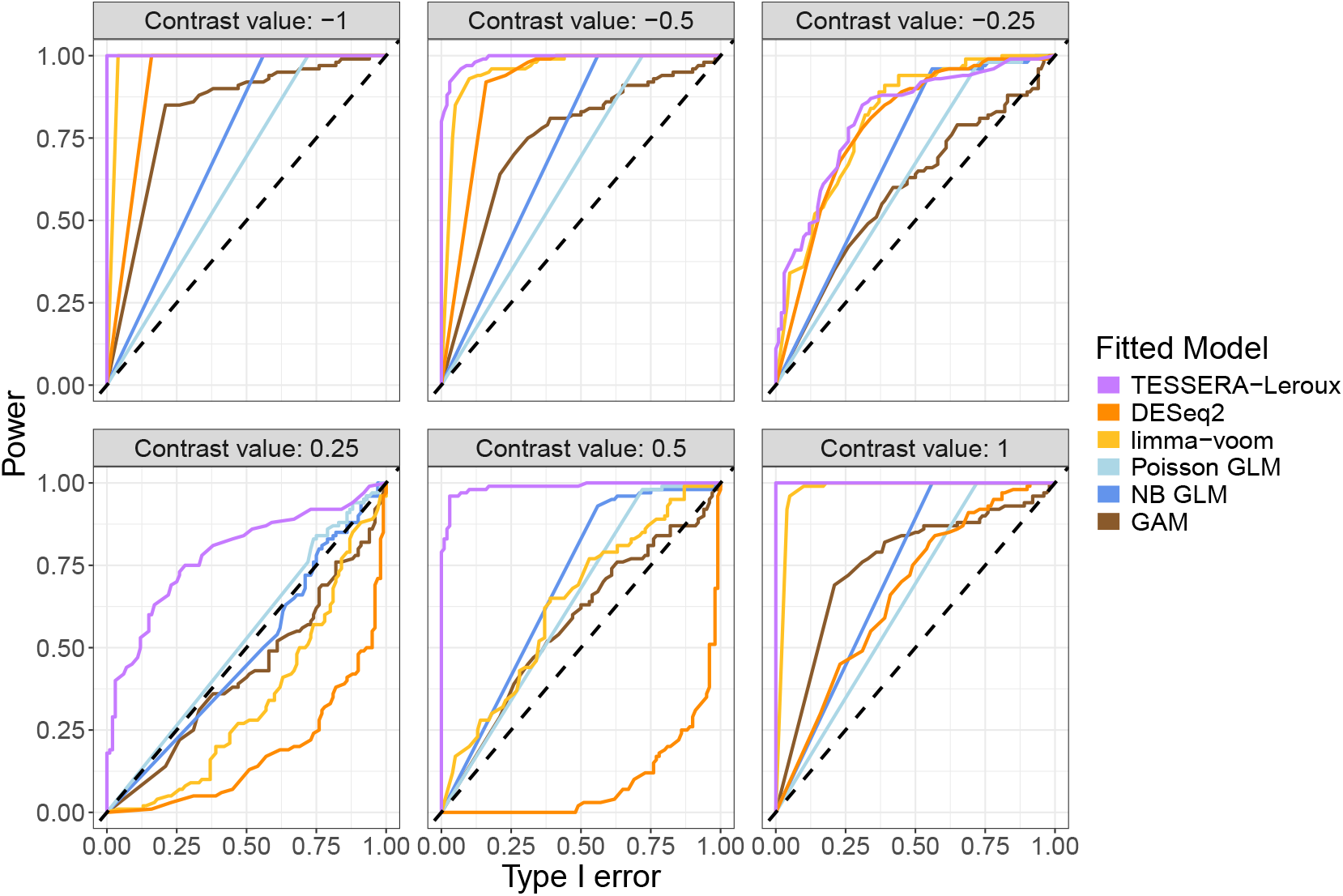
ROC curves for Poisson-Leroux-generated data. This figure evaluates the trade-off between power and Type I error rate for detecting DE between conditions for the *C_TAL* cell type across a range of effect sizes. Synthetic data were generated using a Poisson-Leroux distribution, with generative parameters estimated from the *IGHG2* gene in a multi-sample real-world kidney dataset (Abedini et al., 2024). For each panel shown, we modify two entries of the fixed effects vector ***β***, corresponding to the Control and the DKD conditions, to generate a contrast of specified size. We set the control coefficient to be the average of all coefficients and the DKD coefficient to be the control coefficient plus a given effect size. We then fit the `TESSERA` Poisson-Leroux model as well as a Poisson GLM, an NB GLM, and a GAM to the data and perform a Wald test to see if the contrast is detectable. For comparison, we also evaluate pseudobulk methods `DESeq2` and `limma-voom`. We present ROC curves of power (y-axis) as a function of the Type I error rate (x-axis) across 100 independent simulation trials for each contrast value. The dotted diagonal line represents the *y* = *x* identity line, where the power is exactly equal to the Type I error rate. Curves that rise more steeply toward the top-left corner indicate superior method performance, with `TESSERA` (fit with a Poisson-Leroux model) consistently showing comparable or superior performance to other methods across all contrast values.

**Figure S-14:**
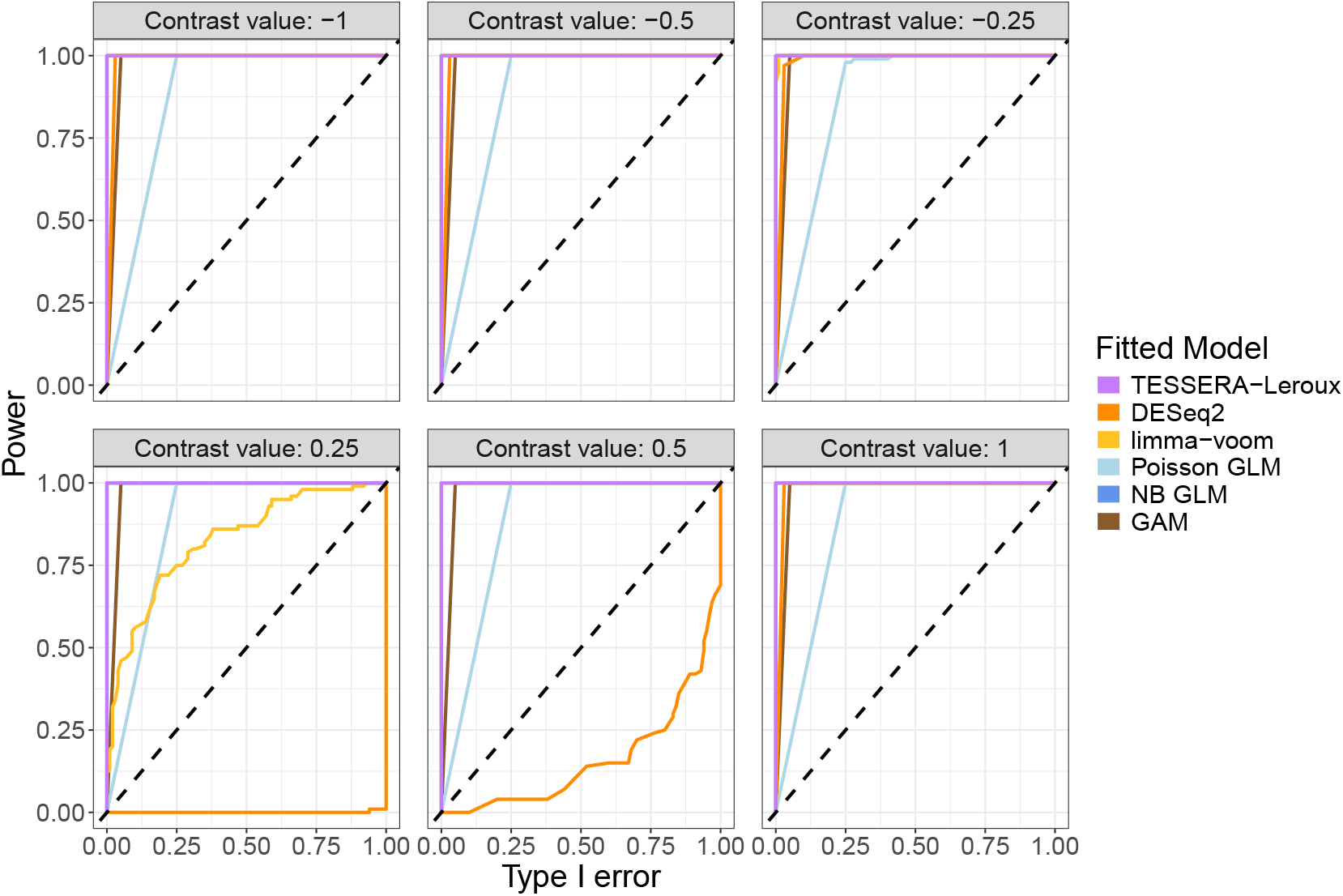
ROC curves for Poisson-spNNGP-generated data. This figure evaluates the trade-off between power and Type I error rate for detecting DE between conditions for the *C_TAL* cell type across a range of effect sizes. Synthetic data were generated using a Poisson-spNNGP distribution, with generative parameters estimated from the *IGHG2* gene in a multi-sample real-world kidney dataset (Abedini et al., 2024). For each panel shown, we modify two entries of the fixed effects vector ***β***, corresponding to the Control and the DKD conditions, to generate a contrast of specified size. We set the control coefficient to be the average of all coefficients and the DKD coefficient to be the control coefficient plus a given effect size. We then fit the `TESSERA` Poisson-Leroux model as well as a Poisson GLM, an NB GLM, and a GAM to the data and perform a Wald test to see if the contrast is detectable. For comparison, we also evaluate pseudobulk methods `DESeq2` and `limma-voom`. We present ROC curves of power (y-axis) as a function of the Type I error rate (x-axis) across 100 independent simulation trials for each contrast value. The dotted diagonal line represents the *y* = *x* identity line, where the power is exactly equal to the Type I error rate. Curves that rise more steeply toward the top-left corner indicate superior model performance, with `TESSERA` (fit with a Poisson-Leroux model) consistently showing comparable or superior performance to other methods across all contrast values.

**Figure S-15:**
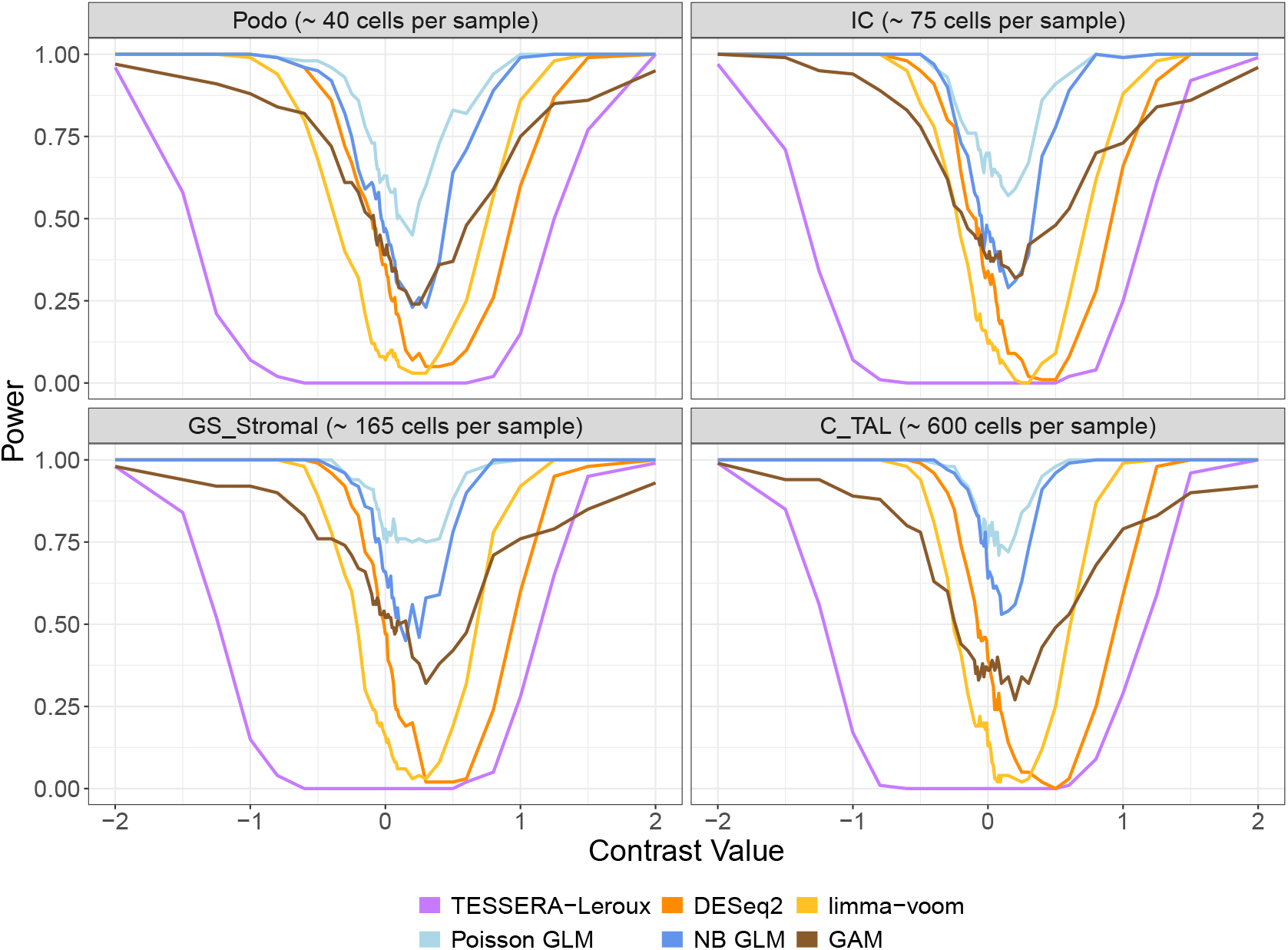
Power for Poisson-Leroux-generated data. This figure examines power for detecting DE between conditions within a cell type for one gene. It presents plots of power (y-axis) as a function of the contrast value (x-axis), where power is estimated by the proportion of 100 independent simulation trials in which the null hypothesis of no DE is rejected at nominal Type I error rate of 0.05. At an effect size of 0, this value represents the Type I error rate. Synthetic data were generated using a Poisson-Leroux distribution, with generative parameters estimated from the *IGHG2* gene in a multi-sample real-world kidney dataset (Abedini et al., 2024). For each cell type shown, we modify two entries of the fixed effects vector ***β***, corresponding to the Control and the DKD conditions, to generate a contrast of specified size. We set the control coefficient to be the average of all coefficients and the DKD coefficient to be the control coefficient plus a given effect size. We then fit the `TESSERA` Poisson-Leroux model as well as a Poisson GLM, an NB GLM, and a GAM to the data and perform a Wald test to see if the contrast is detectable. For comparison, we also evaluate pseudobulk methods `DESeq2` and `limma-voom`.

**Figure S-16:**
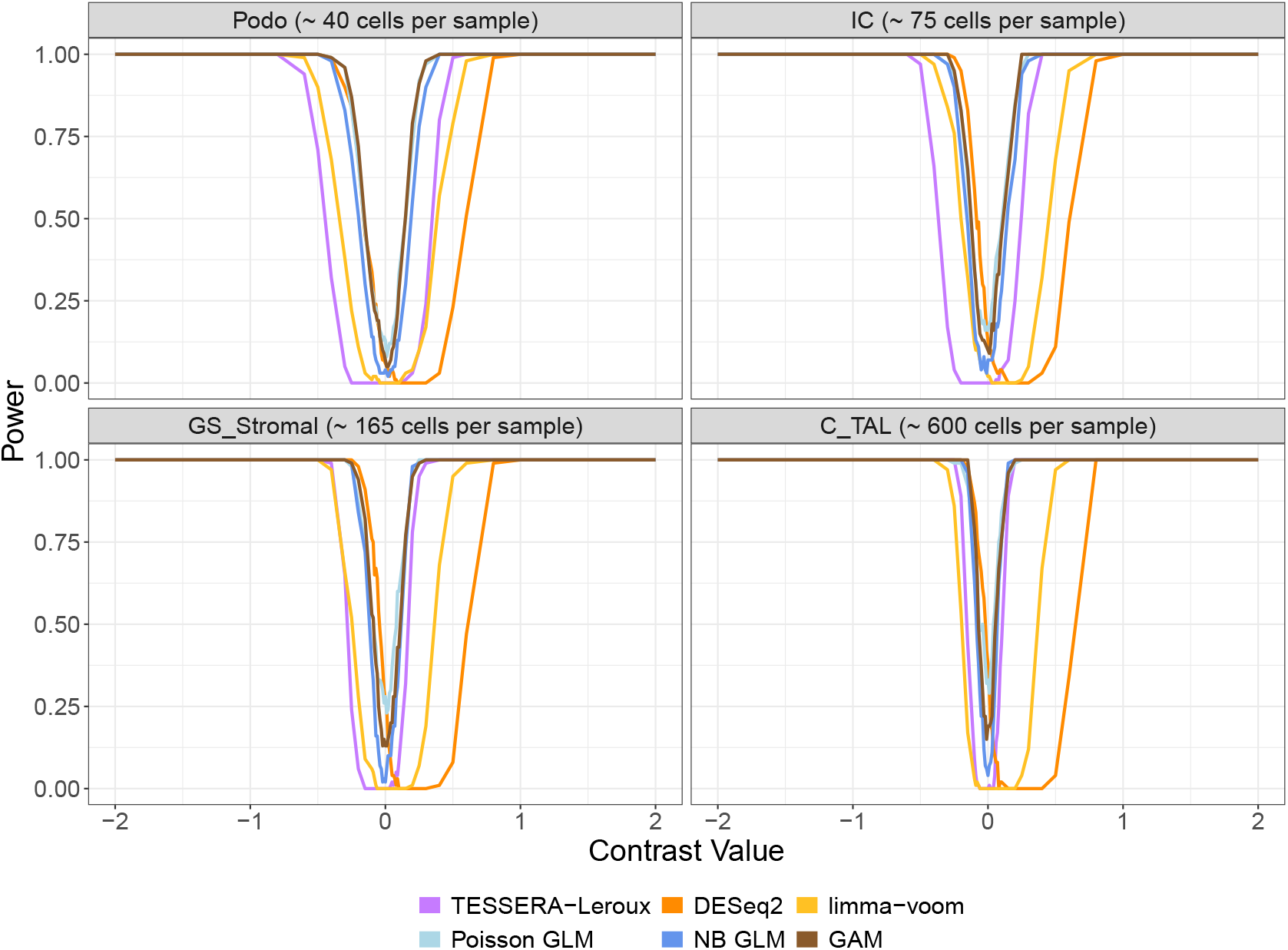
Power for Poisson-spNNGP-generated data. This figure examines power for detecting DE between conditions within a cell type for one gene. It presents plots of power (y-axis) as a function of the contrast value (x-axis), where power is estimated by the proportion of 100 independent simulation trials in which the null hypothesis of no DE is rejected at nominal Type I error rate of 0.05. At an effect size of 0, this value represents the Type I error rate. Synthetic data were generated using a Poisson-spNNGP distribution, with generative parameters estimated from the *IGHG2* gene in a multi-sample real-world kidney dataset (Abedini et al., 2024). For each cell type shown, we modify two entries of the fixed effects vector ***β***, corresponding to the Control and the DKD conditions to generate a contrast of specified size. We set the control coefficient to be the average of all coefficients and the DKD coefficient to be the control coefficient plus a given effect size. We then fit the `TESSERA` Poisson-Leroux model as well as a Poisson GLM, an NB GLM, and a GAM to the data and perform a Wald test to see if the contrast is detectable. For comparison, we also evaluate pseudobulk methods `DESeq2` and `limma-voom`.

**Figure S-17:**
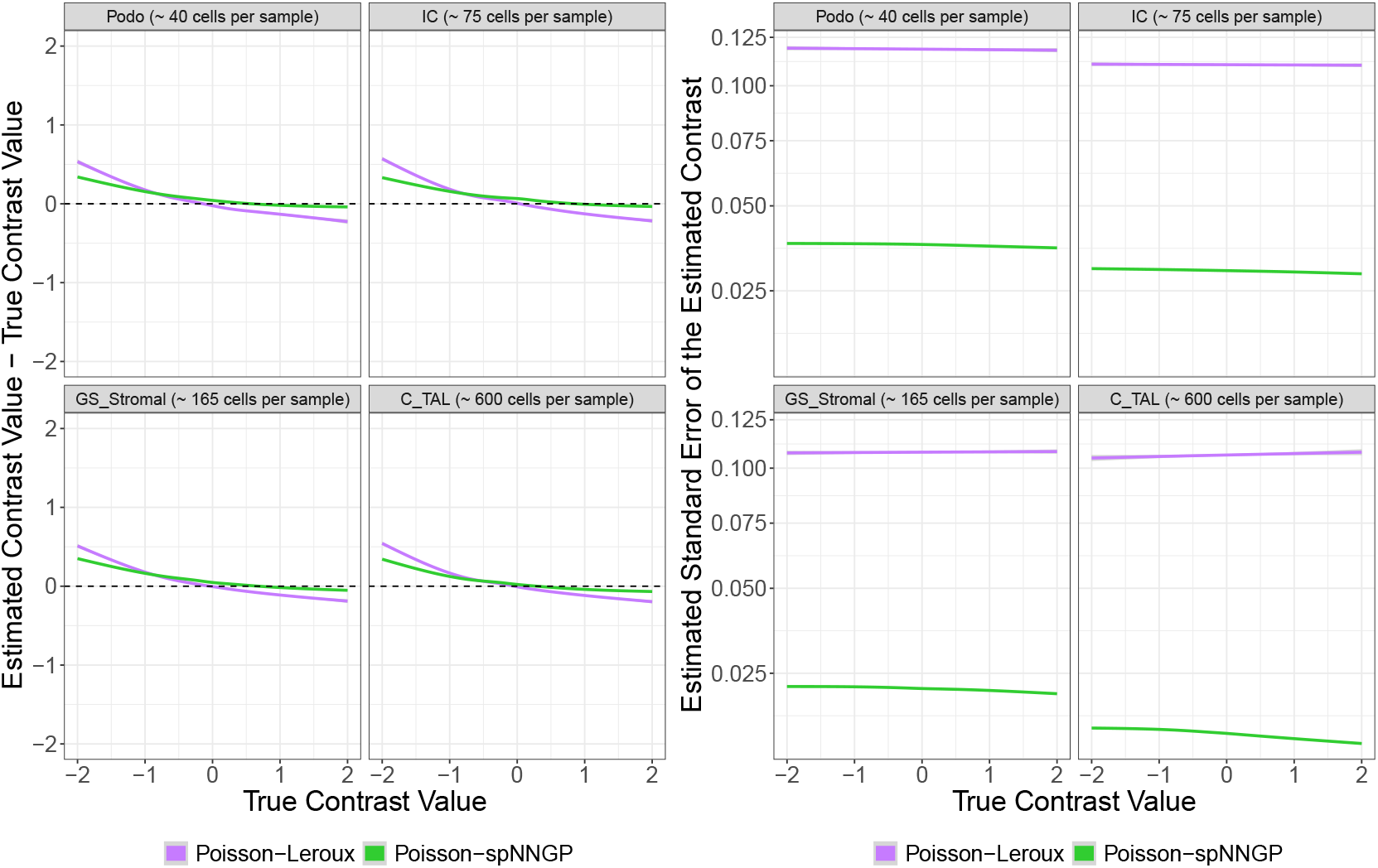
Estimated contrast value and standard error of the estimated contrast for Poisson-Leroux and Poisson-spNNGP-generated data. Synthetic data were generated using a Poisson-Leroux or Poisson-spNNGP distribution, with generative parameters estimated from the *IGHG2* gene in a multi-sample real-world kidney dataset (Abedini et al., 2024). For each cell type shown, we modify two entries of the fixed effects vector ***β***, corresponding to the Control and the DKD conditions to generate a contrast of specified size. We set the control coefficient to be the average of all coefficients and the DKD coefficient to be the control coefficient plus a given effect size. We then fit the `TESSERA` Poisson-Leroux model to the data and perform a Wald test to see if the contrast is detectable. Across 100 independent simulation trials for each contrast and contrast value, we plot performance metrics against the true contrast value. The left-hand plot shows the difference between the average of the estimated contrasts and the true contrast values, while the right-hand plot displays the average estimated standard error of the estimated contrast.

**Figure S-18:**
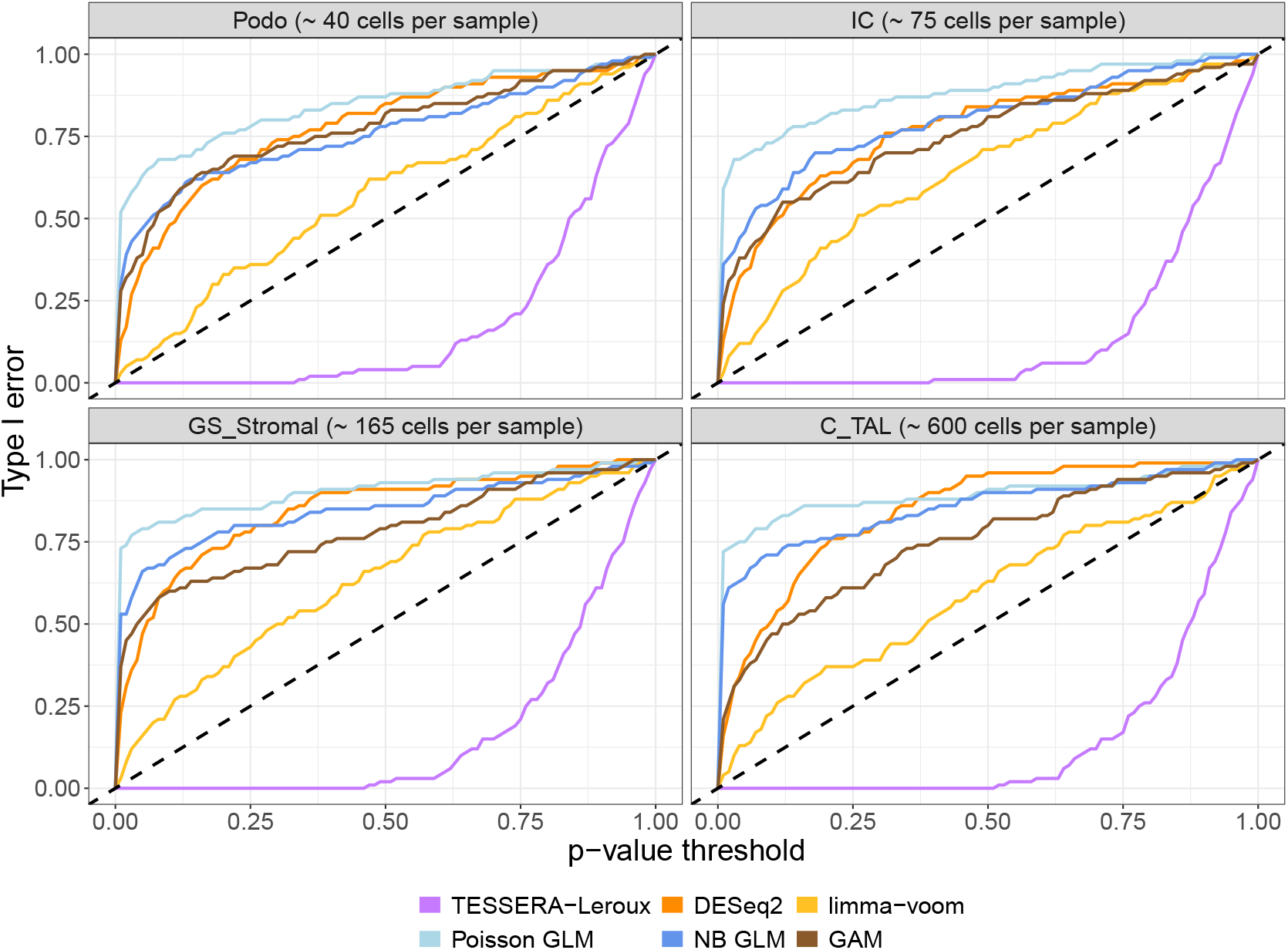
Type I error rate for Poisson-Leroux-generated data. Synthetic data were generated using a Poisson-Leroux distribution, with generative parameters estimated from the *IGHG2* gene in a multi-sample real-world kidney dataset (Abedini et al., 2024). For each cell type shown, we modify two entries of the fixed effects vector ***β***, corresponding to the Control and the DKD conditions, to generate a contrast of specified size. We set the control coefficient to be the average of all coefficients and the DKD coefficient to be the control coefficient plus a given effect size. We then fit the `TESSERA` Poisson-Leroux model as well as a Poisson GLM, an NB GLM, and a GAM to the data and perform a Wald test to see if the contrast is detectable. For comparison, we also evaluate pseudobulk methods `DESeq2` and `limma-voom`. We present curves of the Type I error rate as a function of the *p*-value threshold across 100 independent simulation trials for each contrast and effect size. Exact control of the Type I error rate is represented by the dotted line. Methods with curves exceeding the dotted line fail to control the Type I error rate, with `TESSERA` being the only method that achieves error control by remaining consistently below the threshold.

**Figure S-19:**
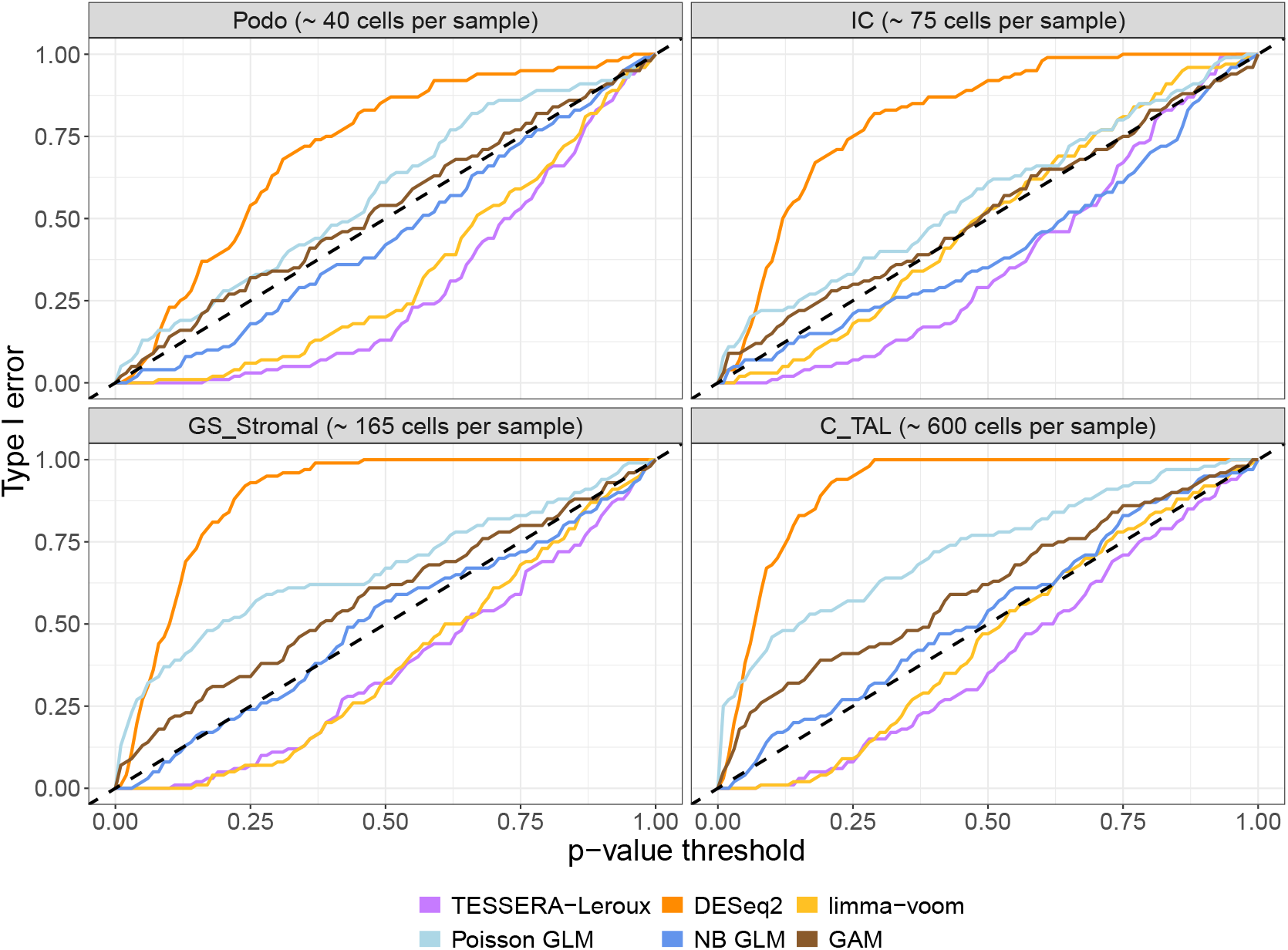
Type I error rate for Poisson-spNNGP-generated data. Synthetic data were generated using a Poisson-spNNGP distribution, with generative parameters estimated from the *IGHG2* gene in a multi-sample real-world kidney dataset (Abedini et al., 2024). For each cell type shown, we modify two entries of the fixed effects vector ***β***, corresponding to the Control and the DKD conditions to generate a contrast of specified size. We set the control coefficient to be the average of all coefficients and the DKD coefficient to be the control coefficient plus a given effect size. We then fit the `TESSERA` Poisson-Leroux model as well as a Poisson GLM, an NB GLM, and a GAM to the data and perform a Wald test to see if the contrast is detectable. For comparison, we also evaluate pseudobulk methods `DESeq2` and `limma-voom`. We present curves of the Type I error rate as a function of the *p*-value threshold across 100 independent simulation trials for each contrast and effect size. Exact control of the Type I error rate is represented by the dotted line. Methods with curves exceeding the dotted line fail to control the Type I error rate, with `TESSERA` being the only method that achieves error control by remaining consistently below the threshold.

**Figure S-20:**
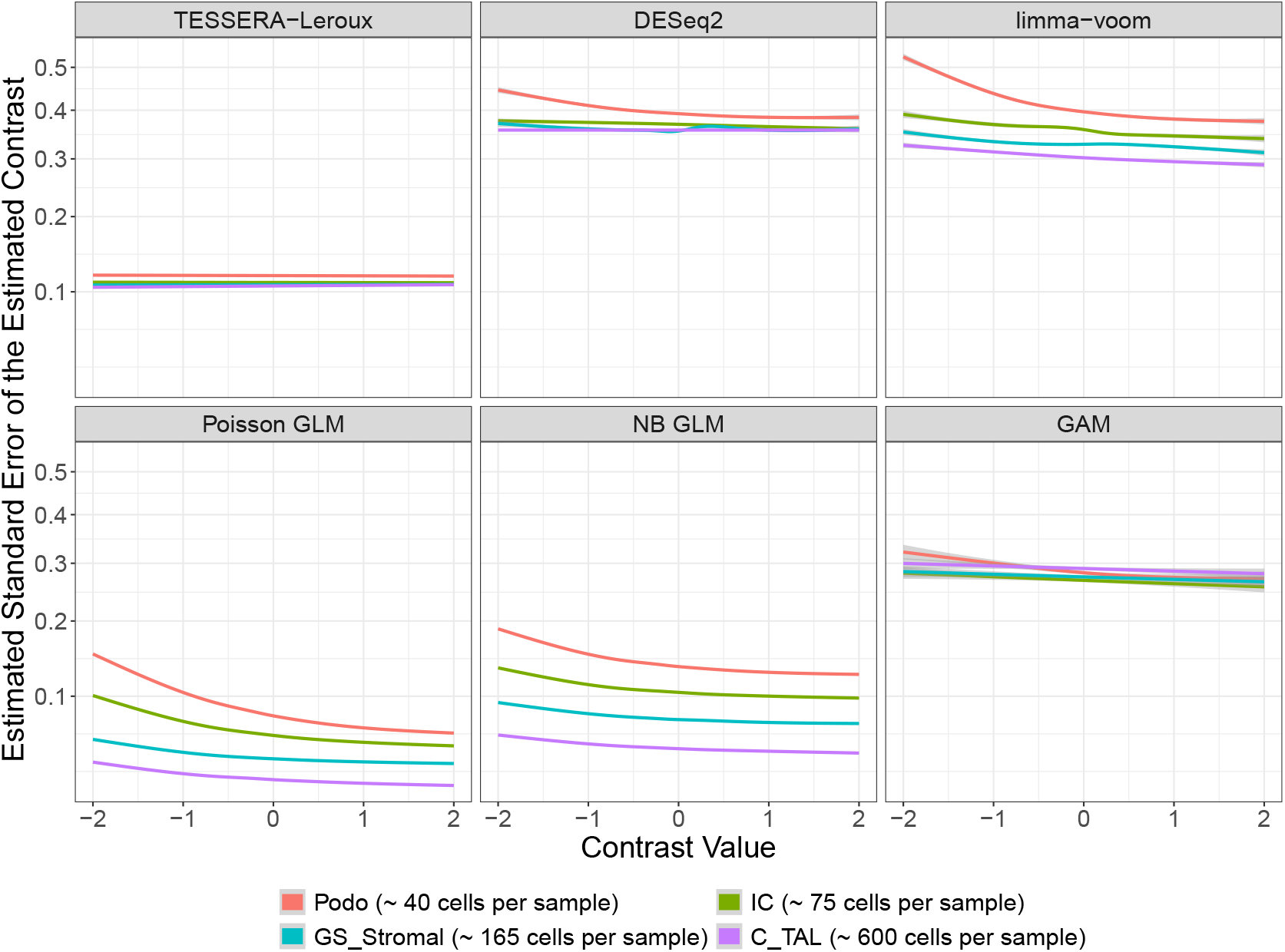
Estimated standard error of the estimated contrast for Poisson-Leroux-generated data. Synthetic data were generated using a Poisson-Leroux distribution, with generative parameters estimated from the *IGHG2* gene in a multi-sample real-world kidney dataset (Abedini et al., 2024). For each cell type shown, we modify two entries of the fixed effects vector ***β***, corresponding to the Control and the DKD conditions to generate a contrast of specified size. We set the control coefficient to be the average of all coefficients and the DKD coefficient to be the control coefficient plus a given effect size. We then fit the `TESSERA` Poisson-Leroux model as well as a Poisson GLM, an NB GLM, and a GAM to the data and perform a Wald test to see if the contrast is detectable. For comparison, we also evaluate pseudobulk methods `DESeq2` and `limma-voom`. Across 100 independent simulation trials, for each contrast and contrast value, we plot the average estimated standard error of the estimated contrast against the true contrast value.

**Figure S-21:**
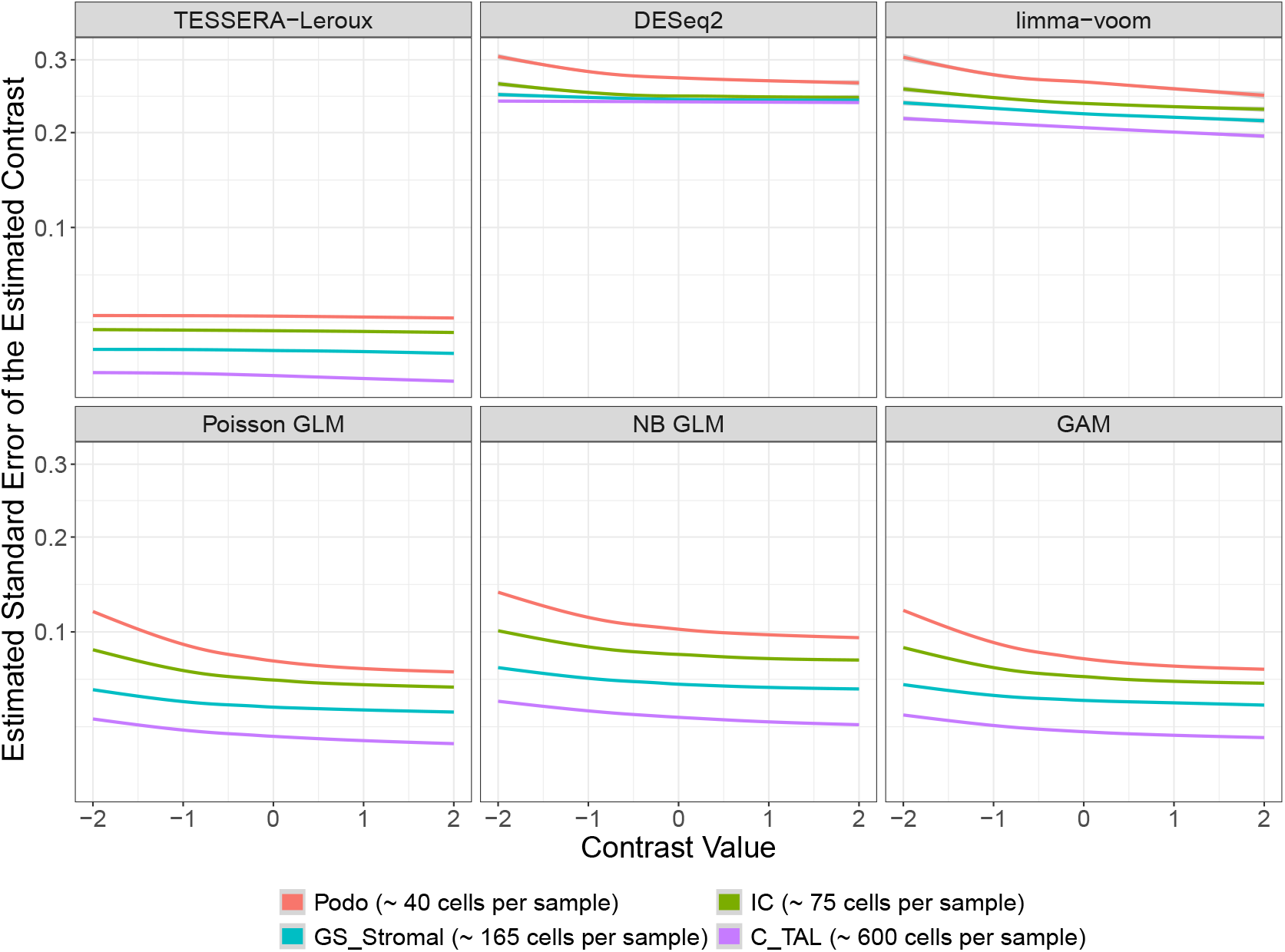
Estimated standard error of the estimated contrast for Poisson-spNNGP-generated data. Synthetic data were generated using a Poisson-spNNGP distribution, with generative parameters estimated from the *IGHG2* gene in a multi-sample real-world kidney dataset (Abedini et al., 2024). For each cell type shown, we modify two entries of the fixed effects vector ***β***, corresponding to the Control and the DKD conditions to generate a contrast of specified size. We set the control coefficient to be the average of all coefficients and the DKD coefficient to be the control coefficient plus a given effect size. We then fit the `TESSERA` Poisson-Leroux model as well as a Poisson GLM, an NB GLM, and a GAM to the data and perform a Wald test to see if the contrast is detectable. For comparison, we also evaluate pseudobulk methods `DESeq2` and `limma-voom`. Across 100 independent simulation trials, for each contrast and contrast value, we plot the average estimated standard error of the estimated contrast against the true contrast value.

**Figure S-22:**
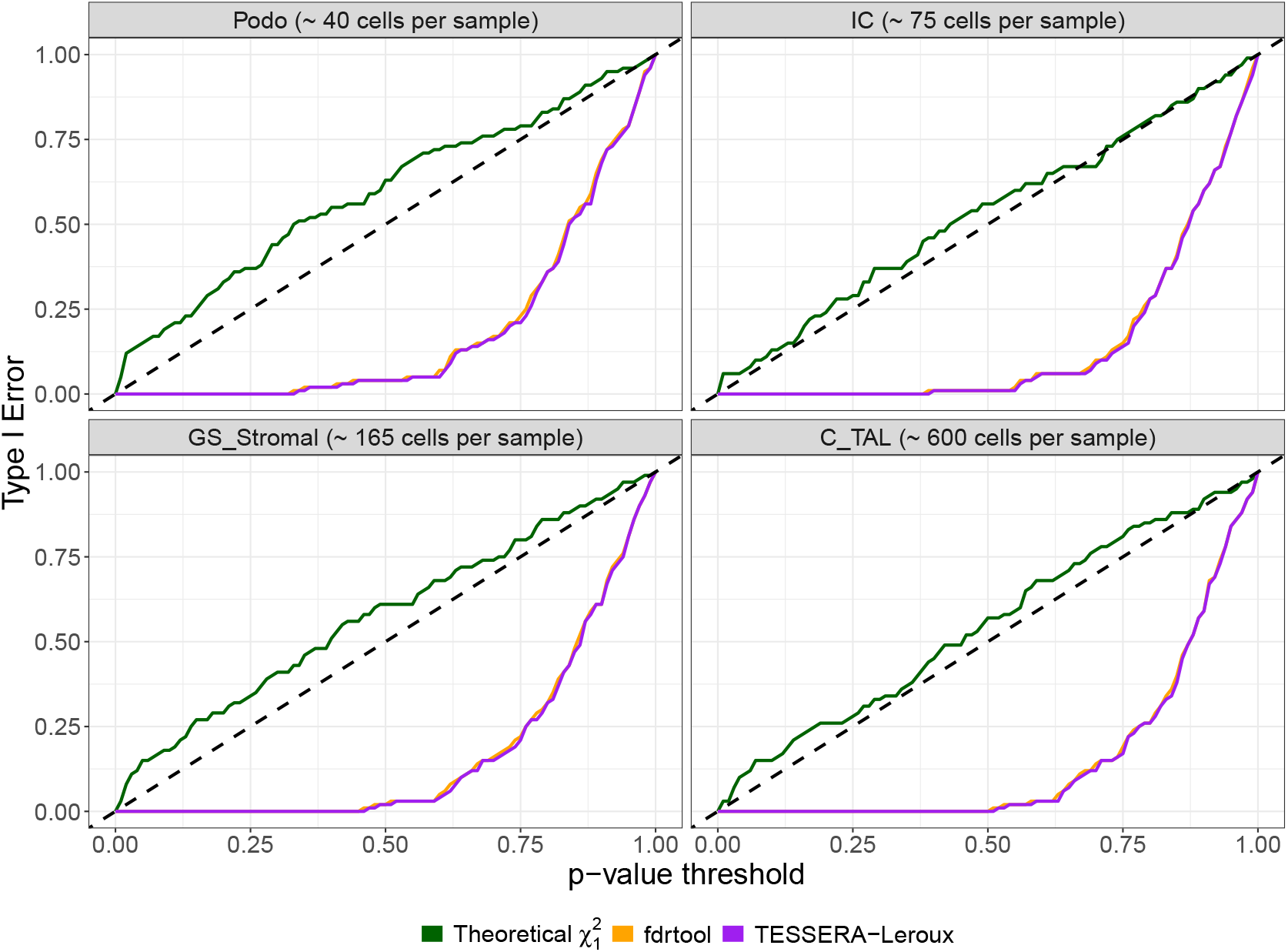
Type I error rate of empirical null distribution estimation procedures and theoretical 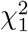 baseline in Poisson-Leroux-generated data. Synthetic data were generated using a Poisson-Leroux distribution, with generative parameters estimated from the *IGHG2* gene in a multi-sample real-world kidney dataset (Abedini et al., 2024). For each cell type shown, we modify two entries of the fixed effects vector ***β***, corresponding to the Control and the DKD conditions to generate a contrast of specified size. We set the control coefficient to be the average of all coefficients and the DKD coefficient to be the control coefficient plus a given effect size. We then fit the `TESSERA` Poisson-Leroux model and evaluate the detectability of the contrasts by comparing two empirical null distribution estimation procedures, the one used by `TESSERA` (Section 2.4) and the default `fdrtool` (Strimmer, 2008) implementation, against a theoretical 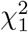 baseline. We present curves of the Type I error rate as a function of the *p*-value threshold across 100 independent simulation trials for each contrast and effect size. Exact control of the Type I error rate is represented by the dotted line. Methods with curves exceeding the dotted line fail to control the Type I error rate, with `TESSERA` being the only method that achieves error control by remaining consistently below the threshold.

**Figure S-23:**
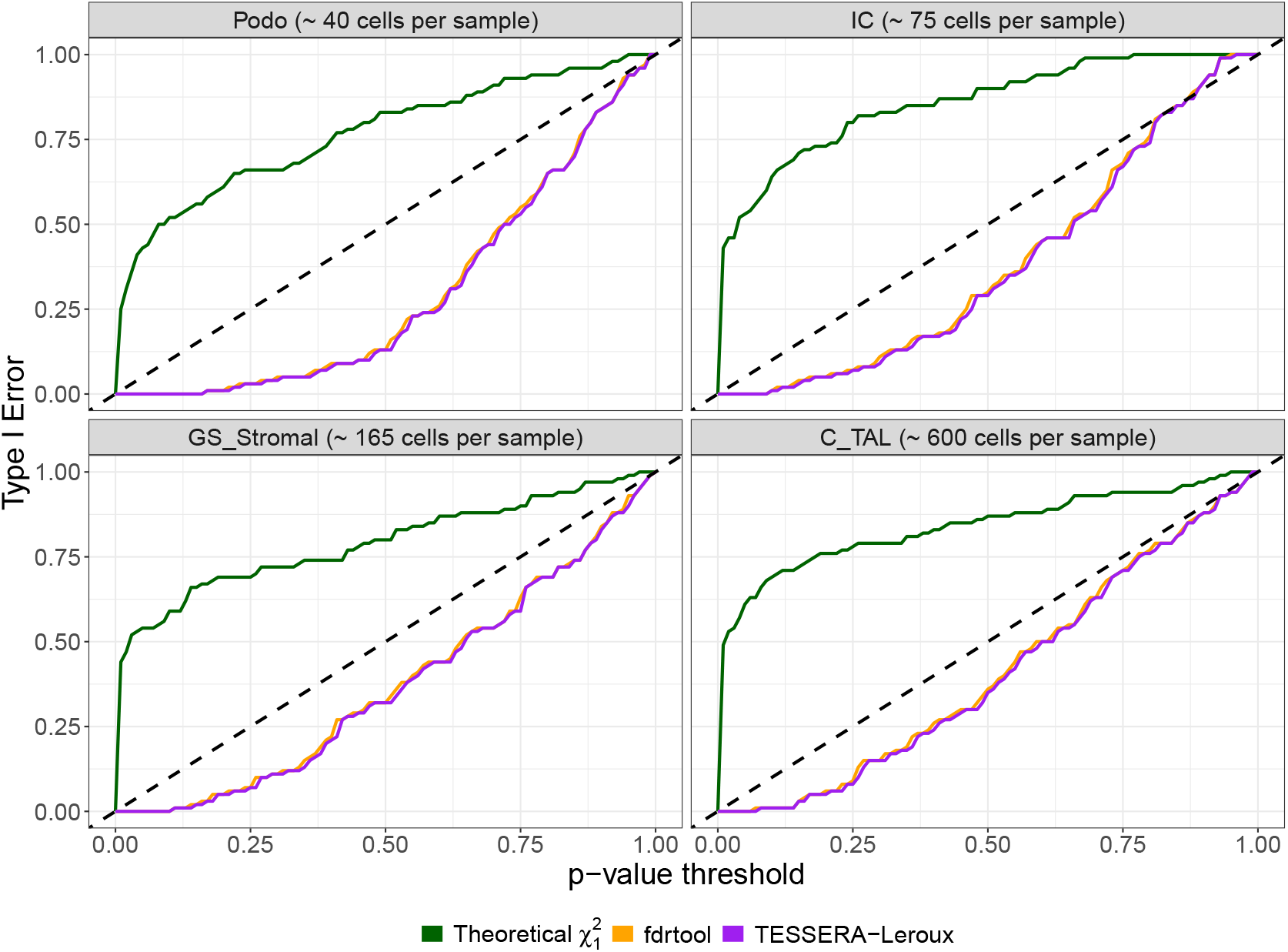
Type I error rate of empirical null distribution estimation procedures and theoretical 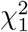 baseline in Poisson-spNNGP-generated data. Synthetic data were generated using a Poisson-spNNGP distribution, with generative parameters estimated from the *IGHG2* gene in a multi-sample real-world kidney dataset (Abedini et al., 2024). For each cell type shown, we modify two entries of the fixed effects vector ***β***, corresponding to the Control and the DKD conditions to generate a contrast of specified size. We set the control coefficient to be the average of all coefficients and the DKD coefficient to be the control coefficient plus a given effect size. We then fit the `TESSERA` Poisson-Leroux model and evaluate the detectability of the contrasts by comparing two empirical null distribution estimation procedures, the one used by `TESSERA` (Section 2.4) and the default `fdrtool` (Strimmer, 2008) implementation, against a theoretical 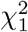 baseline. We present curves of the Type I error rate as a function of the *p*-value threshold across 100 independent simulation trials for each contrast and effect size. Exact control of the Type I error rate is represented by the dotted line. Methods with curves exceeding the dotted line fail to control the Type I error rate, with `TESSERA` being the only method that achieves error control by remaining consistently below the threshold.

**Figure S-24:**
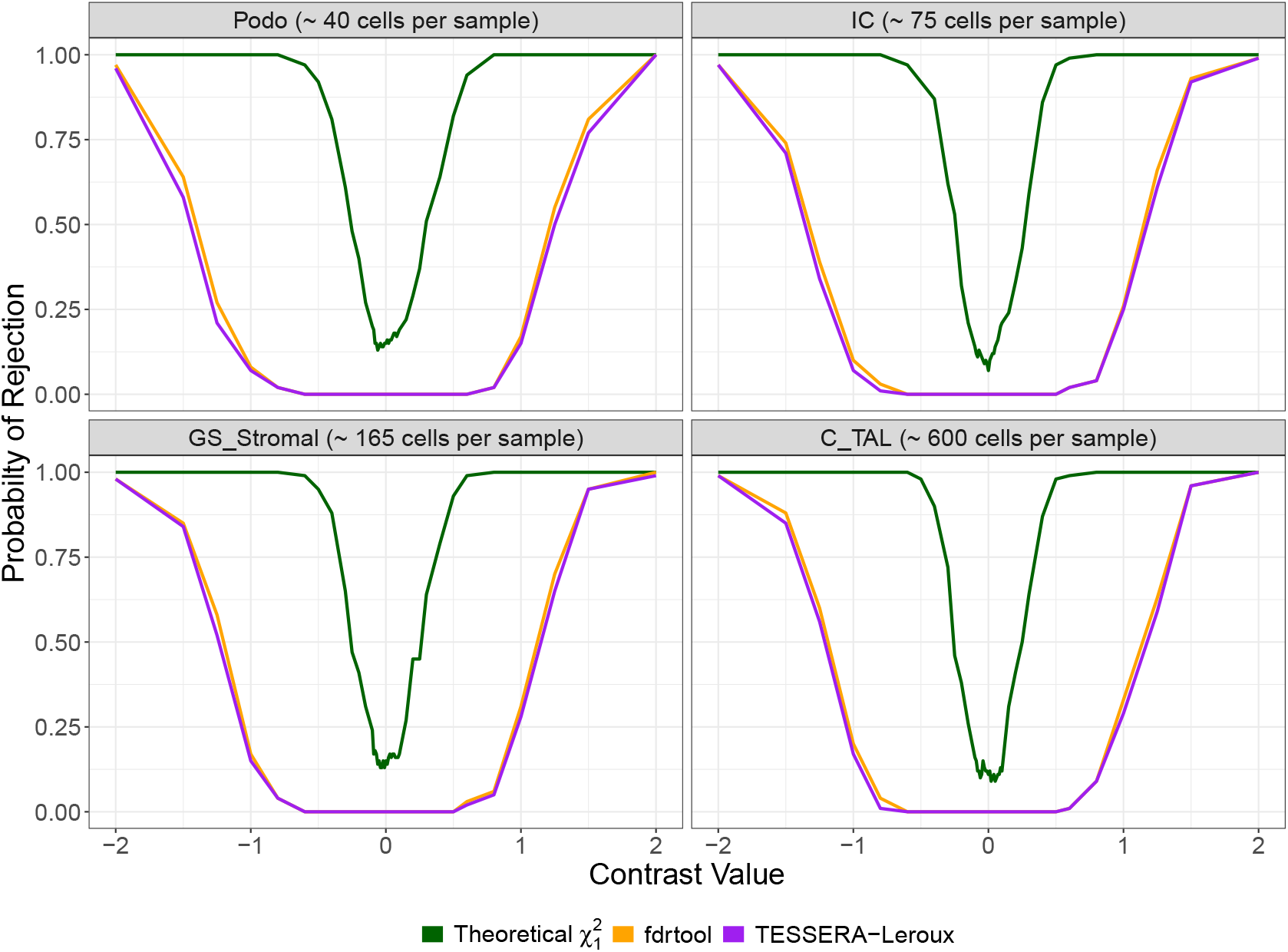
Power of empirical null distribution estimation procedures and theoretical 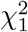 baseline in Poisson-Leroux-generated data. This figure examines power for detecting DE between conditions within a cell type for one gene. It presents plots of power (y-axis) as a function of the contrast value (x-axis), where power is estimated by the proportion of 100 independent simulation trials in which the null hypothesis of no DE is rejected at nominal Type I error rate of 0.05. At an effect size of 0, this value represents the Type I error rate. Synthetic data were generated using a Poisson-Leroux distribution, with generative parameters estimated from the *IGHG2* gene in a multi-sample real-world kidney dataset (Abedini et al., 2024). For each cell type shown, we modify two entries of the fixed effects vector ***β***, corresponding to the Control and the DKD conditions, to generate a contrast of specified size. We set the control coefficient to be the average of all coefficients and the DKD coefficient to be the control coefficient plus a given effect size. We then fit the `TESSERA` Poisson-Leroux model and evaluate the detectability of the contrasts by comparing two empirical null distribution estimation procedures, the one used by `TESSERA` (Section 2.4) and the default `fdrtool` (Strimmer, 2008) implementation, against a theoretical 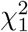 baseline.

**Figure S-25:**
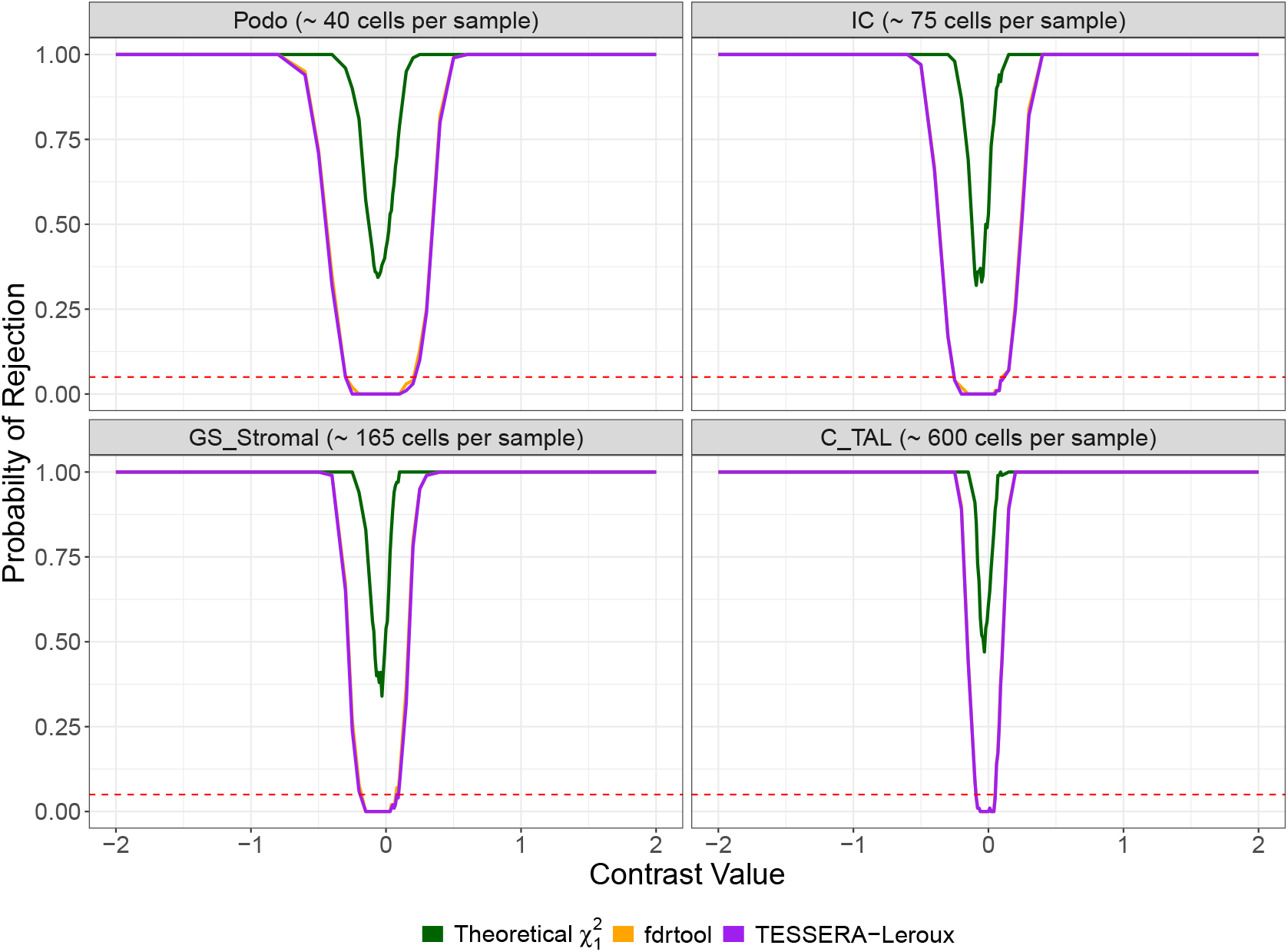
Power of empirical null distribution estimation procedures and theoretical 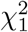 baseline in Poisson-spNNGP-generated data. This figure examines power for detecting DE between conditions within a cell type for one gene. It presents plots of power (y-axis) as a function of the contrast value (x-axis), where power is estimated by the proportion of 100 independent simulation trials in which the null hypothesis of no DE is rejected at nominal Type I error rate of 0.05. At an effect size of 0, this value represents the Type I error rate. Synthetic data were generated using a Poisson-spNNGP distribution, with generative parameters estimated from the *IGHG2* gene in a multi-sample real-world kidney dataset (Abedini et al., 2024). For each cell type shown, we modify two entries of the fixed effects vector ***β***, corresponding to the Control and the DKD conditions, to generate a contrast of specified size. We set the control coefficient to be the average of all coefficients and the DKD coefficient to be the control coefficient plus a given effect size. We then fit the `TESSERA` Poisson-Leroux model and evaluate the detectability of the contrasts by comparing two empirical null distribution estimation procedures, the one used by `TESSERA` (Section 2.4) and the default `fdrtool` (Strimmer, 2008) implementation, against a theoretical 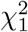 baseline.

#### S7.5. Real Data: Model Fit

**Figure S-26:**
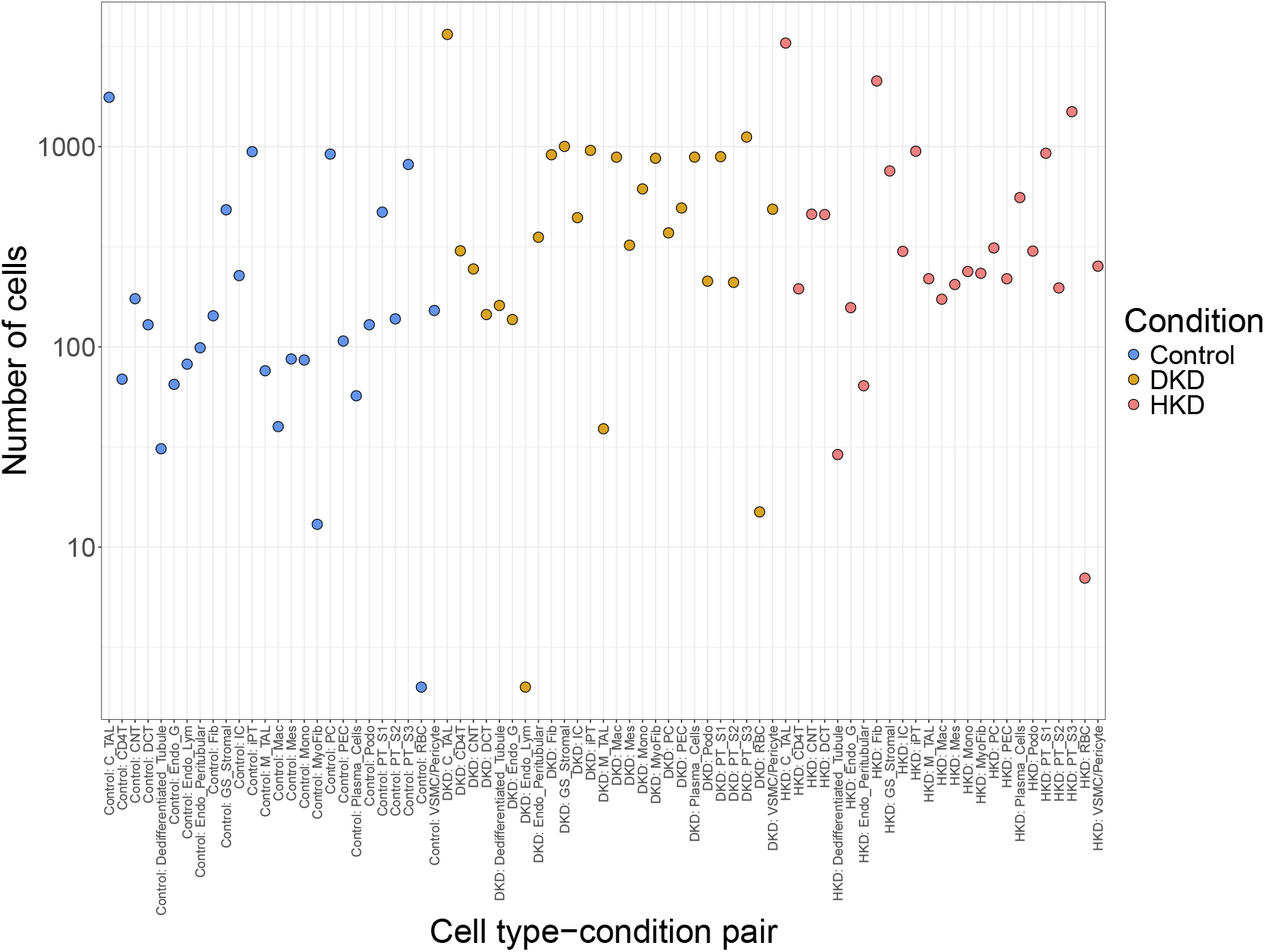
Number of expressed cells across cell type-condition pairs. We plot the total number of cells expressed, i.e., cells with non-zero expression measurements, for each cell type-condition pair (e.g., *C_TAL* cells in DKD samples) in the kidney dataset (Abedini et al., 2024). Each cell type-condition pair corresponds to a column in the design matrix (detailed in Section 4) and the number of expressed cells is the total number of non-zero entries in that column.

**Figure S-27:**
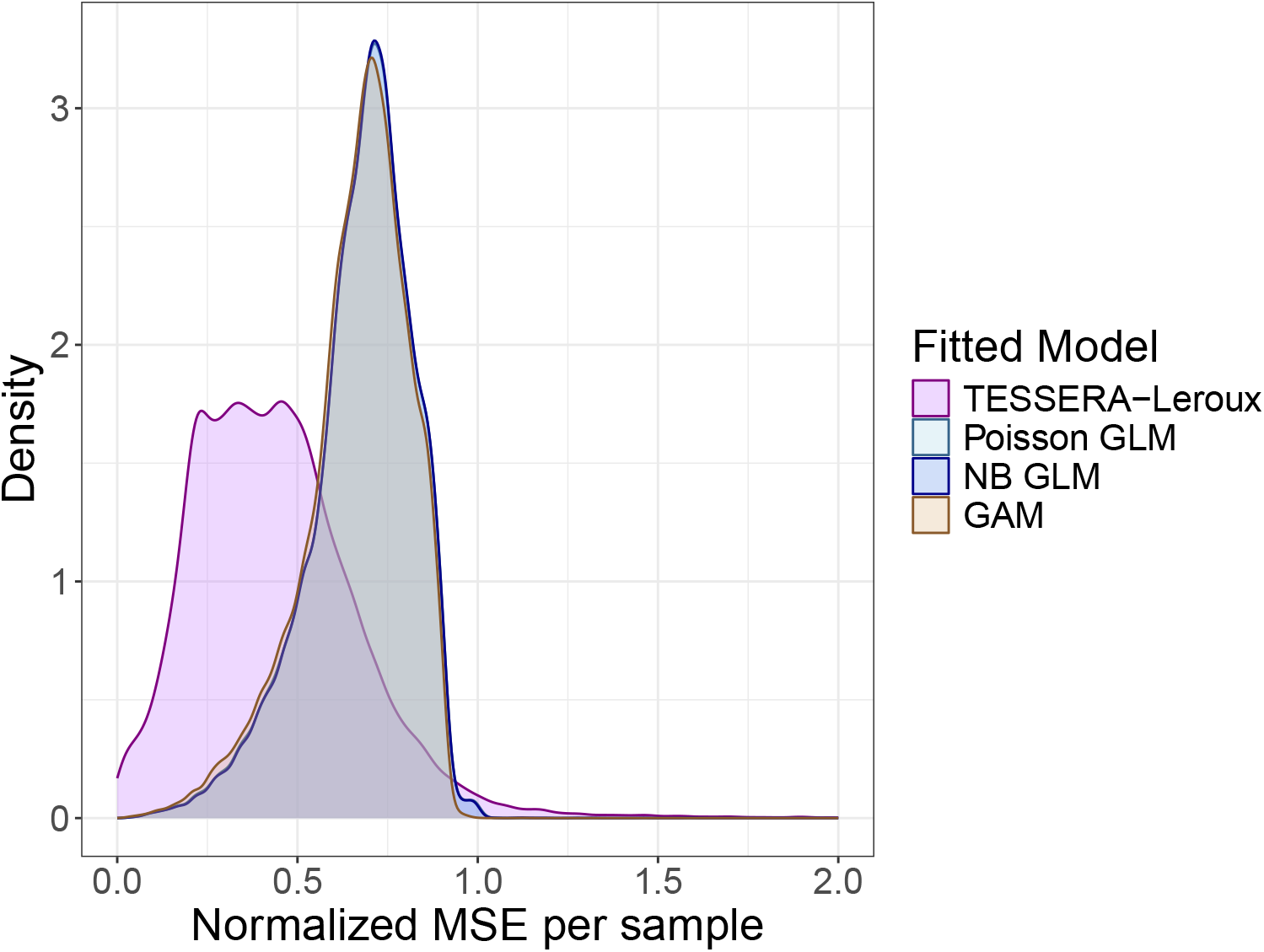
Normalized MSE per sample. For a multi-sample real-world kidney dataset (Abedini et al., 2024), we compare the distribution of the normalized mean squared error (MSE) for `TESSERA` (fitted with the Poisson-Leroux model) alongside a Poisson GLM, an NB GLM, and a GAM. For each sample-gene pair, the MSE is normalized by the mean of the squared counts: 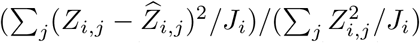. The `TESSERA` Poisson-Leroux model generally has smaller residuals relative to the other methods.

**Figure S-28:**
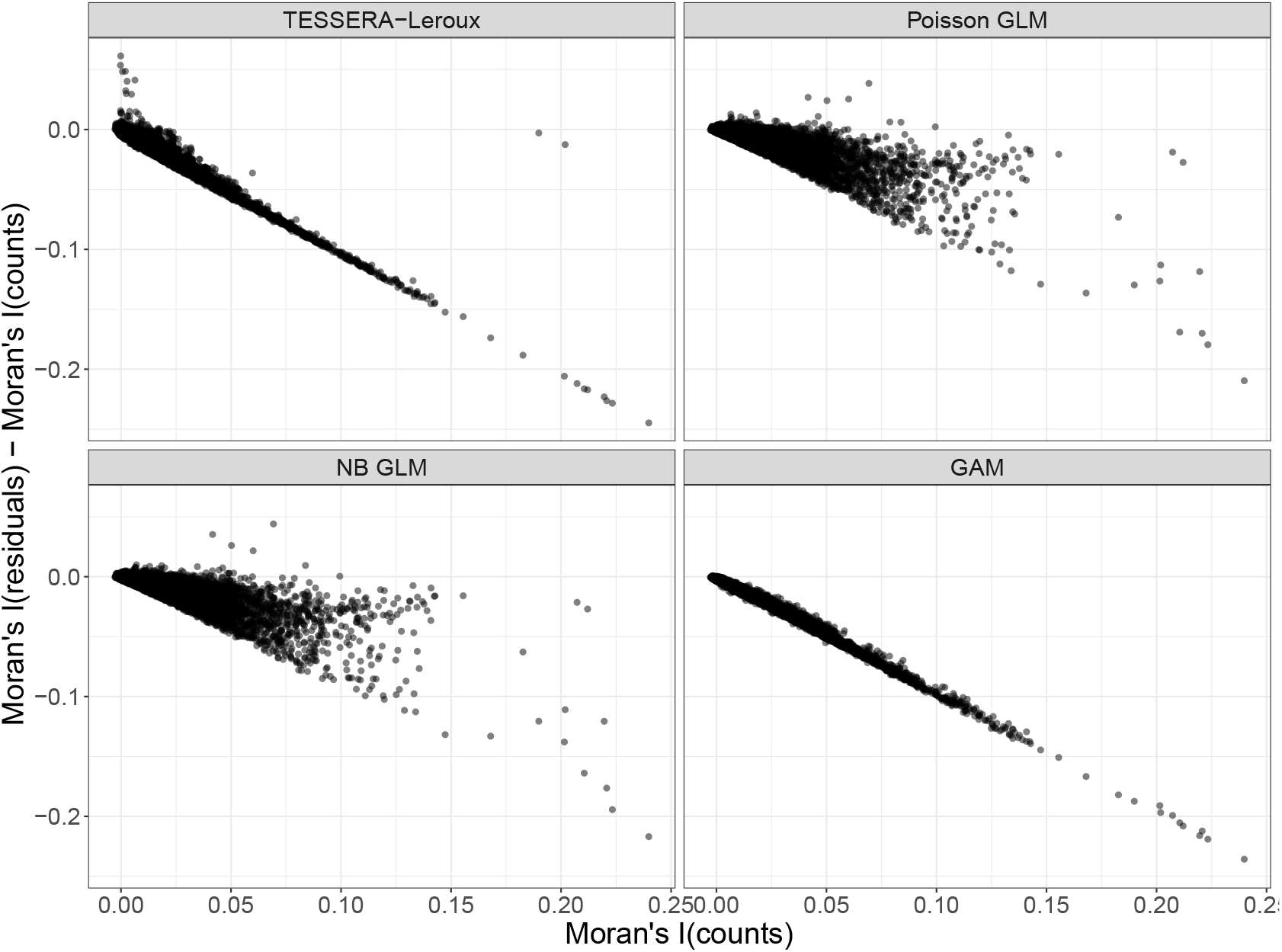
Reduction in Moran’s *I* of expression measurements. For a multi-sample real-world kidney dataset (Abedini et al., 2024), we assess the reduction in autocorrelation resulting from fitting a particular model by computing, for each gene-sample pair, the difference between the Moran’s *I* of the raw counts and that of the residuals. We compare `TESSERA` (fitted with a Poisson-Leroux model) against a Poisson GLM, an NB GLM, and a GAM. Plots of the differences in Moran’s *I* against the Moran’s *I* of the raw counts indicate that, as expected, the larger the raw count spatial dependencies, the larger the reduction in spatial dependencies.

**Figure S-29:**
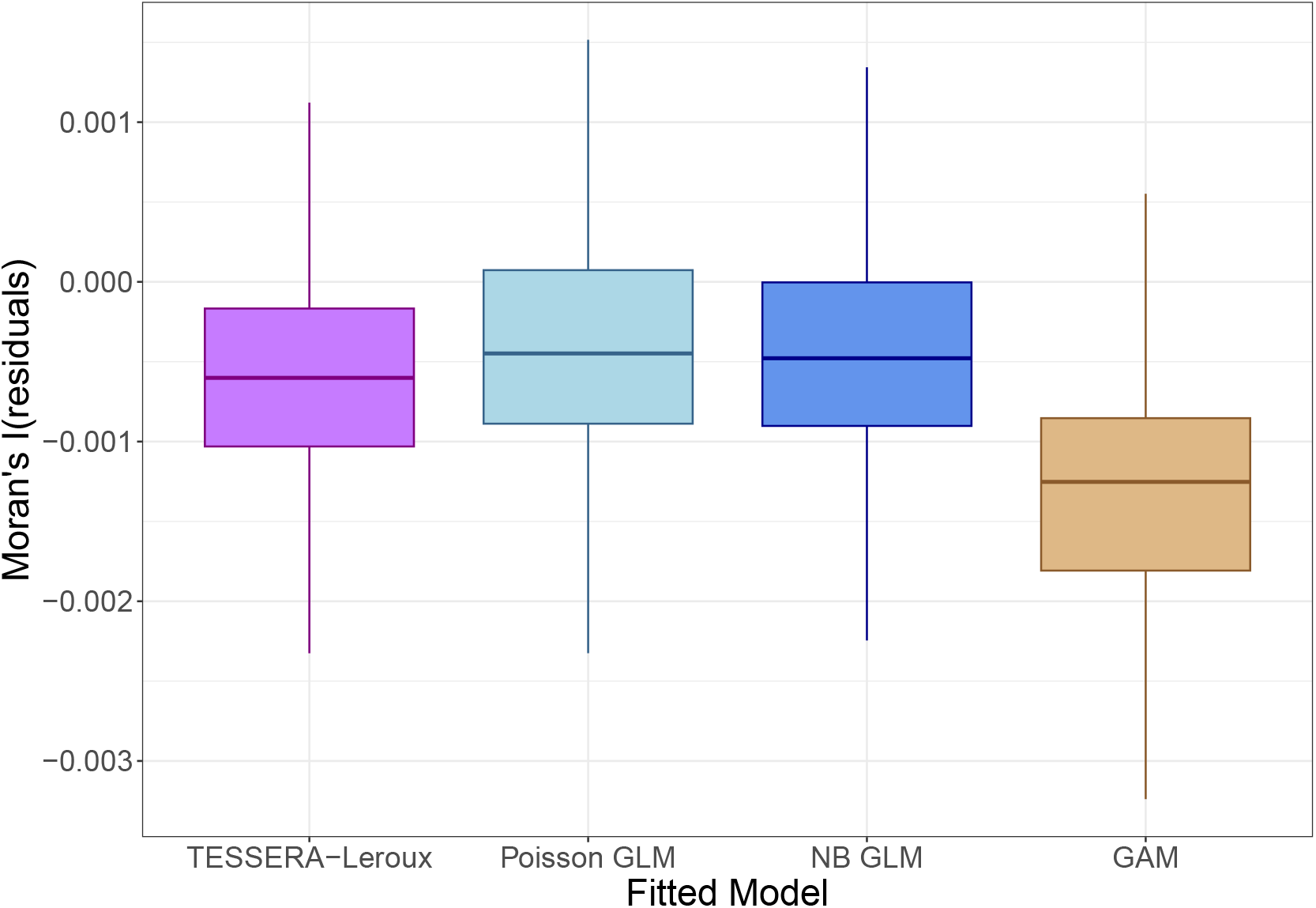
Moran’s *I* of residuals for spatially independent synthetic measurements. For each of 100 independent simulation trials, we generate a synthetic dataset based on the kidney dataset from Abedini et al. (2024) by sampling from a Poisson-Leroux distribution. We use the scale parameters *τ* ^2^ derived from fitting the model to the real data and set the spatial correlation parameter *γ* to zero, yielding a distribution with independent measurements. We then fit the `TESSERA` Poisson-Leroux model as well as a Poisson GLM, an NB GLM, and a GAM to the data. No significant spatial autocorrelation is exhibited, with all methods having Moran’s *I* ≈ 0.

**Figure S-30:**
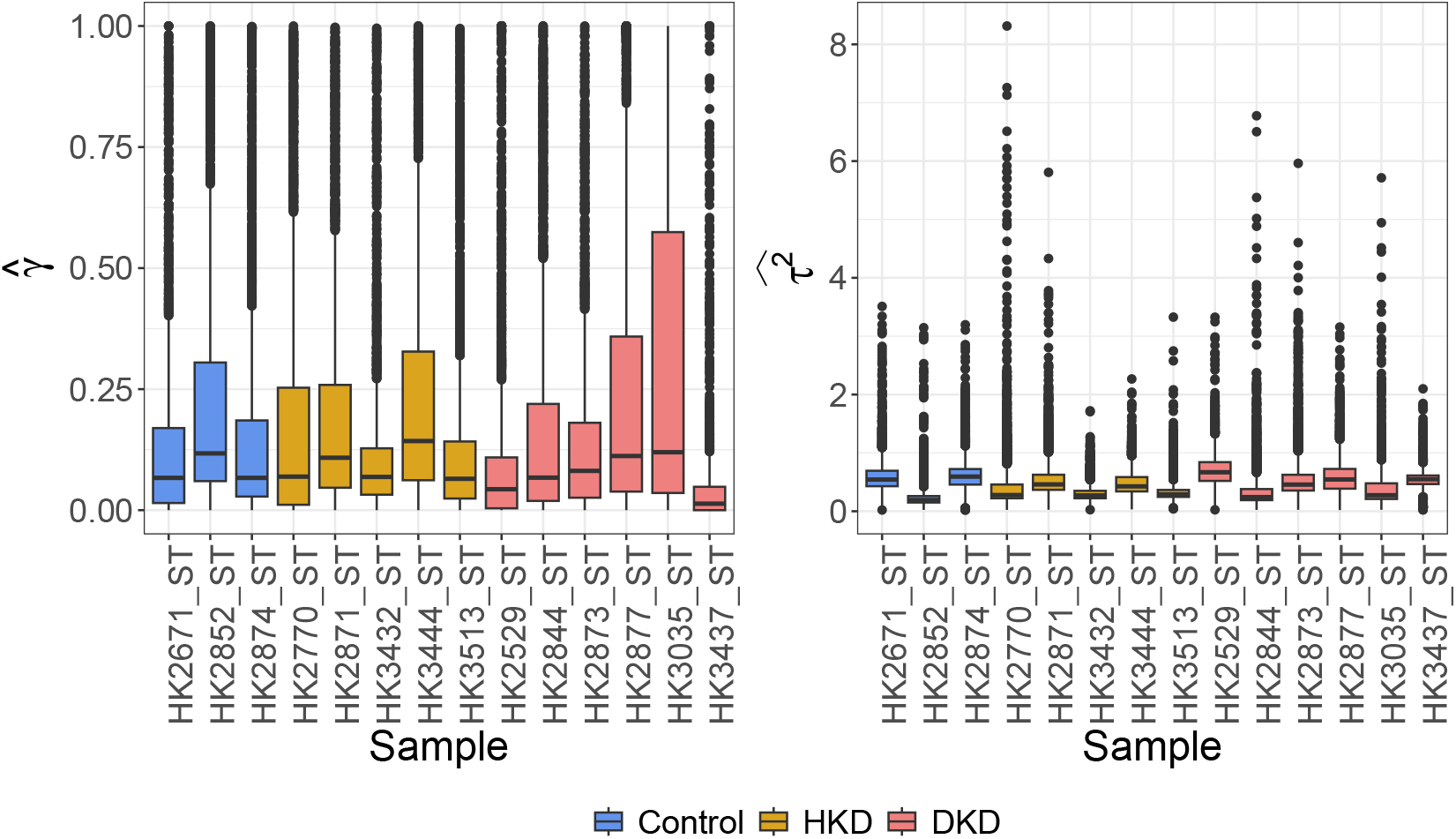
Estimated spatial covariance parameters for TESSERA fit with the Poisson-Leroux model. For a multi-sample real-world kidney dataset (Abedini et al., 2024), we present the gene- and sample-specific estimated spatial covariance parameters *γ* and *τ* ^2^ in boxplots stratified by sample. The boxplots reveal substantial inter-sample heterogeneity, indicating that the magnitude and structure of spatial dependence vary across individual samples. This variation underscores the necessity of a multi-sample method, while the predominantly non-zero estimates 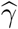 and 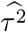 justify the inclusion of spatial random effects.

#### S7.6. Real Data: Differential Expression Between Cell Types

**Figure S-31:**
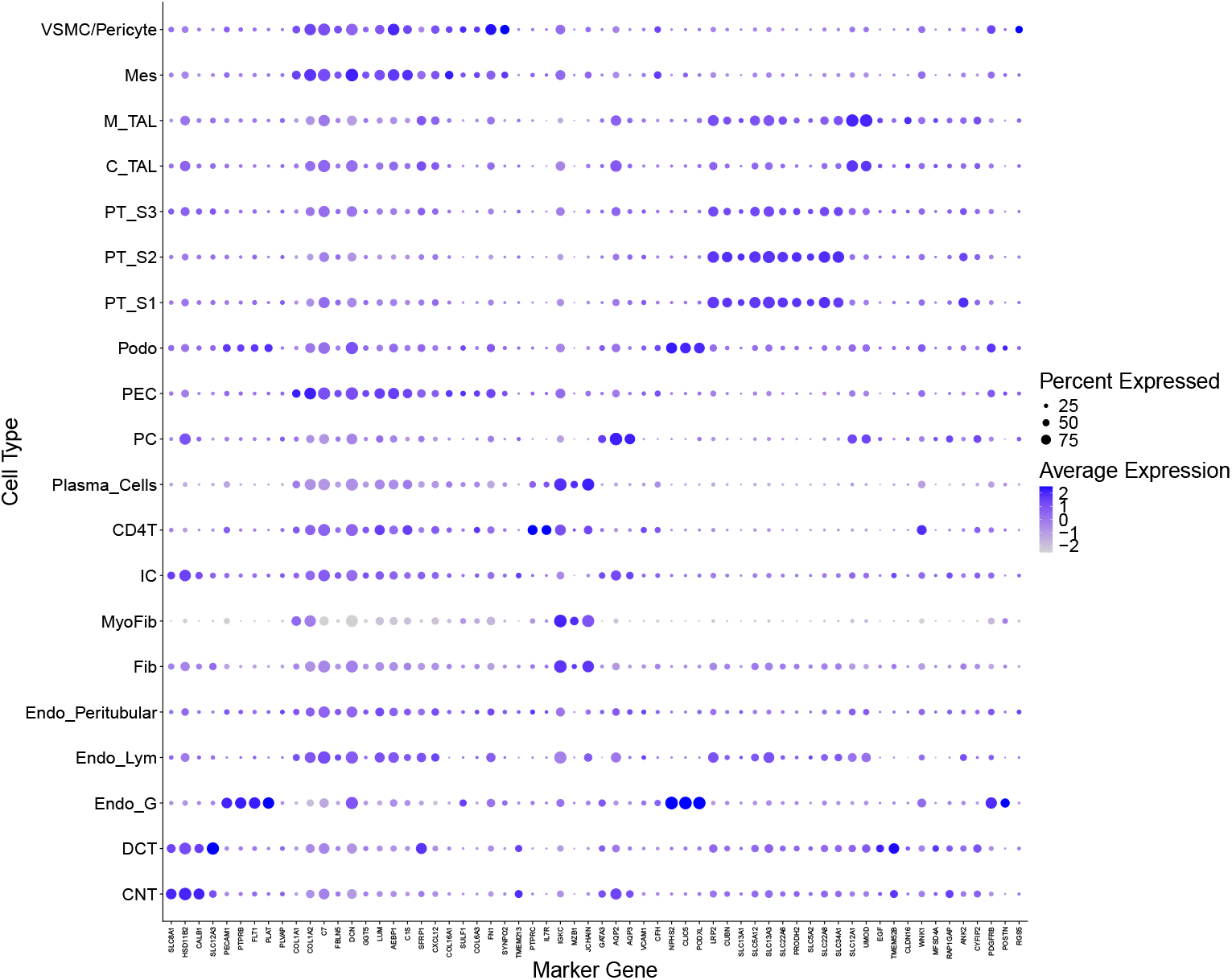
Marker gene expression for each cell type. We visualize the kidney marker genes outlined in Lake et al. (2023, Supp. Table 5) mapped to the same cell types present in the kidney dataset from Abedini et al. (2024). For the marker genes present in the dataset, we plot the percentage of cells expressed and the average expression, scaled to unit variance and zero mean across cell types following library-size normalization and log-transformation.

**Figure S-32:**
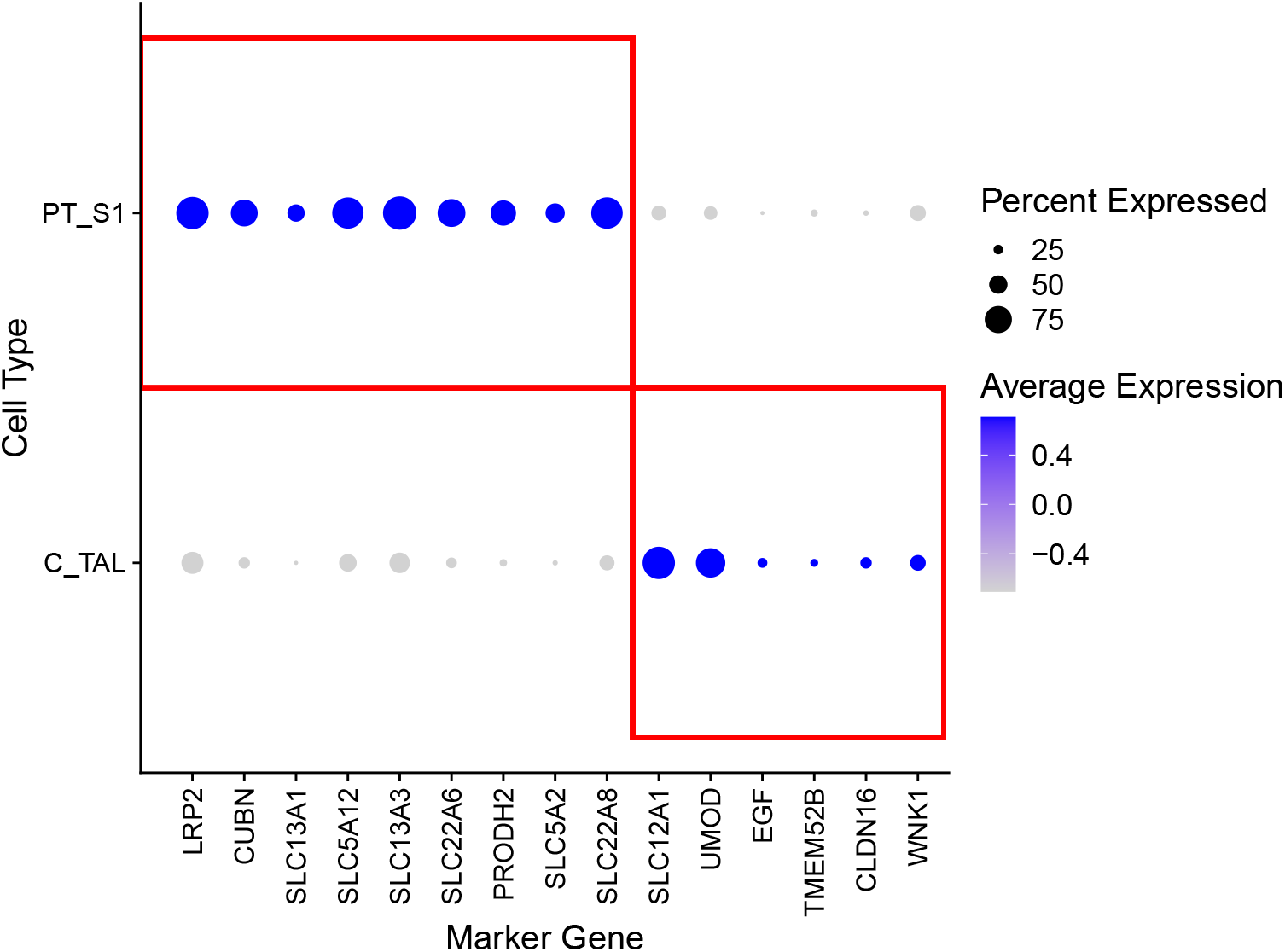
Marker gene expression for *PT_S1* and *C_TAL* cell types. We plot the average expression of marker genes for proximal tubule segment 1 (*PT_S1*) and cortical thick ascending limb (*C_TAL*) cell types in the kidney dataset (Abedini et al., 2024). These genes were selected based on established markers from Lake et al. (2023, Supp. Table 5) that were also present in the dataset. The red boxes indicate the specific marker genes associated with each cell type. We see that the marker genes serve as effective differentiators between the two cell populations.

**Figure S-33:**
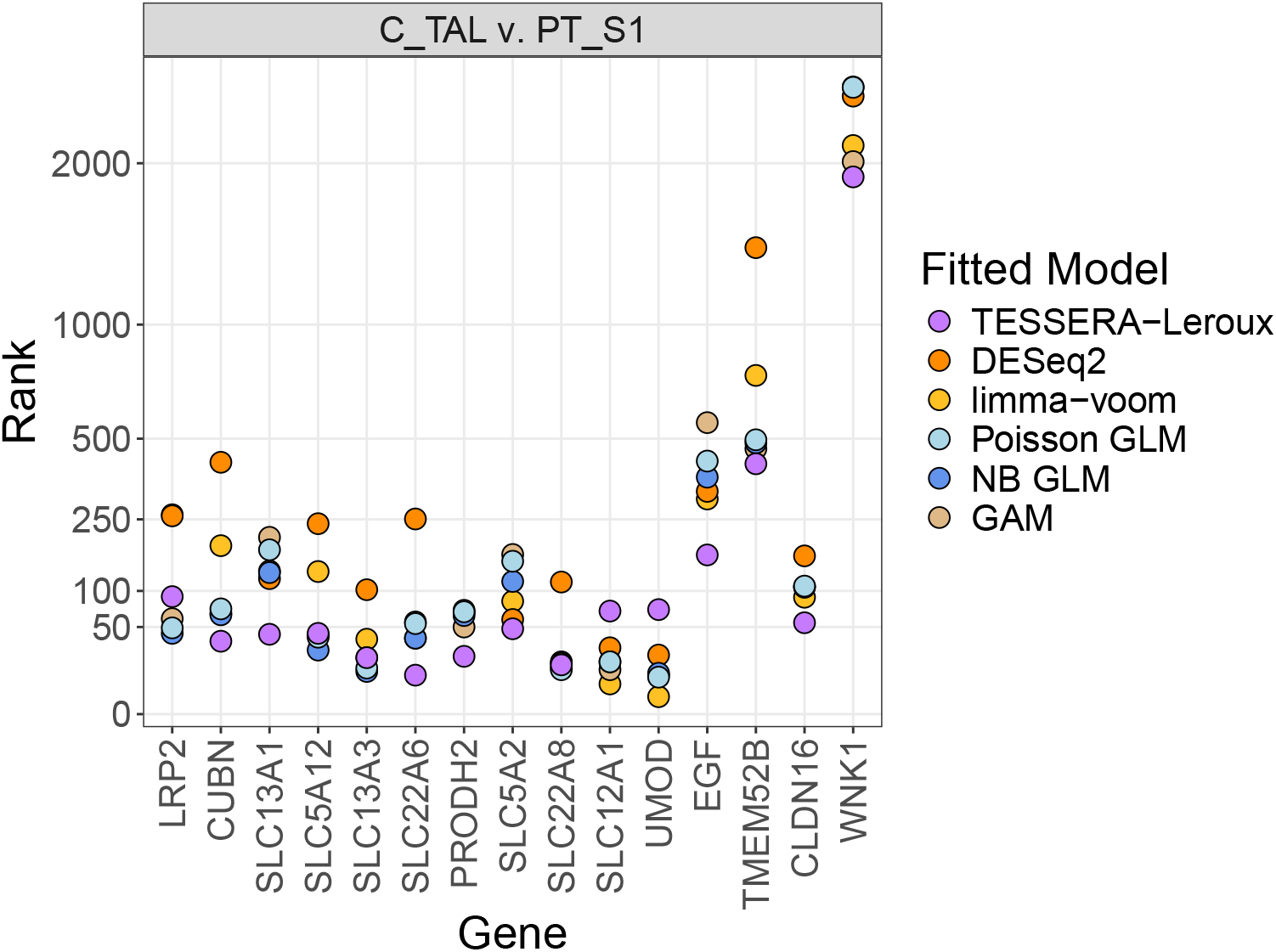
Evaluation of marker gene ranks for DE between cell types. `TESSERA` (fitted with a Poisson-Leroux model), a Poisson GLM, an NB GLM, a GAM, and pseudobulk methods (`DESeq2` and `limma-voom`) are compared in terms of their ability to identify DE genes between the *PT_S1* and *C_TAL* cell types, based on established kidney marker genes for *PT_S1* and *C_TAL* from Lake et al. (2023, Supp. Table 5), which are mapped to the same cell types present in the kidney dataset from Abedini et al. (2024). Genes are ranked according to their test statistics and the plot depicts these rankings specifically for the identified marker genes. A lower numerical rank indicates a larger test statistic, providing stronger evidence of differential expression.

**Figure S-34:**
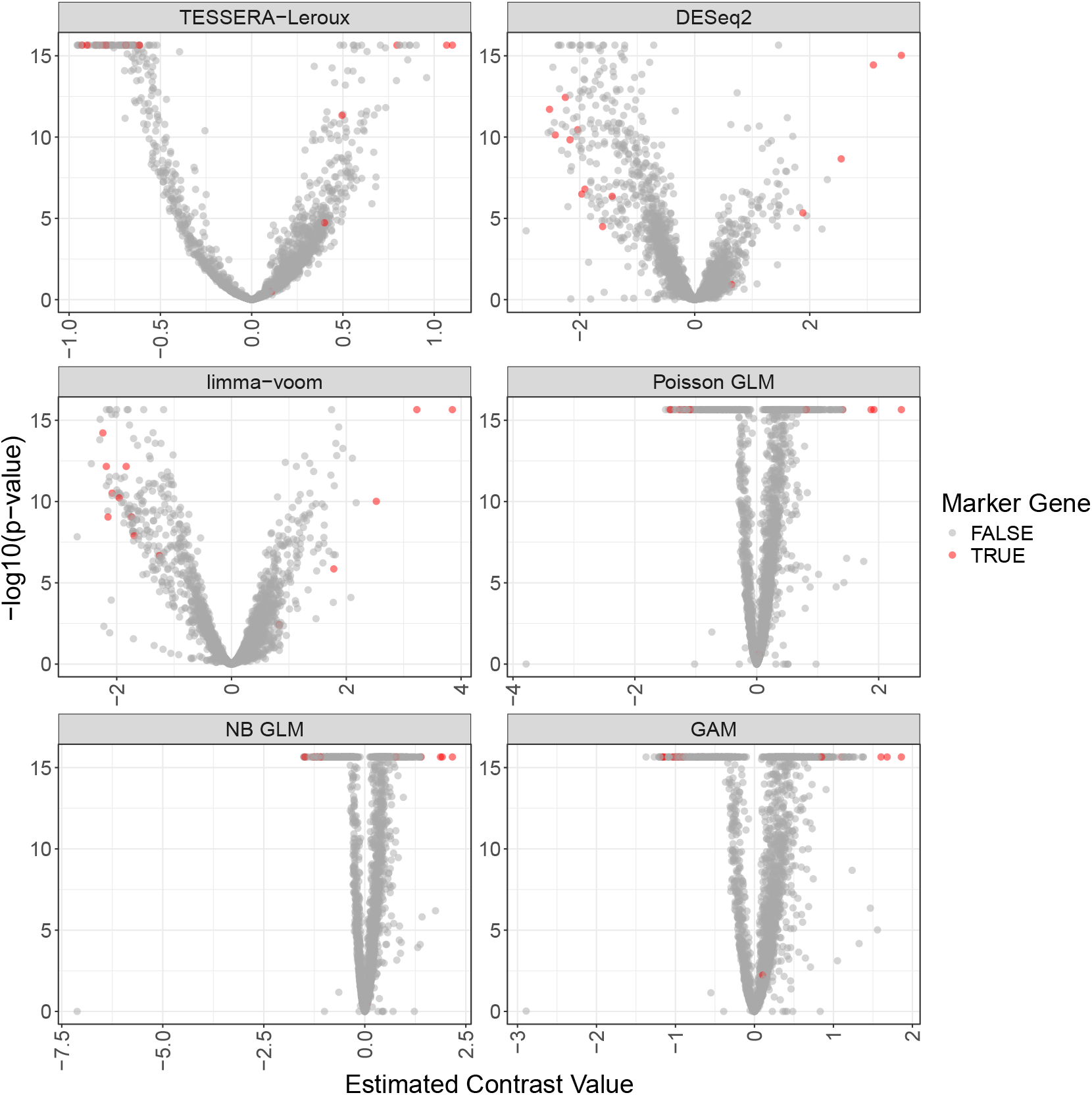
Volcano plot for DE between cell types. `TESSERA` (fitted with a Poisson-Leroux model), a Poisson GLM, an NB GLM, a GAM, and pseudobulk methods (`DESeq2` and `limma-voom`) are compared in terms of their ability to identify DE genes between the *PT_S1* and *C_TAL* cell types using volcano plots of −log_10_ *p*-values against estimated contrast values. Across all methods, established marker genes exhibit larger contrast magnitudes and lower *p*-values, as expected.

**Figure S-35:**
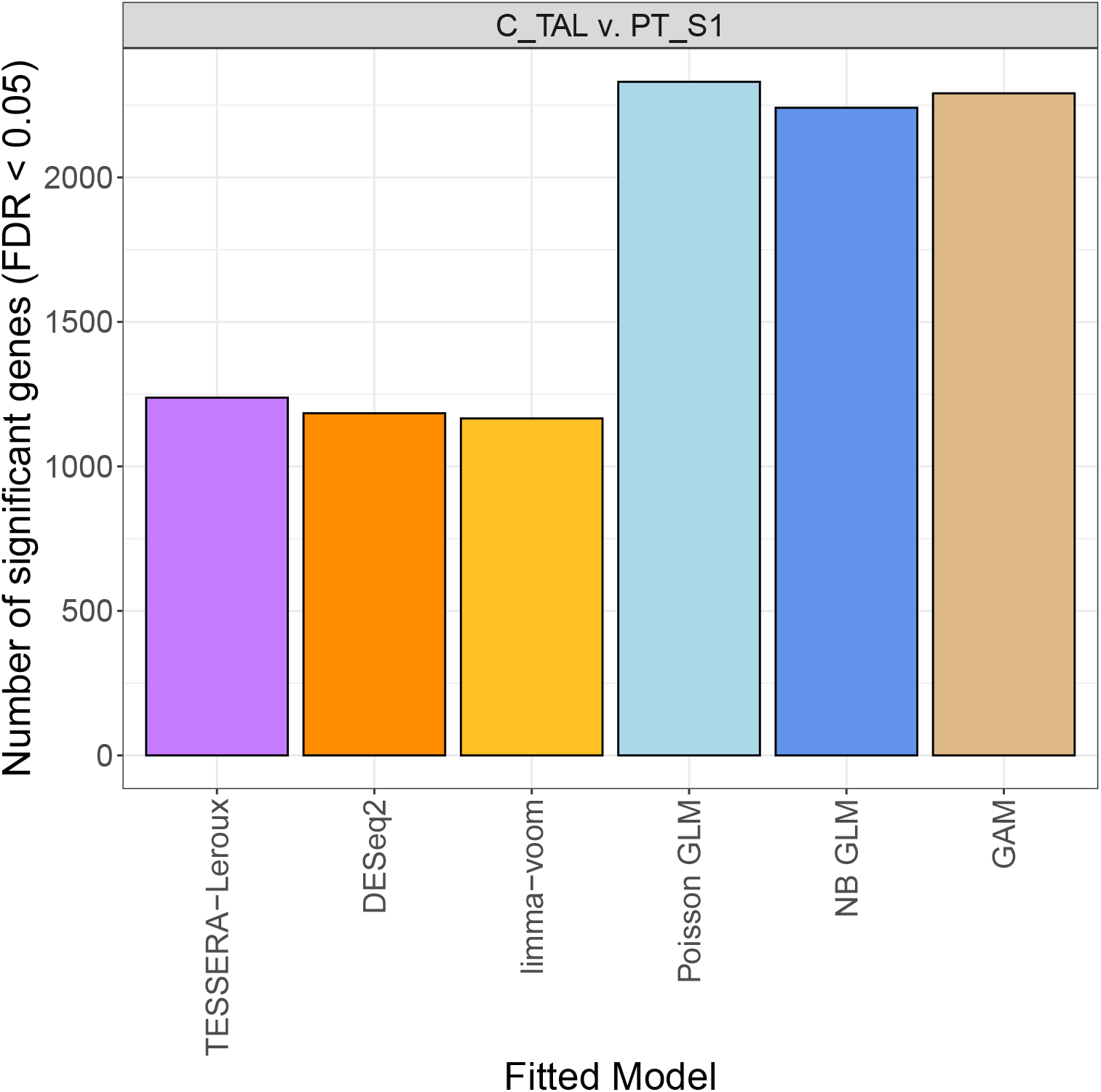
Number of genes found to be significantly DE between cell types. `TESSERA` (fitted with a Poisson-Leroux model), a Poisson GLM, an NB GLM, a GAM, and pseudobulk methods (`DESeq2` and `limma-voom`) are compared in terms of their ability to identify DE genes between the *PT_S1* and *C_TAL* cell types. We plot the total number of genes identified as differentially expressed by each method at a nominal FDR level of 0.05 using Benjamini-Hochberg adjusted *p*-values.

**Figure S-36:**
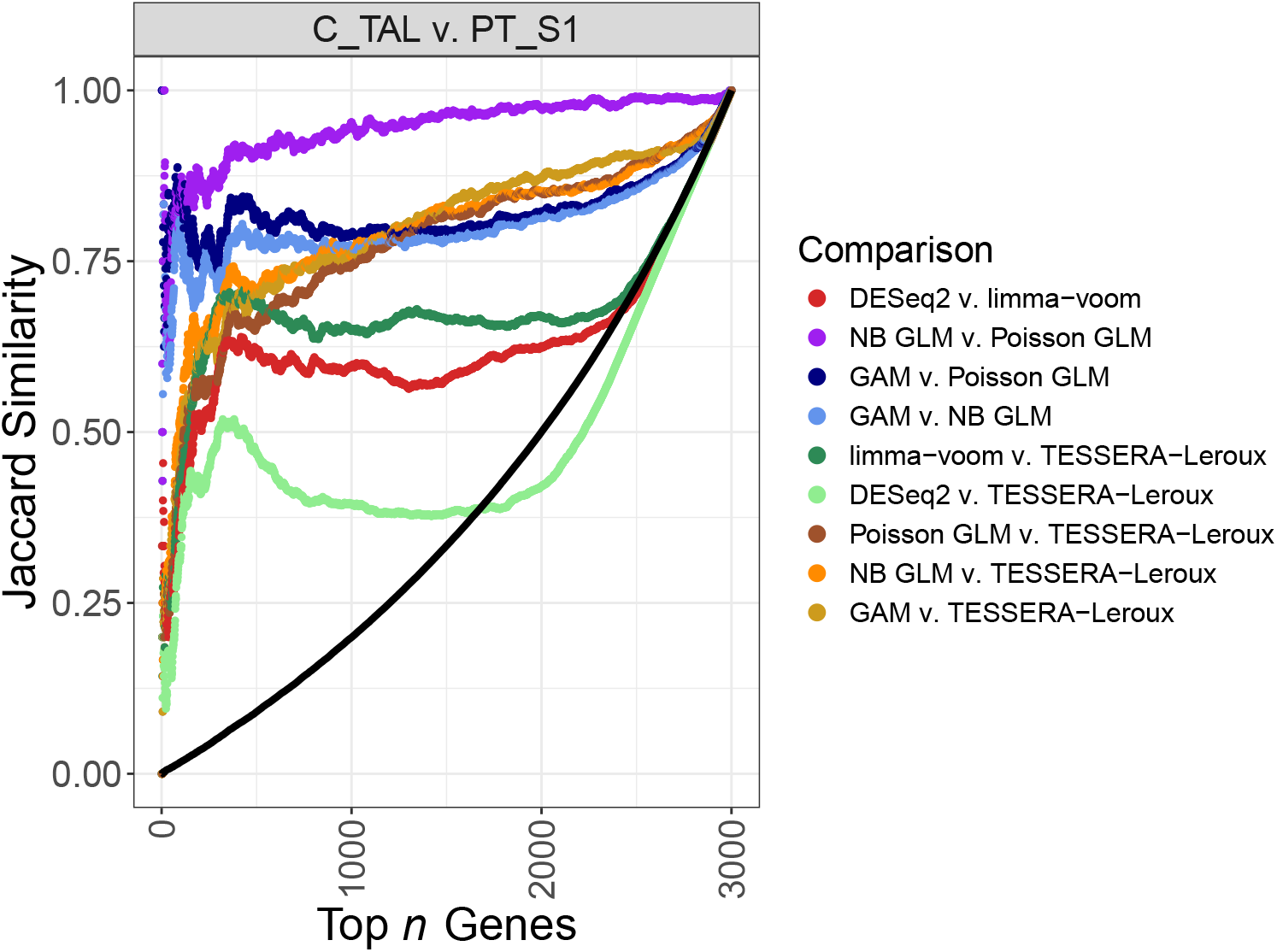
Jaccard similarity analysis of gene rankings for DE between cell types. `TESSERA` (fitted with a Poisson-Leroux model), a Poisson GLM, an NB GLM, a GAM, and pseudobulk methods (`DESeq2` and `limma-voom`) are compared in terms of their ability to identify DE genes between the *PT_S1* and *C_TAL* cell types. Genes are ranked according to their test statistics and rankings between pairs of methods are compared with the Jaccard similarity (the size of the intersection divided by the size of the union) for each possible set of top *n* genes, where *n* ranges from 1 to the total number of 3000 genes. The plot illustrates pairwise comparisons of `TESSERA` (fitted with a Poisson-Leroux model) against all other methods, the two pseudobulk methods against each other, and all combinations among the single-cell level non-spatial methods (Poisson GLM, NB GLM, and GAM). The black curve represents the null baseline, derived via simulation by calculating the average Jaccard similarity between two randomly permuted gene rankings across 100 trials.

#### S7.7. Real Data: Differential Expression Between Conditions, Within Cell Types

**Figure S-37:**
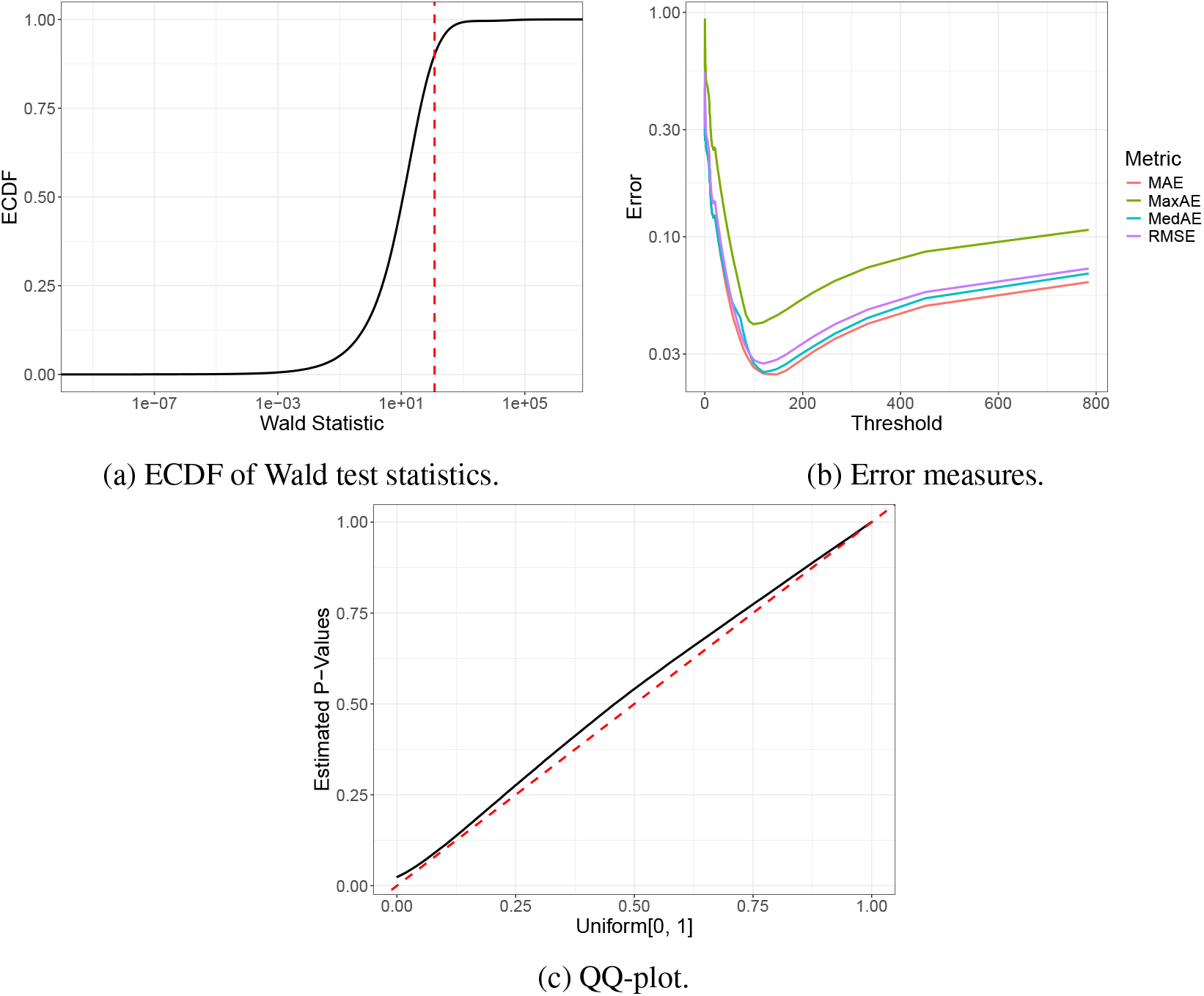
Behavior of TESSERA Wald statistics. For the kidney dataset (Abedini et al., 2024), we apply `TESSERA` fit with a Poisson-Leroux model to identify genes that are differentially expressed between conditions within the proximal tubule segment 1 (*PT_S1*) cell type. We evaluate the Wald test procedure described in Section 2.4 using several diagnostic plots. Figure S-37a: The empirical cumulative distribution function (ECDF) of the Wald test statistics shows that most statistics remain small, with a rapid increase occurring near the threshold indicated by the red dashed vertical line. Figure S-37b: Our procedure involves selecting a threshold below which statistics are considered to follow the “null” distribution. We measure the error between the resulting *p*-values and a uniform distribution on [0, 1] as a function of this threshold. While we formally select the threshold by minimizing the MSE, the sharp trough observed across different error measures identifies a natural choice for the threshold; we plot the RMSE alongside these metrics to maintain a consistent scale for visual comparison. Figure S-37c: A quantile-quantile plot (QQ-plot) of the “null” *p*-values against the Uniform [0, 1] distribution demonstrates a high-quality fit, supported by an *L*^2^-error of 3.4 *×* 10^−4^.

**Figure S-38:**
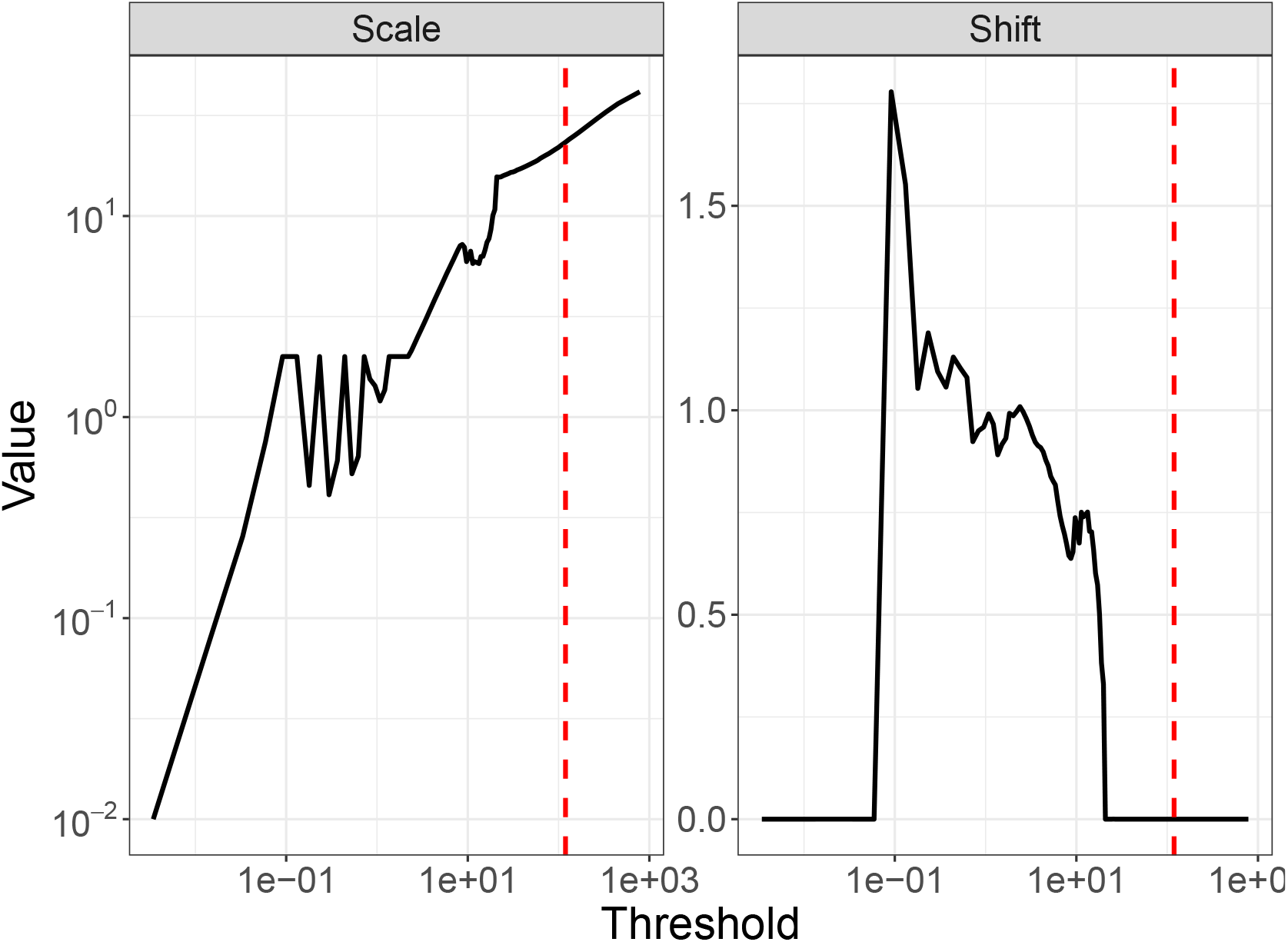
Estimation of null distribution parameters for TESSERA Wald test statistics. For the kidney dataset (Abedini et al., 2024), we apply `TESSERA` fit with a Poisson-Leroux model to identify genes that are differentially expressed between conditions within the proximal tubule segment 1 (*PT_S1*) cell type. We evaluate the Wald test procedure described in Section 2.4 using diagnostic plots of the null distribution parameters. According to our model, the test statistics follow a scaled non-central 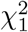 distribution under the null hypothesis. We display the estimated parameters of this distribution as a function of the threshold for determining the statistics that come from the “null” distribution. The threshold, selected to minimize the *L*^2^-error (or MSE) relative to a Uniform [0, 1] distribution, is indicated by a red dashed vertical line in both panels. Notably, this threshold occurs just prior to the point where the scale and shift parameters reach a plateau.

**Figure S-39:**
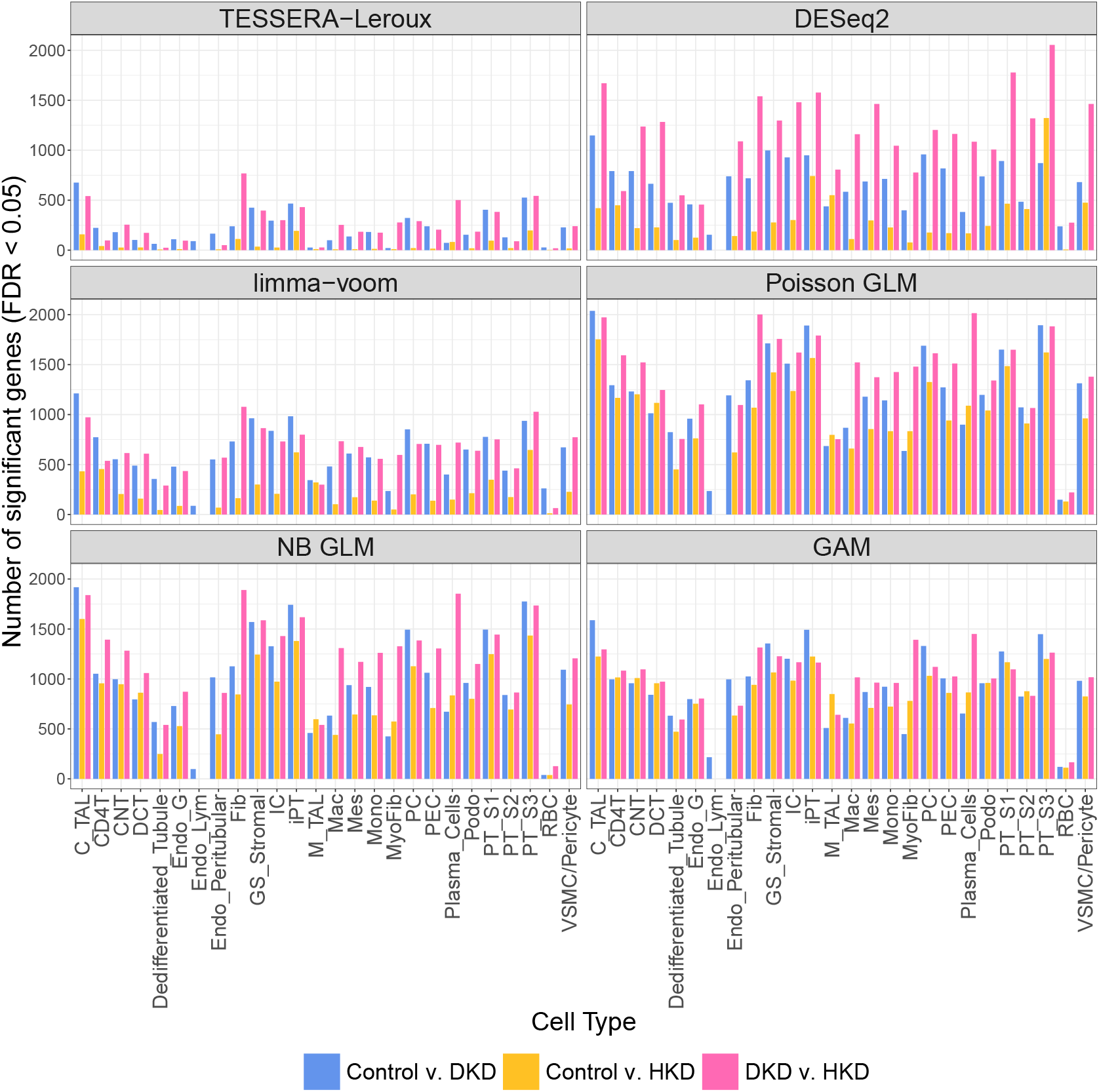
Number of differentially expressed genes (faceted by method). For the kidney dataset (Abedini et al., 2024), we apply several methods to identify genes that are differentially expressed between pairs of conditions within the proximal tubule segment 1 (*PT_S1*) cell type. Significance is determined at a nominal FDR level of 0.05 using Benjamini-Hochberg adjusted *p*-values. We compare `TESSERA` (fitted with a Poisson-Leroux model) against a Poisson GLM, an NB GLM, a GAM, and pseudobulk methods (`DESeq2` and `limma-voom`). The individual facets of the plot represent the specific methods used to identify these differentially expressed genes. Within each facet, the bars represent the total number of genes identified as differentially expressed for each condition pair across the various cell types.

**Figure S-40:**
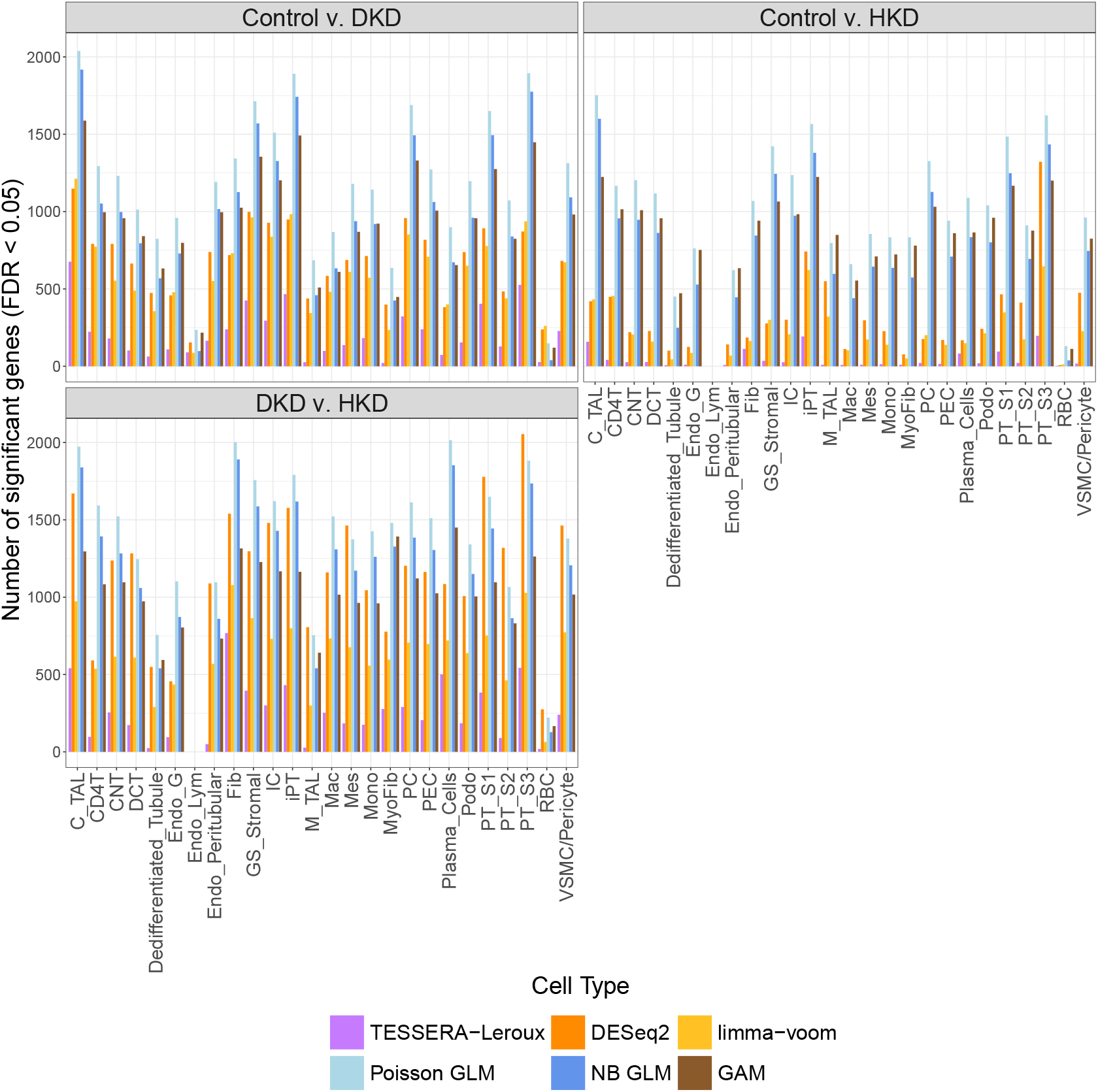
Number of differentially expressed genes (faceted by condition pair). For the kidney dataset (Abedini et al., 2024), we apply several methods to identify genes that are differentially expressed between pairs of conditions within the proximal tubule segment 1 (*PT_S1*) cell type. Significance is determined at a nominal FDR level of 0.05 using Benjamini-Hochberg adjusted *p*-values. We compare `TESSERA` (fitted with a Poisson-Leroux model) against a Poisson GLM, an NB GLM, a GAM, and pseudobulk methods (`DESeq2` and `limma-voom`). The individual facets of the plot represent the specific condition pairs being compared. Within each facet, the bars represent the total number of genes identified as differentially expressed by each respective method.

**Figure S-41:**
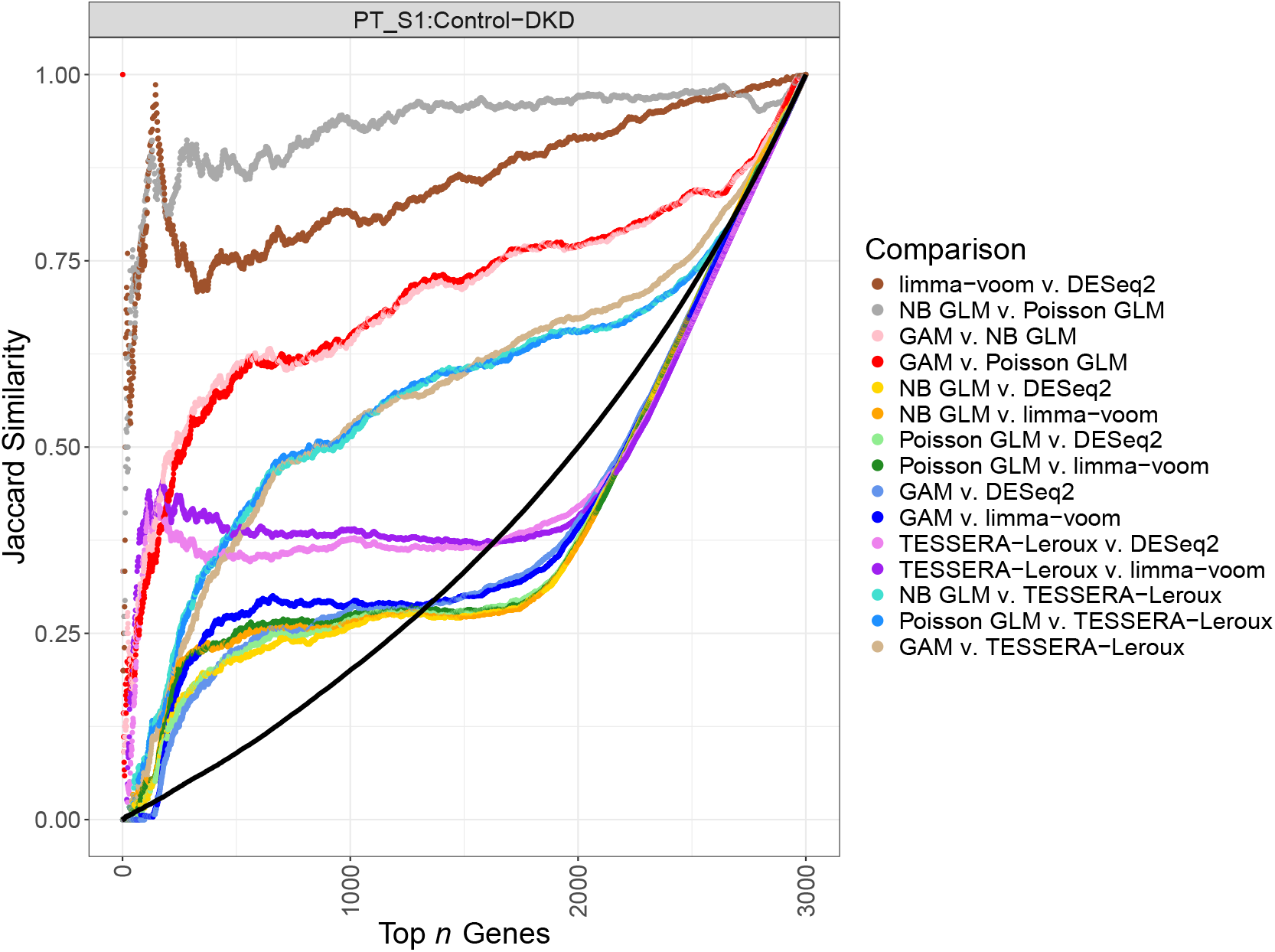
Jaccard similarity analysis of gene rankings. For the kidney dataset (Abedini et al., 2024), we apply several methods to identify genes that are differentially expressed between the Control and DKD conditions within the proximal tubule segment 1 (*PT_S1*) cell type. Genes are ranked by their test statistics calculated from `TESSERA` (fitted with a Poisson-Leroux model), a Poisson GLM, an NB GLM, a GAM, and pseudobulk methods (`DESeq2` and `limma-voom`). We compare the rankings between pairs of methods with the Jaccard similarity (the size of the intersection divided by the size of the union) for each possible set of top *n* genes, where *n* ranges from 1 to the total 3, 000 genes. The plot illustrates pairwise comparisons of `TESSERA` (fitted with a Poisson-Leroux model) against all other methods, alongside comparisons between each pair of non-spatial methods, which include both pseudobulk approaches (`limma-voom`, `DESeq2`) and single-cell level models (Poisson GLM, NB GLM, and GAM). The black curve represents the null baseline, derived via simulation by calculating the average Jaccard similarity between two randomly permuted gene rankings across 100 trials.

**Figure S-42:**
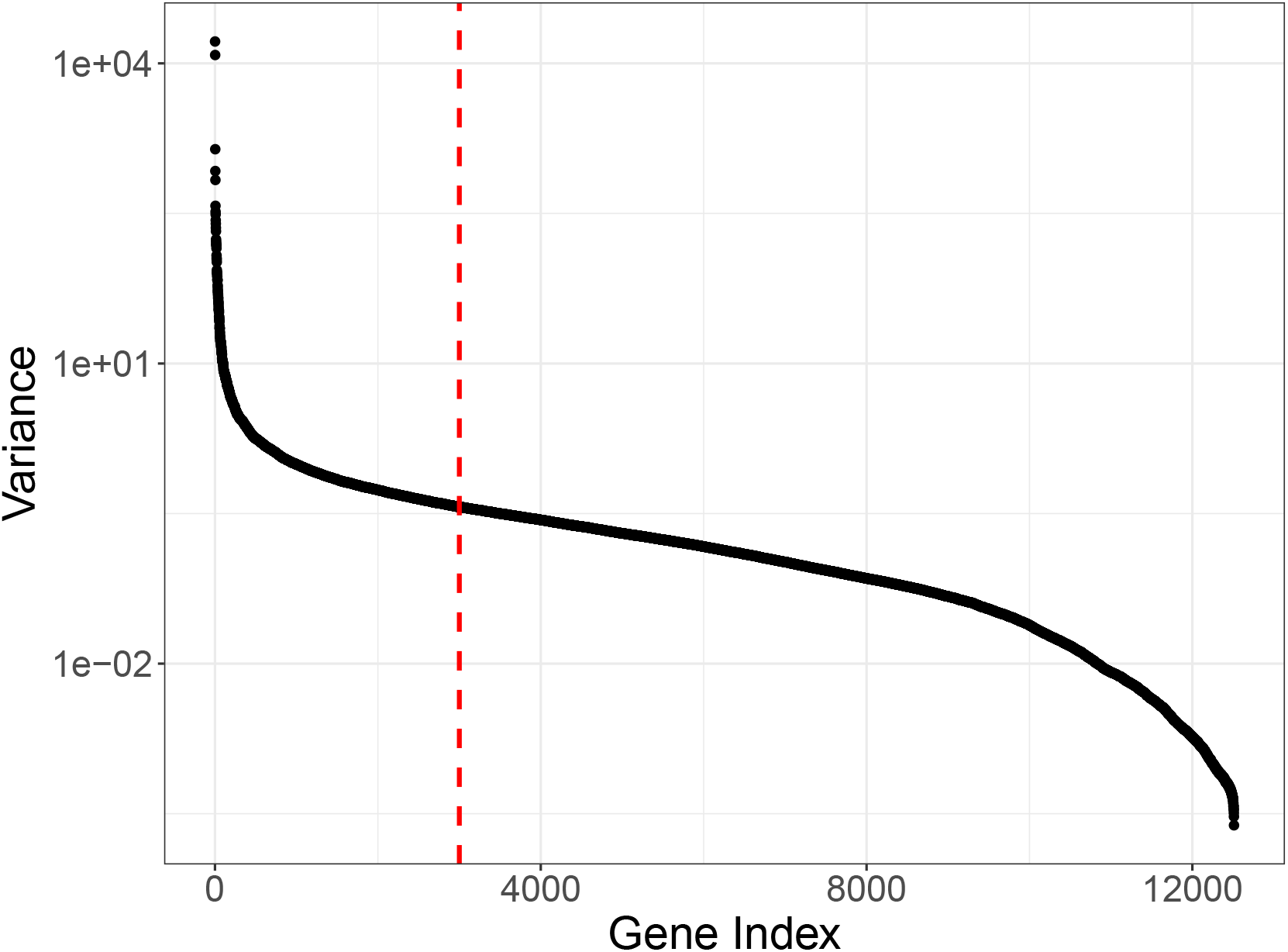
Gene selection based on raw count variance. We plot the sorted variance of raw counts (computed for each gene across all cells) in the kidney dataset (Abedini et al., 2024). The 3, 000 most highly variable genes were selected for downstream analysis as they capture the majority of the variation; beyond this threshold (indicated by the dashed vertical line), the variance profile flattens significantly, indicating a transition toward genes which we also found to have low expression.

